## Supplementary_Table_1 for "High-Throughput Functional Evaluation of *BRCA2* Variants of Unknown Significance"

Supplementary Table 1: Information of 107 initial variants and empty vector.

| # | Variant | HGVS mutation | Barcode | IARC classification | ClinVar cllasification | Align-GVGD Classification From Human To Sea Urchin | Align-GVGD key domains | SPiCE splice abberation probability | SPiCE interpretation | Mutagenesis primer-forward | Mutagenesis primer-reverse |
|---|---|---|---|---|---|---|---|---|---|---|---|
| 1 | Wild-type | - | TCGTTTGTCT | Class 1 | Benign | Class C0 | - | - | - | - | - |
| 2 | Empty-vector | - | GTGTGGTGGG | Class 5 | Pathogenic | Non-sense | - | - | - | - | - |
| 3 | R18H | c.53G>A | ATCGTATTTC | Class 2 | Benign | Class C0 | PALB2 interaction domain | 0.027 | Outside SPiCE Interpretation | CATTTTTTGAAATTTTTAAGACACACTGCAACAAAGCAGATTTAGGACC | GGTCCTAAATCTGCTTTGTTGCAGTGTGTCTTAAAAATTTCAAAAAATG |
| 4 | K21R | c.62A>G | TCTGCGATTC | Unclassified | VUS | Class C0 | PALB2 interaction domain | 0.027 | Outside SPiCE Interpretation | TTTGAAATTTTTAAGACACGCTGCAACAGAGCAGATTTAGGACC | GGTCCTAAATCTGCTCTGTTGCAGCGTGTCTTAAAAATTTCAAA |
| 5 | W31C | c.93G>T | TCTTTATGGT | Unclassified | Conflicting | Class C65 | PALB2 interaction domain | 0.027 | Outside SPiCE Interpretation | TTAGGACCAATAAGTCTTAATTGTTTTGAAGAACTTTCTTCAGAAGC | GCTTCTGAAGAAAGTTCTTCAAAACAATTAAGACTTATTGGTCCTAA |
| 6 | P41L | c.122C>T | CCTTGTGCAT | Unclassified | Conflicting | Class C0 | - | 0.027 | Outside SPiCE Interpretation | CTTTCTTCAGAAGCTCCACTCTATAATTCTGAACCTGCA | TGCAGGTTCAGAATTATAGAGTGGAGCTTCTGAAGAAAG |
| 7 | Y42C | c.125A>G | GGGCGGTTGT | Class 1 | Benign | Class C0 | - | 0.027 | Outside SPiCE Interpretation | TTCAGAAGCTCCACCCTGTAATTCTGAACCTGCAG | CTGCAGGTTCAGAATTACAGGGTGGAGCTTCTGAA |
| 8 | N55S | c.164A>G | TTCCTTTACT | Unclassified | VUS | Class C0 | - | 0.027 | Outside SPiCE Interpretation | GCAGAAGAATCTGAACATAAAAACAGCAATTACGAACCAAACCTATTTAA | TTAAATAGGTTTGGTTCGTAATTGCTGTTTTTATGTTCAGATTCTTCTGC |
| 9 | P59A | c.175C>G | ATGGGACTTG | Unclassified | Conflicting | Class C0 | - | 0.027 | Outside SPiCE Interpretation | TGAACATAAAAACAACAATTACGAAGCAAACCTATTTAAAACTCCACAAAG | CTTTGTGGAGTTTTAAATAGGTTTGCTTCGTAATTGTTGTTTTTATGTTCA |
| 10 | N108S | c.323A>G | ATTCTGTTTC | Unclassified | VUS | Class C0 | - | 0.027 | Outside SPiCE Interpretation | ATTAGATAAATTCAAATTAGACTTAGGAAGGAGTGTTCCCAATAGTAGAC | GTCTACTATTGGGAACACTCCTTCCTAAGTCTAATTTGAATTTATCTAAT |
| 11 | Q147R | c.440A>G | CGCCTGACGT | Unclassified | Benign/Likely benign | Class C0 | - | 0.027 | Outside SPiCE Interpretation | TGAAAGTCCTGTTGTTCTACGATGTACACATGTAACACCAC | GTGGTGTTACATGTGTACATCGTAGAACAACAGGACTTTCA |
| 12 | D156G | c.467A>G | GCACTTTCTA | Unclassified | Conflicting | Class C0 | - | 0.027 | Outside SPiCE Interpretation | CATGTAACACCACAAAGAGGTAAGTCAGTGGTATGTGGG | CCCACATACCACTGACTTACCTCTTTGTGGTGTTACATG |
| 13 | V159E | c.476T>A | TGGCCGGTGC | Unclassified | VUS* | Class C0 | - | 0.007 | low | CCACAAAGAGATAAGTCAGAGGTATGTGGGAGTTTGTTT | AAACAAACTCCCACATACCTCTGACTTATCTCTTTGTGG |
| 14 | P168T | c.502C>A | GGTACCTCGT | Class 1 | Benign | Class C0 | - | 0.027 | Outside SPiCE Interpretation | TGGGAGTTTGTTTCATACAACAAAGTTTGTGAAGGGTCG | CGACCCTTCACAAACTTTGTTGTATGAAACAAACTCCCA |
| 15 | V208G | c.623T>G | TTTGCTCGTT | Unclassified | VUS | Class C15 | - | 0.027 | Outside SPiCE Interpretation | CCACCCTTAGTTCTACTGGGCTCATAGTCAGAAATGA | TCATTTCTGACTATGAGCCCAGTAGAACTAAGGGTGG |
| 16 | F266L | c.796T>C | TAGCCTCTGC | Unclassified | Conflicting | Class C0 | - | 0.023 | Outside SPiCE Interpretation | GAAGCTGCAAGTCATGGACTTGGAAAAACATCAGGGA | TCCCTGATGTTTTTCCAAGTCCATGACTTGCAGCTTC |
| 17 | R324T | c.971G>C | GTTGAGGTTG | Unclassified | VUS | Class C0 | - | 0.027 | Outside SPiCE Interpretation | TCTAAATGTAGAACAAAAAATCTACAAAAAGTAACAACTAGCAAGACTAGGAAAAA | TTTTTCCTAGTCTTGCTAGTTGTTACTTTTTGTAGATTTTTTGTTCTACATTTAGA |
| 18 | S326R | c.978C>A | TTGAATGTCC | Class 1 | Benign | Class C0 | - | 0.027 | Outside SPiCE Interpretation | CAAAAAATCTACAAAAAGTAAGAACTAGAAAGACTAGGAAAAAAATTTTCCATGAAG | CTTCATGGAAAATTTTTTTCCTAGTCTTTCTAGTTCTTACTTTTTGTAGATTTTTTG |
| 19 | S445Y | c.1334C>A | TAGGTGGTCC | Unclassified | VUS | Class C0 | - | 0.027 | Outside SPiCE Interpretation | AAGATTTTCTTACTTCAGAGAATTATTTGCCACGTATTTCTAGCCT | AGGCTAGAAATACGTGGCAAATAATTCTCTGAAGTAAGAAAATCTT |
| 20 | E462G | c.1385A>G | TGGCTCGGGC | Class 1 | Benign | Class C0 | - | 0.027 | Outside SPiCE Interpretation | CAGAGAAGCCATTAAATGAGGGAACAGTGGTAAATAAGAGAGA | TCTCTCTTATTTACCACTGTTCCCTCATTTAATGGCTTCTCTG |
| 21 | N588D | c.1762A>G | TGGCGAGAGG | Class 2 | Likely benign | Class C0 | - | 0.027 | Outside SPiCE Interpretation | GGTTTAATATCCACTTTGAAAAAGAAAACAGATAAGTTTATTTATGCTATACATGATGAA | TTCATCATGTATAGCATAAATAAACTTATCTGTTTTCTTTTTCAAAGTGGATATTAAACC |
| 22 | H595Y | c.1783C>T | TGATGTCCGC | Unclassified | VUS | Class C0 | - | 0.027 | Outside SPiCE Interpretation | CCACTTTGAAAAAGAAAACAAATAAGTTTATTTATGCTATATATGATGAAACATCTTATAAAGG | CCTTTATAAGATGTTTCATCATATATAGCATAAATAAACTTATTTGTTTTCTTTTTCAAAGTGG |
| 23 | T598A | c.1792A>G | GTGTCTGTTG | Class 2 | Benign/Likely benign | Class C0 | - | 0.027 | Outside SPiCE Interpretation | ATAAGTTTATTTATGCTATACATGATGAAGCATCTTATAAAGGAAAAAAAATACCGAAAG | CTTTCGGTATTTTTTTTCCTTTATAAGATGCTTCATCATGTATAGCATAAATAAACTTAT |
| 24 | Y600C | c.1799A>G | TTATTAATGA | Unclassified | Conflicting | Class C0 | - | 0.027 | Outside SPiCE Interpretation | TTATGCTATACATGATGAAACATCTTGTAAAGGAAAAAAAATACCGAAAGACC | GGTCTTTCGGTATTTTTTTTCCTTTACAAGATGTTTCATCATGTATAGCATAA |
| 25 | G602R | c.1804G>A | TCTCCCGACC | Class 1 | Benign | Class C0 | - | 0.027 | Outside SPiCE Interpretation | TGCTATACATGATGAAACATCTTATAAAAGAAAAAAAATACCGAAAGACCAAAAATC | GATTTTTGGTCTTTCGGTATTTTTTTTCTTTTATAAGATGTTTCATCATGTATAGCA |
| 26 | G602V | c.1805G>T | TTCTTTGTCA | Unclassified | VUS | Class C0 | - | 0.027 | Outside SPiCE Interpretation | GCTATACATGATGAAACATCTTATAAAGTAAAAAAAATACCGAAAGACCAAAAATCA | TGATTTTTGGTCTTTCGGTATTTTTTTTACTTTATAAGATGTTTCATCATGTATAGC |
| 27 | E747G | c.2240A>G | CTGGGACATC | Unclassified | VUS | Class C0 | - | 0.027 | Outside SPiCE Interpretation | CCAGTACAACATTCAAAAGTGGGATACAGTGATACTGACTTTCAA | TTGAAAGTCAGTATCACTGTATCCCACTTTTGAATGTTGTACTGG |
| 28 | S755C | c.2264C>G | TGCCTAGTTT | Unclassified | VUS | Class C0 | - | 0.027 | Outside SPiCE Interpretation | AAAAGACTTTTCTGGCATTGAAAGTCAGTATCACTGTATTCCACT | AGTGGAATACAGTGATACTGACTTTCAATGCCAGAAAAGTCTTTT |
| 29 | N854S | c.2561A>G | ACCGGTCAGC | Unclassified | VUS | Class C0 | - | 0.027 | Outside SPiCE Interpretation | CAAGAAAGGTACAATTCAACCAAAGCACAAATCTAAGAGTAATCCAAA | TTTGGATTACTCTTAGATTTGTGCTTTGGTTGAATTGTACCTTTCTTG |
| 30 | L1019V | c.3055C>G | ACTTGGAATC | Class 1 | Benign | Class C0 | - | 0.027 | Outside SPiCE Interpretation | CAGCTTCAAATAAGGAAATCAAGGTCTCTGAACATAACATTAAGAAG | CTTCTTAATGTTATGTTCAGAGACCTTGATTTCCTTATTTGAAGCTG |
| 31 | S1074C | c.3220A>T | AATAGGTTTT | Unclassified | VUS | Class C0 | - | 0.027 | Outside SPiCE Interpretation | TACTGTATCTGCACATTTACAGTGTAGTGTAGTTGTTTCTGATTG | CAATCAGAAACAACTACACTACACTGTAAATGTGCAGATACAGTA |
| 32 | I1167V | c.3499A>G | CTCTTGGGGG | Unclassified | VUS | Class C0 | - | 0.027 | Outside SPiCE Interpretation | GATGCTGATCTTCATGTCGTAATGAATGCCCCATCGA | TCGATGGGGCATTCATTACGACATGAAGATCAGCATC |
| 33 | M1168I | c.3504G>T | TGTGTGTTCA | Unclassified | Likely pathogenic* | Class C0 | - | 0.027 | Outside SPiCE Interpretation | GAGATGCTGATCTTCATGTCATAATTAATGCCCCATCGAT | ATCGATGGGGCATTAATTATGACATGAAGATCAGCATCTC |
| 34 | A1170V | c.3509C>T | TTCAGGCCCA | Unclassified | Conflicting | Class C0 | - | 0.027 | Outside SPiCE Interpretation | CTTCATGTCATAATGAATGTCCCATCGATTGGTCAGGTA | TACCTGACCAATCGATGGGACATTCATTATGACATGAAG |
| 35 | S1172L | c.3515C>T | TTTAGTCGGC | Class 1 | Benign | Class C0 | - | 0.027 | Outside SPiCE Interpretation | TCATAATGAATGCCCCATTGATTGGTCAGGTAGACAG | CTGTCTACCTGACCAATCAATGGGGCATTCATTATGA |
| 36 | N1228D | c.3682A>G | TTTGCTTTTG | Class 1 | Benign | Class C0 | - | 0.027 | Outside SPiCE Interpretation | GCTCATGGCACAAAACTGGATGTTTCTACTGAAGCTC | GAGCTTCAGTAGAAACATCCAGTTTTGTGCCATGAGC |
| 37 | R1329S | c.3987A>T | CTGCATTGTA | Unclassified | VUS | Class C0 | - | 0.027 | Outside SPiCE Interpretation | ATAACAAATATACTGCTGCCAGTAGTAATTCTCATAACTTAGAATTTGATG | CATCAAATTCTAAGTTATGAGAATTACTACTGGCAGCAGTATATTTGTTAT |
| 38 | D1420Y | c.4258G>T | TCGCTAGGGG | Class 1 | Benign | Class C0 | - | 0.027 | Outside SPiCE Interpretation | ACTGCTACTAAAACGGAGCAAAATATAAAATATTTTGAGACTTCTGATAC | GTATCAGAAGTCTCAAAATATTTTATATTTTGCTCCGTTTTAGTAGCAGT |
| 39 | F1524V | c.4570T>G | CCGCACATTC | Class 1 | Benign | Class C45 | - | 0.027 | Outside SPiCE Interpretation | AACCTACTCTGTTGGGTGTTCATACAGCTAGCGGG | CCCGCTAGCTGTATGAACACCCAACAGAGTAGGTT |
| 40 | K1530N | c.4590A>T | GTTGGGGGTG | Unclassified | Pathogenic* | Class C0 | - | 0.027 | Outside SPiCE Interpretation | GTTTTCATACAGCTAGCGGGAATAAAGTTAAAATTGCAAAGGAATC | GATTCCTTTGCAATTTTAACTTTATTCCCGCTAGCTGTATGAAAAC |
| 41 | V1532F | c.4594G>T | TTATCTCAGT | Unclassified | VUS | Class C0 | - | 0.027 | Outside SPiCE Interpretation | TTCATACAGCTAGCGGGAAAAAATTTAAAATTGCAAAGGAATCTTTG | CAAAGATTCCTTTGCAATTTTAAATTTTTTCCCGCTAGCTGTATGAA |
| 42 | K1690N | c.5070A>C | ACGCGGATTG | Class 1 | Benign | Class C0 | - | 0.027 | Outside SPiCE Interpretation | CAGACTTCATTACTTGAAGCAAACAAATGGCTTAGAGAAGGAATATT | AATATTCCTTCTCTAAGCCATTTGTTTGCTTCAAGTAATGAAGTCTG |
| 43 | E1695V | c.5084A>T | GATCATTGCG | Unclassified | VUS | Class C15 | - | 0.027 | Outside SPiCE Interpretation | TTGAAGCAAAAAAATGGCTTAGAGTAGGAATATTTGATGGTCAACC | GGTTGACCATCAAATATTCCTACTCTAAGCCATTTTTTTGCTTCAA |
| 44 | G1696V | c.5087G>T | TCGTGTCCCC | Unclassified | Pathogenic* | Class C0 | - | 0.027 | Outside SPiCE Interpretation | AGCAAAAAAATGGCTTAGAGAAGTAATATTTGATGGTCAACCAGAAA | TTTCTGGTTGACCATCAAATATTACTTCTCTAAGCCATTTTTTTGCT |
| 45 | D1699N | c.5095G>A | CGTGGTTAAG | Unclassified | Benign/Likely benign | Class C0 | - | 0.027 | Outside SPiCE Interpretation | CAAAAAAATGGCTTAGAGAAGGAATATTTAATGGTCAACCAGAAAG | CTTTCTGGTTGACCATTAAATATTCCTTCTCTAAGCCATTTTTTTG |
| 46 | D1728N | c.5182G>A | GATCGACCGC | Unclassified | VUS | Class C0 | - | 0.027 | Outside SPiCE Interpretation | GAAAATAATTCAAACAGTACTATAGCTGAAAATAACAAAAATCATCTCTCCGAA | TTCGGAGAGATGATTTTTGTTATTTTCAGCTATAGTACTGTTTGAATTATTTTC |
| 47 | D1737V | c.5210A>T | CTCGTCGGCG | Unclassified | VUS | Class C0 | - | 0.027 | Outside SPiCE Interpretation | AAAAATCATCTCTCCGAAAAACAAGTTACTTATTTAAGTAACAGTAGCATG | CATGCTACTGTTACTTAAATAAGTAACTTGTTTTTCGGAGAGATGATTTTT |
| 48 | N1878K | c.5634C>G | TGCGGGCCTA | Class 1 | Conflicting | Class C0 | - | 0.027 | Outside SPiCE Interpretation | ACAGTTTCAGTAAAGTAATTAAGGAAAACAAGGAGAATAAATCAAAAATTTGCC | GGCAAATTTTTGATTTATTCTCCTTGTTTTCCTTAATTACTTTACTGAAACTGT |
| 49 | H1918R | c.5753A>G | GCATTATGGT | Class 1 | Conflicting | Class C0 | - | 0.027 | Outside SPiCE Interpretation | GAATGTCAGCAAAAACCTTACGTGAATGCGTGCTACATTCATC | GATGAATGTAGCACGCATTCACGTAAGGTTTTTGCTGACATTC |
| 50 | S1970L | c.5909C>T | CGGAGGACGT | Unclassified | VUS | Class C0 | - | 0.027 | Outside SPiCE Interpretation | GAAGCTTCATAAGTCAGTCTTATCTGCAAATACTTGTGGGA | TCCCACAAGTATTTGCAGATAAGACTGACTTATGAAGCTTC |
| 51 | E2020K | c.6058G>A | CGGTGGCCTC | Class 2 | VUS | Class C0 | - | 0.027 | Outside SPiCE Interpretation | TTTCCAAAGTATTGTTTAAAAGTAACAAACATTCAGACCAGCTCACA | TGTGAGCTGGTCTGAATGTTTGTTACTTTTAAACAATACTTTGGAAA |
| 52 | H2074N | c.6220C>A | GGTGTCGCGT | Class 1 | Benign | Class C0 | - | 0.027 | Outside SPiCE Interpretation | AGCAAGTTTCCATTTTAGAAAGTTCCTTAAACAAAGTTAAGGGAGTG | CACTCCCTTAACTTTGTTTAAGGAACTTTCTAAAATGGAAACTTGCT |
| 53 | K2075N | c.6225A>C | GTCGTTCAGT | Unclassified | Conflicting | Class C0 | - | 0.027 | Outside SPiCE Interpretation | ATTTTAGAAAGTTCCTTACACAACGTTAAGGGAGTGTTAGAGGAATT | AATTCCTCTAACACTCCCTTAACGTTGTGTAAGGAACTTTCTAAAAT |
| 54 | V2138D | c.6413T>A | TTCTTAGGAG | Unclassified | VUS | Class C0 | - | 0.027 | Outside SPiCE Interpretation | GTAAAGAATTTAAATTATCAAATAACTTAAATGATGAAGGTGGTTCTTCAGAAAATAATC | GATTATTTTCTGAAGAACCACCTTCATCATTTAAGTTATTTGATAATTTAAATTCTTTAC |
| 55 | H2324R | c.6971A>G | TGTGGGATGG | Unclassified | VUS | Class C0 | - | 0.027 | Outside SPiCE Interpretation | GATCGAAGATTGTTTATGCATCGTGTTTCTTTAGAGCCGATTACC | GGTAATCGGCTCTAAAGAAACACGATGCATAAACAATCTTCGATC |
| 56 | P2329L | c.6986C>T | TTCATAGTTT | Unclassified | Conflicting | Class C0 | - | 0.027 | Outside SPiCE Interpretation | GCATCATGTTTCTTTAGAGCTGATTACCTGTGTACCCTTTC | GAAAGGGTACACAGGTAATCAGCTCTAAAGAAACATGATGC |
| 57 | R2336G | c.7006C>G | CGTACTTGGA | Unclassified | Likely benign | Class C0 | - | 0.087 | low | CGATTACCTGTGTACCCTTTGGCACAACTAAGGAAC | GTTCCTTAGTTGTGCCAAAGGGTACACAGGTAATCG |
| 58 | T2337A | c.7009A>G | CTTTGTGGTC | Unclassified | VUS | Class C0 | - | 0.017 | low | CCTGTGTACCCTTTCGCGCAACTAAGGAACGTCAA | TTGACGTTCCTTAGTTGCGCGAAAGGGTACACAGG |
| 59 | K2411T | c.7232A>C | TAGCGTGTGA | Class 1 | Conflicting | Class C0 | - | 0.027 | Outside SPiCE Interpretation | CAACCAAAGTCTTTGTTCCACCTTTTACAACTAAATCACATTTTCACAG | CTGTGAAAATGTGATTTAGTTGTAAAAGGTGGAACAAAGACTTTGGTTG |
| 60 | T2412A | c.7234A>G | TGCGGTTTTC | Unclassified | VUS | Class C0 | - | 0.027 | Outside SPiCE Interpretation | CAAAGTCTTTGTTCCACCTTTTAAAGCTAAATCACATTTTCACAGAGTTG | CAACTCTGTGAAAATGTGATTTAGCTTTAAAAGGTGGAACAAAGACTTTG |
| 61 | T2412I | c.7235C>T | GGCAGTGGTA | Unclassified | VUS | Class C0 | - | 0.027 | Outside SPiCE Interpretation | CCAAAGTCTTTGTTCCACCTTTTAAAATTAAATCACATTTTCACAGAGTTG | CAACTCTGTGAAAATGTGATTTAATTTTAAAAGGTGGAACAAAGACTTTGG |
| 62 | R2418G | c.7252A>G | GGCTTGAATG | Class 2 | Conflicting | Class C0 | - | 0.027 | Outside SPiCE Interpretation | CTTTTAAAACTAAATCACATTTTCACGGAGTTGAACAGTGTGTTAGGAATATT | AATATTCCTAACACACTGTTCAACTCCGTGAAAATGTGATTTAGTTTTAAAAG |
| 63 | N2436I | c.7307A>T | AAAAACATTG | Class 1 | Benign | Class C0 | - | 0.027 | Outside SPiCE Interpretation | GGAAAACAGACAAAAGCAAATCATTGATGGACATGGCTCTG | CAGAGCCATGTCCATCAATGATTTGCTTTTGTCTGTTTTCC |
| 64 | Y2601C | c.7802A>G | AGCTGAGATT | Unclassified | VUS | Class C65 | DNA binding domain | 0.027 | Outside SPiCE Interpretation | TGGAAAGGCTGGAAAAGAAGAATTTTGTAGGGCTCTGTGTG | CACACAGAGCCCTACAAAATTCTTCTTTTCCAGCCTTTCCA |
| 65 | L2604P | c.7811T>C | GCACCCGCTG | Unclassified | VUS | Class C35 | DNA binding domain | 0.027 | Outside SPiCE Interpretation | AGAATTTTATAGGGCTCCGTGTGACACTCCAGGTG | CACCTGGAGTGTCACACGGAGCCCTATAAAATTCT |
| 66 | G2609D | c.7826G>A | TTATAAGGTG | Class 4 | Likely pathogenic | Class C65 | DNA binding domain | 0.027 | Outside SPiCE Interpretation | TCTGTGTGACACTCCAGATGTGGATCCAAAGCTTA | TAAGCTTTGGATCCACATCTGGAGTGTCACACAGA |
| 67 | N2622S | c.7865A>G | AACAGTTTCA | Unclassified | VUS | Class C45 | DNA binding domain | 0.027 | Outside SPiCE Interpretation | AAAGCTTATTTCTAGAATTTGGGTTTATAGTCACTATAGATGGATCATATGG | CCATATGATCCATCTATAGTGACTATAAACCCAAATTCTAGAAATAAGCTTT |
| 68 | H2623R | c.7868A>G | TTTTGTCAGG | Unclassified | Conflicting | Class C25 | DNA binding domain | 0.027 | Outside SPiCE Interpretation | ATTTCTAGAATTTGGGTTTATAATCGCTATAGATGGATCATATGGAAACTG | CAGTTTCCATATGATCCATCTATAGCGATTATAAACCCAAATTCTAGAAAT |
| 69 | W2626R | c.7876T>A | GTGCATCTGT | Unclassified | VUS | Class C65 | DNA binding domain | 0.027 | Outside SPiCE Interpretation | TAGAATTTGGGTTTATAATCACTATAGAAGGATCATATGGAAACTGG | CCAGTTTCCATATGATCCTTCTATAGTGATTATAAACCCAAATTCTA |
| 70 | W2626C | c.7878G>C | TGTTTATTTT | Class 5 | Pathogenic | Class C65 | DNA binding domain | 0.027 | Outside SPiCE Interpretation | GGTTTATAATCACTATAGATGCATCATATGGAAACTGGCAGCT | AGCTGCCAGTTTCCATATGATGCATCTATAGTGATTATAAACC |
| 71 | I2627V | c.7879A>G | GCCGGGCTGG | Unclassified | VUS | Class C25 | DNA binding domain | 0.027 | Outside SPiCE Interpretation | GGTTTATAATCACTATAGATGGGTCATATGGAAACTGGCAGCTAT | ATAGCTGCCAGTTTCCATATGACCCATCTATAGTGATTATAAACC |
| 72 | P2639L | c.7916C>T | TTTACGTCTT | Unclassified | Conflicting | Class C65 | DNA binding domain | 0.027 | Outside SPiCE Interpretation | CAGCTATGGAATGTGCCTTTCTTAAGGAATTTGCTAATAGATG | CATCTATTAGCAAATTCCTTAAGAAAGGCACATTCCATAGCTG |
| 73 | E2663K | c.7987G>A | CGCTGGGCCT | Unclassified | VUS | Class C55 | DNA binding domain | 0.027 | Outside SPiCE Interpretation | CAACTAAAATACAGATATGATACGAAAATTGATAGAAGCAGAAGATCGG | CCGATCTTCTGCTTCTATCAATTTTCGTATCATATCTGTATTTTAGTTG |
| 74 | I2664M | c.7992T>G | GGTTTTGTTA | Unclassified | VUS | Class C0 | DNA binding domain | 0.027 | Outside SPiCE Interpretation | ATACAGATATGATACGGAAATGGATAGAAGCAGAAGATCGGC | GCCGATCTTCTGCTTCTATCCATTTCCGTATCATATCTGTAT |
| 75 | M2676T | c.8027T>C | CCTCTCTTAG | Class 2 | Conflicting | Class C0 | DNA binding domain | 0.027 | Outside SPiCE Interpretation | AGCAGAAGATCGGCTATAAAAAAGATAACGGAAAGGGATGACAC | GTGTCATCCCTTTCCGTTATCTTTTTTATAGCCGATCTTCTGCT |
| 76 | L2688P | c.8063T>C | TGAAGGCGGC | Class 4 | Conflicting | Class C65 | DNA binding domain | 0.027 | Outside SPiCE Interpretation | CAGCTGCAAAAACACTTGTTCCCTGTGTTTCTGACATAATTTC | GAAATTATGTCAGAAACACAGGGAACAAGTGTTTTTGCAGCTG |
| 77 | S2697N | c.8090G>A | CGCTCCCGGC | Class 2 | Conflicting | Class C0 | DNA binding domain | 0.027 | Outside SPiCE Interpretation | GTTCTCTGTGTTTCTGACATAATTTCATTGAACGCAAATATATCTGAA | TTCAGATATATTTGCGTTCAATGAAATTATGTCAGAAACACAGAGAAC |
| 78 | A2717S | c.8149G>T | GTCTTGATTT | Class 1 | Benign | Class C0 | DNA binding domain | 0.027 | Outside SPiCE Interpretation | AGTGCAGATACCCAAAAAGTGTCCATTATTGAACTTACAGATG | CATCTGTAAGTTCAATAATGGACACTTTTTGGGTATCTGCACT |
| 79 | L2721H | c.8162T>A | GTTTCAGATG | Unclassified | VUS | Class C25 | DNA binding domain | 0.027 | Outside SPiCE Interpretation | AAGTGGCCATTATTGAACATACAGATGGGTGGTATGC | GCATACCACCCATCTGTATGTTCAATAATGGCCACTT |
| 80 | T2722R | c.8165C>G | TGACTGGCCT | Class 5 | Pathogenic | Class C65 | DNA binding domain | 0.027 | Outside SPiCE Interpretation | CCCAAAAAGTGGCCATTATTGAACTTAGAGATGGGTGGTATG | CATACCACCCATCTCTAAGTTCAATAATGGCCACTTTTTGGG |
| 81 | T2722K | c.8165C>A | GTTAGTTGTA | Unclassified | Likely pathogenic | Class C65 | DNA binding domain | 0.027 | Outside SPiCE Interpretation | GATACCCAAAAAGTGGCCATTATTGAACTTAAAGATGGGTGGTATG | CATACCACCCATCTTTAAGTTCAATAATGGCCACTTTTTGGGTATC |
| 82 | D2723H | c.8167G>C | GGCTTTTCTT | Class 5 | Pathogenic | Class C65 | DNA binding domain | 0.027 | Outside SPiCE Interpretation | AGTGGCCATTATTGAACTTACACATGGGTGGTATGCTG | CAGCATACCACCCATGTGTAAGTTCAATAATGGCCACT |
| 83 | D2723E | c.8169T>A | GACTGTCGCG | Unclassified | Conflicting | Class C35 | DNA binding domain | 0.027 | Outside SPiCE Interpretation | GTGGCCATTATTGAACTTACAGAAGGGTGGTATGCT | AGCATACCACCCTTCTGTAAGTTCAATAATGGCCAC |
| 84 | G2724W | c.8170G>T | GCTTCCCATG | Unclassified | Likely pathogenic | Class C35 | DNA binding domain | 0.027 | Outside SPiCE Interpretation | GCCATTATTGAACTTACAGATTGGTGGTATGCTGTTAAGG | CCTTAACAGCATACCACCAATCTGTAAGTTCAATAATGGC |
| 85 | K2729N | c.8187G>T | GCTTGCGGGC | Class 1 | Benign | Class C0 | DNA binding domain | 0.027 | Outside SPiCE Interpretation | AGATGGGTGGTATGCTGTTAATGCCCAGTTAGATC | GATCTAACTGGGCATTAACAGCATACCACCCATCT |
| 86 | V2747I | c.8239G>A | TCGCGTTCGG | Unclassified | VUS | Class C0 | DNA binding domain | 0.027 | Outside SPiCE Interpretation | CTTAAAGAATGGCAGACTGACAATTGGTCAGAAGATTATTCTTCA | TGAAGAATAATCTTCTGACCAATTGTCAGTCTGCCATTCTTTAAG |
| 87 | G2748D | c.8243G>A | TCCATCGTTT | Class 5 | Pathogenic | Class C65 | DNA binding domain | 0.027 | Outside SPiCE Interpretation | GAATGGCAGACTGACAGTTGATCAGAAGATTATTCTTCATG | CATGAAGAATAATCTTCTGATCAACTGTCAGTCTGCCATTC |
| 88 | R2842H | c.8525G>A | GGTCTTGGAC | Class 1 | Benign | Class C25 | DNA binding domain | 0.027 | Outside SPiCE Interpretation | ATCATCTGGATTATACATATTTCACAATGAAAGAGAGGAAGAAAAGG | CCTTTTCTTCCTCTCTTTCATTGTGAAATATGTATAATCCAGATGAT |
| 89 | R2896C | c.8686C>T | ATTTTTTTCT | Unclassified | VUS | Class C0 | DNA binding domain | 0.027 | Outside SPiCE Interpretation | CGTGCACTAACAAGACAGCAAGTTTGTGCTTTGCAAGA | TCTTGCAAAGCACAAACTTGCTGTCTTGTTAGTGCACG |
| 90 | V2908G | c.8723T>G | CCGTGTCAGG | Class 2 | Likely benign | Class C35 | DNA binding domain | 0.027 | Outside SPiCE Interpretation | AGAGCTTTATGAAGCAGGGAAGAATGCAGCAGACC | GGTCTGCTGCATTCTTCCCTGCTTCATAAAGCTCT |
| 91 | A2911E | c.8732C>A | CCATCGCAGC | Unclassified | Pathogenic* | Class C0 | DNA binding domain | 0.027 | Outside SPiCE Interpretation | TGAAGCAGTGAAGAATGAAGCAGACCCAGCTTAC | GTAAGCTGGGTCTGCTTCATTCTTCACTGCTTCA |
| 92 | A2911V | c.8732C>T | TCCTGGGTTG | Unclassified | VUS | Class C0 | DNA binding domain | 0.027 | Outside SPiCE Interpretation | TGAAGCAGTGAAGAATGTAGCAGACCCAGCTTAC | GTAAGCTGGGTCTGCTACATTCTTCACTGCTTCA |
| 93 | D2913H | c.8737G>C | TTTGTGGCTG | Unclassified | VUS | Class C65 | DNA binding domain | 0.027 | Outside SPiCE Interpretation | TGAAGCAGTGAAGAATGCAGCACACCCAGCTTACC | GGTAAGCTGGGTGTGCTGCATTCTTCACTGCTTCA |
| 94 | I2944F | c.8830A>T | AACTCTACCT | Class 1 | Benign | Class C0 | DNA binding domain | 0.027 | Outside SPiCE Interpretation | AATGATAAGAAACAAGCTCAGTTCCAGTTGGAAATTAGGAAGG | CCTTCCTAATTTCCAACTGGAACTGAGCTTGTTTCTTATCATT |
| 95 | I3044V | c.9130A>G | CTGCTGTGGC | Unclassified | Likely benign | Class C0 | DNA binding domain | 0.027 | Outside SPiCE Interpretation | ACAACTACCGGTTTCAGATGAAGTTTTATTTCAGATTTACCAGCC | GGCTGGTAAATCTGAAATAAAACTTCATCTGAAACCGGTAGTTGT |

| | | | | | | | | | | | |
|---|---|---|---|---|---|---|---|---|---|---|---|
| 96 | R3052W | c.9154C>T | TCTGTTGACC | Class 5 | Pathogenic | Class C65 | DNA binding domain | 0.027 | Outside SPiCE Interpretation | GAAGGGGCTCCCATGGCTGGTAAATCTGAAATAAAA | TTTTATTTCAGATTTACCAGCCATGGGAGCCCCTTC |
| 97 | P3054S | c.9160C>T | TTGTTCTGGT | Unclassified | VUS | Class C0 | DNA binding domain | 0.027 | Outside SPiCE Interpretation | CCAGCCACGGGAGTCCCTTCACTTCAG | CTGAAGTGAAGGGACTCCCGTGGCTGG |
| 98 | D3095E | c.9285C>A | GGACATCATT | Class 5 | Pathogenic | Class C35 | DNA binding domain | 0.027 | Outside SPiCE Interpretation | CCCTTTCGTCTATTTGTCAGAAGAATGTTACAATTTACTGGCA | TGCCAGTAAATTGTAACATTCTTCTGACAAATAGACGAAAGGG |
| 99 | E3096K | c.9286G>A | AGGCGTTTCC | Class 2 | VUS | Class C0 | DNA binding domain | 0.027 | Outside SPiCE Interpretation | CCCTTTCGTCTATTTGTCAGACAAATGTTACAATTTACTGGCAAT | ATTGCCAGTAAATTGTAACATTTGTCTGACAAATAGACGAAAGGG |
| 100 | Y3098H | c.9292T>C | ACTTAGTCAT | Class 1 | Benign | Class C0 | DNA binding domain | 0.027 | Outside SPiCE Interpretation | CTTTCGTCTATTTGTCAGACGAATGTCACAATTTACTGGCAATAAA | TTTATTGCCAGTAAATTGTGACATTCGTCTGACAAATAGACGAAAG |
| 101 | N3124I | c.9371A>T | TTTTCTTTAT | Class 5 | Pathogenic/likely pathogenic | Class C65 | DNA binding domain | 0.027 | Outside SPiCE Interpretation | CATATGTTAATTGCTGCAAGCATCCTCCAGTGGCG | CGCCACTGGAGGATGCTTGCAGCAATTAACATATG |
| 102 | I3183V | c.9547A>G | CTTTTTTACT | Unclassified | Conflicting | Class C0 | DNA binding domain | 0.027 | Outside SPiCE Interpretation | CAATGAAGCAGAAAACAAGCTTATGCATGTACTGCATGCAAATGA | TCATTTGCATGCAGTACATGCATAAGCTTGTTTTCTGCTTCATTG |
| 103 | N3187K | c.9561T>A | CATTGTGTTT | Unclassified | Likely pathogenic | Class C0 | - | 0.027 | Outside SPiCE Interpretation | CTTATGCATATACTGCATGCAAAAGATCCCAAGTGGTCC | GGACCACTTGGGATCTTTTGCATGCAGTATATGCATAAG |
| 104 | S3291C | c.9872C>G | TCTTGTTGGG | Unclassified | VUS | Class C65 | TR2 RAD51 binding domain | 0.027 | Outside SPiCE Interpretation | TAGTCCCATTTGTACATTTGTTTGTCCGGCTGCACA | TGTGCAGCCGGACAAACAAATGTACAAATGGGACTA |
| 105 | P3292L | c.9875C>T | GCTGGTCCTC | Class 1 | Conflicting | Class C0 | TR2 RAD52 binding domain | 0.027 | Outside SPiCE Interpretation | CATTTGTACATTTGTTTCTCTGGCTGCACAGAAGGCATTTC | GAAATGCCTTCTGTGCAGCCAGAGAAACAAATGTACAAATG |
| 106 | S3319F | c.9956C>T | AGGTGGTCAG | Unclassified | VUS | Class C0 | - | 0.027 | Outside SPiCE Interpretation | AACACCCATAAAGAAAAAAGAACTGAATTTTCCTCAGATGACTCC | GGAGTCATCTGAGGAAAATTCAGTTCTTTTTTCTTTATGGGTGTT |
| 107 | R3385H | c.10154G>A | ATCATGCCGT | Unclassified | Conflicting | Class C0 | - | 0.027 | Outside SPiCE Interpretation | GATTATCTCAGACTGAAACGACATTGTACTACATCTCTGATCAAA | TTTGATCAGAGATGTAGTACAATGTCGTTTCAGTCTGAGATAATC |
| 108 | T3387A | c.10159A>G | CGCACAGCCT | Unclassified | VUS | Class C0 | - | 0.027 | Outside SPiCE Interpretation | CAGACTGAAACGACGTTGTGCTACATCTCTGATCAAAGA | TCTTTGATCAGAGATGTAGCACAACGTCGTTTCAGTCTG |

Align-GVGD key domains: PALB2 interaction domain (amino acid residues 10-40), DNA binding domain (2481-3186), TR2 RAD51 binding domain (3269-3305)

* ClinVar classification without assertion criteria

| # | Variant | HGVS mutation | Barcode | IARC classification | Align-GVGD Classification From Human To Sea Urchin | Align-GVGD key domains | SPiCE splice abberation probability | SPiCE interpretation | Mutagenesis primer-forward | Mutagenesis primer-reverse |
|---|---|---|---|---|---|---|---|---|---|---|
| 1 | Wild-type | - | TCGTTTGTCT | Class 1 | Class C0 | - | - | - | - | - |
| 2 | Empty-vector | - | GTGTGGTGGG | Class 5 | Non-sense | - | - | - | - | - |
| 3 | R18H | c.53G>A | ATCGTATTTC | Class 2 | Class C0 | PALB2 interaction domain | 0.027 | Outside SPiCE Interpretation | CATTTTTTGAAATTTTTAAGACACACTGCAACAAAGCAGATTTAGGACC | GGTCCTAAATCTGCTTTGTTGCAGTGTGTCTTAAAAATTTCAAAAAATG |
| 4 | K21R | c.62A>G | TCTGCGATTC | Unclassified | Class C0 | PALB2 interaction domain | 0.027 | Outside SPiCE Interpretation | TTTGAAATTTTTAAGACACGCTGCAACAGAGCAGATTTAGGACC | GGTCCTAAATCTGCTCTGTTGCAGCGTGTCTTAAAAATTTCAAA |
| 5 | W31C | c.93G>T | TCTTTATGGT | Unclassified | Class C65 | PALB2 interaction domain | 0.027 | Outside SPiCE Interpretation | TTAGGACCAATAAGTCTTAATTGTTTTGAAGAACTTTCTTCAGAAGC | GCTTCTGAAGAAAGTTCTTCAAAACAATTAAGACTTATTGGTCCTAA |
| 6 | P41L | c.122C>T | CCTTGTGCAT | Unclassified | Class C0 | - | 0.027 | Outside SPiCE Interpretation | CTTTCTTCAGAAGCTCCACTCTATAATTCTGAACCTGCA | TGCAGGTTCAGAATTATAGAGTGGAGCTTCTGAAGAAAG |
| 7 | Y42C | c.125A>G | GGGCGGTTGT | Class 1 | Class C0 | - | 0.027 | Outside SPiCE Interpretation | TTCAGAAGCTCCACCCTGTAATTCTGAACCTGCAG | CTGCAGGTTCAGAATTACAGGGTGGAGCTTCTGAA |
| 8 | N55S | c.164A>G | TTCCTTTACT | Unclassified | Class C0 | - | 0.027 | Outside SPiCE Interpretation | GCAGAAGAATCTGAACATAAAAACAGCAATTACGAACCAAACCTATTTAA | TTAAATAGGTTTGGTTCGTAATTGCTGTTTTTATGTTCAGATTCTTCTGC |
| 9 | P59A | c.175C>G | ATGGGACTTG | Unclassified | Class C0 | - | 0.027 | Outside SPiCE Interpretation | TGAACATAAAAACAACAATTACGAAGCAAACCTATTTAAAACTCCACAAAG | CTTTGTGGAGTTTTAAATAGGTTTGCTTCGTAATTGTTGTTTTTATGTTCA |
| 10 | N108S | c.323A>G | ATTCTGTTTC | Unclassified | Class C0 | - | 0.027 | Outside SPiCE Interpretation | ATTAGATAAATTCAAATTAGACTTAGGAAGGAGTGTTCCCAATAGTAGAC | GTCTACTATTGGGAACACTCCTTCCTAAGTCTAATTTGAATTTATCTAAT |
| 11 | Q147R | c.440A>G | CGCCTGACGT | Unclassified | Class C0 | - | 0.027 | Outside SPiCE Interpretation | TGAAAGTCCTGTTGTTCTACGATGTACACATGTAACACCAC | GTGGTGTTACATGTGTACATCGTAGAACAACAGGACTTTCA |
| 12 | D156G | c.467A>G | GCACTTTCTA | Unclassified | Class C0 | - | 0.027 | Outside SPiCE Interpretation | CATGTAACACCACAAAGAGGTAAGTCAGTGGTATGTGGG | CCCACATACCACTGACTTACCTCTTTGTGGTGTTACATG |
| 13 | V159E | c.476T>A | TGGCCGGTGC | Unclassified | Class C0 | - | 0.007 | low | CCACAAAGAGATAAGTCAGAGGTATGTGGGAGTTTGTTT | AAACAAACTCCCACATACCTCTGACTTATCTCTTTGTGG |
| 14 | P168T | c.502C>A | GGTACCTCGT | Class 1 | Class C0 | - | 0.027 | Outside SPiCE Interpretation | TGGGAGTTTGTTTCATACAACAAAGTTTGTGAAGGGTCG | CGACCCTTCACAAACTTTGTTGTATGAAACAAACTCCCA |
| 15 | V208G | c.623T>G | TTTGCTCGTT | Unclassified | Class C15 | - | 0.027 | Outside SPiCE Interpretation | CCACCCTTAGTTCTACTGGGCTCATAGTCAGAAATGA | TCATTTCTGACTATGAGCCCAGTAGAACTAAGGGTGG |
| 16 | F266L | c.796T>C | TAGCCTCTGC | Unclassified | Class C0 | - | 0.023 | Outside SPiCE Interpretation | GAAGCTGCAAGTCATGGACTTGGAAAAACATCAGGGA | TCCCTGATGTTTTTCCAAGTCCATGACTTGCAGCTTC |
| 17 | R324T | c.971G>C | GTTGAGGTTG | Unclassified | Class C0 | - | 0.027 | Outside SPiCE Interpretation | TCTAAATGTAGAACAAAAAATCTACAAAAAGTAACAACTAGCAAGACTAGGAAAAA | TTTTTCCTAGTCTTGCTAGTTGTTACTTTTTGTAGATTTTTTGTTCTACATTTAGA |
| 18 | S326R | c.978C>A | TTGAATGTCC | Class 1 | Class C0 | - | 0.027 | Outside SPiCE Interpretation | CAAAAAATCTACAAAAAGTAAGAACTAGAAAGACTAGGAAAAAAATTTTCCATGAAG | CTTCATGGAAAATTTTTTTCCTAGTCTTTCTAGTTCTTACTTTTTGTAGATTTTTTG |
| 19 | S445Y | c.1334C>A | TAGGTGGTCC | Unclassified | Class C0 | - | 0.027 | Outside SPiCE Interpretation | AAGATTTTCTTACTTCAGAGAATTATTTGCCACGTATTTCTAGCCT | AGGCTAGAAATACGTGGCAAATAATTCTCTGAAGTAAGAAAATCTT |
| 20 | E462G | c.1385A>G | TGGCTCGGGC | Class 1 | Class C0 | - | 0.027 | Outside SPiCE Interpretation | CAGAGAAGCCATTAAATGAGGGAACAGTGGTAAATAAGAGAGA | TCTCTCTTATTTACCACTGTTCCCTCATTTAATGGCTTCTCTG |
| 21 | N588D | c.1762A>G | TGGCGAGAGG | Class 2 | Class C0 | - | 0.027 | Outside SPiCE Interpretation | GGTTTAATATCCACTTTGAAAAAGAAAACAGATAAGTTTATTTATGCTATACATGATGAA | TTCATCATGTATAGCATAAATAAACTTATCTGTTTTCTTTTTCAAAGTGGATATTAAACC |
| 22 | H595Y | c.1783C>T | TGATGTCCGC | Unclassified | Class C0 | - | 0.027 | Outside SPiCE Interpretation | CCACTTTGAAAAAGAAAACAAATAAGTTTATTTATGCTATATATGATGAAACATCTTATAAAGG | CCTTTATAAGATGTTTCATCATATATAGCATAAATAAACTTATTTGTTTTCTTTTTCAAAGTGG |
| 23 | T598A | c.1792A>G | GTGTCTGTTG | Class 2 | Class C0 | - | 0.027 | Outside SPiCE Interpretation | ATAAGTTTATTTATGCTATACATGATGAAGCATCTTATAAAGGAAAAAAAATACCGAAAG | CTTTCGGTATTTTTTTTCCTTTATAAGATGCTTCATCATGTATAGCATAAATAAACTTAT |
| 24 | Y600C | c.1799A>G | TTATTAATGA | Unclassified | Class C0 | - | 0.027 | Outside SPiCE Interpretation | TTATGCTATACATGATGAAACATCTTGTAAAGGAAAAAAAATACCGAAAGACC | GGTCTTTCGGTATTTTTTTTCCTTTACAAGATGTTTCATCATGTATAGCATAA |
| 25 | G602R | c.1804G>A | TCTCCCGACC | Class 1 | Class C0 | - | 0.027 | Outside SPiCE Interpretation | TGCTATACATGATGAAACATCTTATAAAAGAAAAAAAATACCGAAAGACCAAAAATC | GATTTTTGGTCTTTCGGTATTTTTTTTCTTTTATAAGATGTTTCATCATGTATAGCA |
| 26 | G602V | c.1805G>T | TTCTTTGTCA | Unclassified | Class C0 | - | 0.027 | Outside SPiCE Interpretation | GCTATACATGATGAAACATCTTATAAAGTAAAAAAAATACCGAAAGACCAAAAATCA | TGATTTTTGGTCTTTCGGTATTTTTTTTACTTTATAAGATGTTTCATCATGTATAGC |
| 27 | E747G | c.2240A>G | CTGGGACATC | Unclassified | Class C0 | - | 0.027 | Outside SPiCE Interpretation | CCAGTACAACATTCAAAAGTGGGATACAGTGATACTGACTTTCAA | TTGAAAGTCAGTATCACTGTATCCCACTTTTGAATGTTGTACTGG |
| 28 | S755C | c.2264C>G | TGCCTAGTTT | Unclassified | Class C0 | - | 0.027 | Outside SPiCE Interpretation | AAAAGACTTTTCTGGCATTGAAAGTCAGTATCACTGTATTCCACT | AGTGGAATACAGTGATACTGACTTTCAATGCCAGAAAAGTCTTTT |
| 29 | N854S | c.2561A>G | ACCGGTCAGC | Unclassified | Class C0 | - | 0.027 | Outside SPiCE Interpretation | CAAGAAAGGTACAATTCAACCAAAGCACAAATCTAAGAGTAATCCAAA | TTTGGATTACTCTTAGATTTGTGCTTTGGTTGAATTGTACCTTTCTTG |
| 30 | L1019V | c.3055C>G | ACTTGGAATC | Class 1 | Class C0 | - | 0.027 | Outside SPiCE Interpretation | CAGCTTCAAATAAGGAAATCAAGGTCTCTGAACATAACATTAAGAAG | CTTCTTAATGTTATGTTCAGAGACCTTGATTTCCTTATTTGAAGCTG |
| 31 | S1074C | c.3220A>T | AATAGGTTTT | Unclassified | Class C0 | - | 0.027 | Outside SPiCE Interpretation | TACTGTATCTGCACATTTACAGTGTAGTGTAGTTGTTTCTGATTG | CAATCAGAAACAACTACACTACACTGTAAATGTGCAGATACAGTA |
| 32 | I1167V | c.3499A>G | CTCTTGGGGG | Unclassified | Class C0 | - | 0.027 | Outside SPiCE Interpretation | GATGCTGATCTTCATGTCGTAATGAATGCCCCATCGA | TCGATGGGGCATTCATTACGACATGAAGATCAGCATC |
| 33 | M1168I | c.3504G>T | TGTGTGTTCA | Unclassified | Class C0 | - | 0.027 | Outside SPiCE Interpretation | GAGATGCTGATCTTCATGTCATAATTAATGCCCCATCGAT | ATCGATGGGGCATTAATTATGACATGAAGATCAGCATCTC |
| 34 | A1170V | c.3509C>T | TTCAGGCCCA | Unclassified | Class C0 | - | 0.027 | Outside SPiCE Interpretation | CTTCATGTCATAATGAATGTCCCATCGATTGGTCAGGTA | TACCTGACCAATCGATGGGACATTCATTATGACATGAAG |
| 35 | S1172L | c.3515C>T | TTTAGTCGGC | Class 1 | Class C0 | - | 0.027 | Outside SPiCE Interpretation | TCATAATGAATGCCCCATTGATTGGTCAGGTAGACAG | CTGTCTACCTGACCAATCAATGGGGCATTCATTATGA |
| 36 | N1228D | c.3682A>G | TTTGCTTTTG | Class 1 | Class C0 | - | 0.027 | Outside SPiCE Interpretation | GCTCATGGCACAAAACTGGATGTTTCTACTGAAGCTC | GAGCTTCAGTAGAAACATCCAGTTTTGTGCCATGAGC |
| 37 | R1329S | c.3987A>T | CTGCATTGTA | Unclassified | Class C0 | - | 0.027 | Outside SPiCE Interpretation | ATAACAAATATACTGCTGCCAGTAGTAATTCTCATAACTTAGAATTTGATG | CATCAAATTCTAAGTTATGAGAATTACTACTGGCAGCAGTATATTTGTTAT |
| 38 | D1420Y | c.4258G>T | TCGCTAGGGG | Class 1 | Class C0 | - | 0.027 | Outside SPiCE Interpretation | ACTGCTACTAAAACGGAGCAAAATATAAAATATTTTGAGACTTCTGATAC | GTATCAGAAGTCTCAAAATATTTTATATTTTGCTCCGTTTTAGTAGCAGT |
| 39 | F1524V | c.4570T>G | CCGCACATTC | Class 1 | Class C45 | - | 0.027 | Outside SPiCE Interpretation | AACCTACTCTGTTGGGTGTTCATACAGCTAGCGGG | CCCGCTAGCTGTATGAACACCCAACAGAGTAGGTT |
| 40 | K1530N | c.4590A>T | GTTGGGGGTG | Unclassified | Class C0 | - | 0.027 | Outside SPiCE Interpretation | GTTTTCATACAGCTAGCGGGAATAAAGTTAAAATTGCAAAGGAATC | GATTCCTTTGCAATTTTAACTTTATTCCCGCTAGCTGTATGAAAAC |
| 41 | V1532F | c.4594G>T | TTATCTCAGT | Unclassified | Class C0 | - | 0.027 | Outside SPiCE Interpretation | TTCATACAGCTAGCGGGAAAAAATTTAAAATTGCAAAGGAATCTTTG | CAAAGATTCCTTTGCAATTTTAAATTTTTTCCCGCTAGCTGTATGAA |
| 42 | K1690N | c.5070A>C | ACGCGGATTG | Class 1 | Class C0 | - | 0.027 | Outside SPiCE Interpretation | CAGACTTCATTACTTGAAGCAAACAAATGGCTTAGAGAAGGAATATT | AATATTCCTTCTCTAAGCCATTTGTTTGCTTCAAGTAATGAAGTCTG |
| 43 | E1695V | c.5084A>T | GATCATTGCG | Unclassified | Class C15 | - | 0.027 | Outside SPiCE Interpretation | TTGAAGCAAAAAAATGGCTTAGAGTAGGAATATTTGATGGTCAACC | GGTTGACCATCAAATATTCCTACTCTAAGCCATTTTTTTGCTTCAA |
| 44 | G1696V | c.5087G>T | TCGTGTCCCC | Unclassified | Class C0 | - | 0.027 | Outside SPiCE Interpretation | AGCAAAAAAATGGCTTAGAGAAGTAATATTTGATGGTCAACCAGAAA | TTTCTGGTTGACCATCAAATATTACTTCTCTAAGCCATTTTTTTGCT |
| 45 | D1699N | c.5095G>A | CGTGGTTAAG | Unclassified | Class C0 | - | 0.027 | Outside SPiCE Interpretation | CAAAAAAATGGCTTAGAGAAGGAATATTTAATGGTCAACCAGAAAG | CTTTCTGGTTGACCATTAAATATTCCTTCTCTAAGCCATTTTTTTG |
| 46 | D1728N | c.5182G>A | GATCGACCGC | Unclassified | Class C0 | - | 0.027 | Outside SPiCE Interpretation | GAAAATAATTCAAACAGTACTATAGCTGAAAATAACAAAAATCATCTCTCCGAA | TTCGGAGAGATGATTTTTGTTATTTTCAGCTATAGTACTGTTTGAATTATTTTC |
| 47 | D1737V | c.5210A>T | CTCGTCGGCG | Unclassified | Class C0 | - | 0.027 | Outside SPiCE Interpretation | AAAAATCATCTCTCCGAAAAACAAGTTACTTATTTAAGTAACAGTAGCATG | CATGCTACTGTTACTTAAATAAGTAACTTGTTTTTCGGAGAGATGATTTTT |
| 48 | N1878K | c.5634C>G | TGCGGGCCTA | Class 1 | Class C0 | - | 0.027 | Outside SPiCE Interpretation | ACAGTTTCAGTAAAGTAATTAAGGAAAACAAGGAGAATAAATCAAAAATTTGCC | GGCAAATTTTTGATTTATTCTCCTTGTTTTCCTTAATTACTTTACTGAAACTGT |
| 49 | H1918R | c.5753A>G | GCATTATGGT | Class 1 | Class C0 | - | 0.027 | Outside SPiCE Interpretation | GAATGTCAGCAAAAACCTTACGTGAATGCGTGCTACATTCATC | GATGAATGTAGCACGCATTCACGTAAGGTTTTTGCTGACATTC |
| 50 | S1970L | c.5909C>T | CGGAGGACGT | Unclassified | Class C0 | - | 0.027 | Outside SPiCE Interpretation | GAAGCTTCATAAGTCAGTCTTATCTGCAAATACTTGTGGGA | TCCCACAAGTATTTGCAGATAAGACTGACTTATGAAGCTTC |
| 51 | E2020K | c.6058G>A | CGGTGGCCTC | Class 2 | Class C0 | - | 0.027 | Outside SPiCE Interpretation | TTTCCAAAGTATTGTTTAAAAGTAACAAACATTCAGACCAGCTCACA | TGTGAGCTGGTCTGAATGTTTGTTACTTTTAAACAATACTTTGGAAA |
| 52 | H2074N | c.6220C>A | GGTGTCGCGT | Class 1 | Class C0 | - | 0.027 | Outside SPiCE Interpretation | AGCAAGTTTCCATTTTAGAAAGTTCCTTAAACAAAGTTAAGGGAGTG | CACTCCCTTAACTTTGTTTAAGGAACTTTCTAAAATGGAAACTTGCT |
| 53 | K2075N | c.6225A>C | GTCGTTCAGT | Unclassified | Class C0 | - | 0.027 | Outside SPiCE Interpretation | ATTTTAGAAAGTTCCTTACACAACGTTAAGGGAGTGTTAGAGGAATT | AATTCCTCTAACACTCCCTTAACGTTGTGTAAGGAACTTTCTAAAAT |
| 54 | V2138D | c.6413T>A | TTCTTAGGAG | Unclassified | Class C0 | - | 0.027 | Outside SPiCE Interpretation | GTAAAGAATTTAAATTATCAAATAACTTAAATGATGAAGGTGGTTCTTCAGAAAATAATC | GATTATTTTCTGAAGAACCACCTTCATCATTTAAGTTATTTGATAATTTAAATTCTTTAC |
| 55 | H2324R | c.6971A>G | TGTGGGATGG | Unclassified | Class C0 | - | 0.027 | Outside SPiCE Interpretation | GATCGAAGATTGTTTATGCATCGTGTTTCTTTAGAGCCGATTACC | GGTAATCGGCTCTAAAGAAACACGATGCATAAACAATCTTCGATC |
| 56 | P2329L | c.6986C>T | TTCATAGTTT | Unclassified | Class C0 | - | 0.027 | Outside SPiCE Interpretation | GCATCATGTTTCTTTAGAGCTGATTACCTGTGTACCCTTTC | GAAAGGGTACACAGGTAATCAGCTCTAAAGAAACATGATGC |
| 57 | R2336G | c.7006C>G | CGTACTTGGA | Unclassified | Class C0 | - | 0.087 | low | CGATTACCTGTGTACCCTTTGGCACAACTAAGGAAC | GTTCCTTAGTTGTGCCAAAGGGTACACAGGTAATCG |
| 58 | T2337A | c.7009A>G | CTTTGTGGTC | Unclassified | Class C0 | - | 0.017 | low | CCTGTGTACCCTTTCGCGCAACTAAGGAACGTCAA | TTGACGTTCCTTAGTTGCGCGAAAGGGTACACAGG |
| 59 | K2411T | c.7232A>C | TAGCGTGTGA | Class 1 | Class C0 | - | 0.027 | Outside SPiCE Interpretation | CAACCAAAGTCTTTGTTCCACCTTTTACAACTAAATCACATTTTCACAG | CTGTGAAAATGTGATTTAGTTGTAAAAGGTGGAACAAAGACTTTGGTTG |
| 60 | T2412A | c.7234A>G | TGCGGTTTTC | Unclassified | Class C0 | - | 0.027 | Outside SPiCE Interpretation | CAAAGTCTTTGTTCCACCTTTTAAAGCTAAATCACATTTTCACAGAGTTG | CAACTCTGTGAAAATGTGATTTAGCTTTAAAAGGTGGAACAAAGACTTTG |
| 61 | T2412I | c.7235C>T | GGCAGTGGTA | Unclassified | Class C0 | - | 0.027 | Outside SPiCE Interpretation | CCAAAGTCTTTGTTCCACCTTTTAAAATTAAATCACATTTTCACAGAGTTG | CAACTCTGTGAAAATGTGATTTAATTTTAAAAGGTGGAACAAAGACTTTGG |
| 62 | R2418G | c.7252A>G | GGCTTGAATG | Class 2 | Class C0 | - | 0.027 | Outside SPiCE Interpretation | CTTTTAAAACTAAATCACATTTTCACGGAGTTGAACAGTGTGTTAGGAATATT | AATATTCCTAACACACTGTTCAACTCCGTGAAAATGTGATTTAGTTTTAAAAG |
| 63 | N2436I | c.7307A>T | AAAAACATTG | Class 1 | Class C0 | - | 0.027 | Outside SPiCE Interpretation | GGAAAACAGACAAAAGCAAATCATTGATGGACATGGCTCTG | CAGAGCCATGTCCATCAATGATTTGCTTTTGTCTGTTTTCC |
| 64 | Y2601C | c.7802A>G | AGCTGAGATT | Unclassified | Class C65 | DNA binding domain | 0.027 | Outside SPiCE Interpretation | TGGAAAGGCTGGAAAAGAAGAATTTTGTAGGGCTCTGTGTG | CACACAGAGCCCTACAAAATTCTTCTTTTCCAGCCTTTCCA |
| 65 | L2604P | c.7811T>C | GCACCCGCTG | Unclassified | Class C35 | DNA binding domain | 0.027 | Outside SPiCE Interpretation | AGAATTTTATAGGGCTCCGTGTGACACTCCAGGTG | CACCTGGAGTGTCACACGGAGCCCTATAAAATTCT |
| 66 | G2609D | c.7826G>A | TTATAAGGTG | Class 4 | Class C65 | DNA binding domain | 0.027 | Outside SPiCE Interpretation | TCTGTGTGACACTCCAGATGTGGATCCAAAGCTTA | TAAGCTTTGGATCCACATCTGGAGTGTCACACAGA |
| 67 | N2622S | c.7865A>G | AACAGTTTCA | Unclassified | Class C45 | DNA binding domain | 0.027 | Outside SPiCE Interpretation | AAAGCTTATTTCTAGAATTTGGGTTTATAGTCACTATAGATGGATCATATGG | CCATATGATCCATCTATAGTGACTATAAACCCAAATTCTAGAAATAAGCTTT |
| 68 | H2623R | c.7868A>G | TTTTGTCAGG | Unclassified | Class C25 | DNA binding domain | 0.027 | Outside SPiCE Interpretation | ATTTCTAGAATTTGGGTTTATAATCGCTATAGATGGATCATATGGAAACTG | CAGTTTCCATATGATCCATCTATAGCGATTATAAACCCAAATTCTAGAAAT |
| 69 | W2626R | c.7876T>A | GTGCATCTGT | Unclassified | Class C65 | DNA binding domain | 0.027 | Outside SPiCE Interpretation | TAGAATTTGGGTTTATAATCACTATAGAAGGATCATATGGAAACTGG | CCAGTTTCCATATGATCCTTCTATAGTGATTATAAACCCAAATTCTA |
| 70 | W2626C | c.7878G>C | TGTTTATTTT | Class 5 | Class C65 | DNA binding domain | 0.027 | Outside SPiCE Interpretation | GGTTTATAATCACTATAGATGCATCATATGGAAACTGGCAGCT | AGCTGCCAGTTTCCATATGATGCATCTATAGTGATTATAAACC |
| 71 | I2627V | c.7879A>G | GCCGGGCTGG | Unclassified | Class C25 | DNA binding domain | 0.027 | Outside SPiCE Interpretation | GGTTTATAATCACTATAGATGGGTCATATGGAAACTGGCAGCTAT | ATAGCTGCCAGTTTCCATATGACCCATCTATAGTGATTATAAACC |
| 72 | P2639L | c.7916C>T | TTTACGTCTT | Unclassified | Class C65 | DNA binding domain | 0.027 | Outside SPiCE Interpretation | CAGCTATGGAATGTGCCTTTCTTAAGGAATTTGCTAATAGATG | CATCTATTAGCAAATTCCTTAAGAAAGGCACATTCCATAGCTG |
| 73 | E2663K | c.7987G>A | CGCTGGGCCT | Unclassified | Class C55 | DNA binding domain | 0.027 | Outside SPiCE Interpretation | CAACTAAAATACAGATATGATACGAAAATTGATAGAAGCAGAAGATCGG | CCGATCTTCTGCTTCTATCAATTTTCGTATCATATCTGTATTTTAGTTG |
| 74 | I2664M | c.7992T>G | GGTTTTGTTA | Unclassified | Class C0 | DNA binding domain | 0.027 | Outside SPiCE Interpretation | ATACAGATATGATACGGAAATGGATAGAAGCAGAAGATCGGC | GCCGATCTTCTGCTTCTATCCATTTCCGTATCATATCTGTAT |
| 75 | M2676T | c.8027T>C | CCTCTCTTAG | Class 2 | Class C0 | DNA binding domain | 0.027 | Outside SPiCE Interpretation | AGCAGAAGATCGGCTATAAAAAAGATAACGGAAAGGGATGACAC | GTGTCATCCCTTTCCGTTATCTTTTTTATAGCCGATCTTCTGCT |
| 76 | L2688P | c.8063T>C | TGAAGGCGGC | Class 4 | Class C65 | DNA binding domain | 0.027 | Outside SPiCE Interpretation | CAGCTGCAAAAACACTTGTTCCCTGTGTTTCTGACATAATTTC | GAAATTATGTCAGAAACACAGGGAACAAGTGTTTTTGCAGCTG |
| 77 | S2697N | c.8090G>A | CGCTCCCGGC | Class 2 | Class C0 | DNA binding domain | 0.027 | Outside SPiCE Interpretation | GTTCTCTGTGTTTCTGACATAATTTCATTGAACGCAAATATATCTGAA | TTCAGATATATTTGCGTTCAATGAAATTATGTCAGAAACACAGAGAAC |
| 78 | A2717S | c.8149G>T | GTCTTGATTT | Class 1 | Class C0 | DNA binding domain | 0.027 | Outside SPiCE Interpretation | AGTGCAGATACCCAAAAAGTGTCCATTATTGAACTTACAGATG | CATCTGTAAGTTCAATAATGGACACTTTTTGGGTATCTGCACT |
| 79 | L2721H | c.8162T>A | GTTTCAGATG | Unclassified | Class C25 | DNA binding domain | 0.027 | Outside SPiCE Interpretation | AAGTGGCCATTATTGAACATACAGATGGGTGGTATGC | GCATACCACCCATCTGTATGTTCAATAATGGCCACTT |
| 80 | T2722R | c.8165C>G | TGACTGGCCT | Class 5 | Class C65 | DNA binding domain | 0.027 | Outside SPiCE Interpretation | CCCAAAAAGTGGCCATTATTGAACTTAGAGATGGGTGGTATG | CATACCACCCATCTCTAAGTTCAATAATGGCCACTTTTTGGG |
| 81 | T2722K | c.8165C>A | GTTAGTTGTA | Unclassified | Class C65 | DNA binding domain | 0.027 | Outside SPiCE Interpretation | GATACCCAAAAAGTGGCCATTATTGAACTTAAAGATGGGTGGTATG | CATACCACCCATCTTTAAGTTCAATAATGGCCACTTTTTGGGTATC |
| 82 | D2723H | c.8167G>C | GGCTTTTCTT | Class 5 | Class C65 | DNA binding domain | 0.027 | Outside SPiCE Interpretation | AGTGGCCATTATTGAACTTACACATGGGTGGTATGCTG | CAGCATACCACCCATGTGTAAGTTCAATAATGGCCACT |
| 83 | D2723E | c.8169T>A | GACTGTCGCG | Unclassified | Class C35 | DNA binding domain | 0.027 | Outside SPiCE Interpretation | GTGGCCATTATTGAACTTACAGAAGGGTGGTATGCT | AGCATACCACCCTTCTGTAAGTTCAATAATGGCCAC |
| 84 | G2724W | c.8170G>T | GCTTCCCATG | Unclassified | Class C35 | DNA binding domain | 0.027 | Outside SPiCE Interpretation | GCCATTATTGAACTTACAGATTGGTGGTATGCTGTTAAGG | CCTTAACAGCATACCACCAATCTGTAAGTTCAATAATGGC |
| 85 | K2729N | c.8187G>T | GCTTGCGGGC | Class 1 | Class C0 | DNA binding domain | 0.027 | Outside SPiCE Interpretation | AGATGGGTGGTATGCTGTTAATGCCCAGTTAGATC | GATCTAACTGGGCATTAACAGCATACCACCCATCT |
| 86 | V2747I | c.8239G>A | TCGCGTTCGG | Unclassified | Class C0 | DNA binding domain | 0.027 | Outside SPiCE Interpretation | CTTAAAGAATGGCAGACTGACAATTGGTCAGAAGATTATTCTTCA | TGAAGAATAATCTTCTGACCAATTGTCAGTCTGCCATTCTTTAAG |
| 87 | G2748D | c.8243G>A | TCCATCGTTT | Class 5 | Class C65 | DNA binding domain | 0.027 | Outside SPiCE Interpretation | GAATGGCAGACTGACAGTTGATCAGAAGATTATTCTTCATG | CATGAAGAATAATCTTCTGATCAACTGTCAGTCTGCCATTC |
| 88 | R2842H | c.8525G>A | GGTCTTGGAC | Class 1 | Class C25 | DNA binding domain | 0.027 | Outside SPiCE Interpretation | ATCATCTGGATTATACATATTTCACAATGAAAGAGAGGAAGAAAAGG | CCTTTTCTTCCTCTCTTTCATTGTGAAATATGTATAATCCAGATGAT |
| 89 | R2896C | c.8686C>T | ATTTTTTTCT | Unclassified | Class C0 | DNA binding domain | 0.027 | Outside SPiCE Interpretation | CGTGCACTAACAAGACAGCAAGTTTGTGCTTTGCAAGA | TCTTGCAAAGCACAAACTTGCTGTCTTGTTAGTGCACG |
| 90 | V2908G | c.8723T>G | CCGTGTCAGG | Class 2 | Class C35 | DNA binding domain | 0.027 | Outside SPiCE Interpretation | AGAGCTTTATGAAGCAGGGAAGAATGCAGCAGACC | GGTCTGCTGCATTCTTCCCTGCTTCATAAAGCTCT |
| 91 | A2911E | c.8732C>A | CCATCGCAGC | Unclassified | Class C0 | DNA binding domain | 0.027 | Outside SPiCE Interpretation | TGAAGCAGTGAAGAATGAAGCAGACCCAGCTTAC | GTAAGCTGGGTCTGCTTCATTCTTCACTGCTTCA |
| 92 | A2911V | c.8732C>T | TCCTGGGTTG | Unclassified | Class C0 | DNA binding domain | 0.027 | Outside SPiCE Interpretation | TGAAGCAGTGAAGAATGTAGCAGACCCAGCTTAC | GTAAGCTGGGTCTGCTACATTCTTCACTGCTTCA |
| 93 | D2913H | c.8737G>C | TTTGTGGCTG | Unclassified | Class C65 | DNA binding domain | 0.027 | Outside SPiCE Interpretation | TGAAGCAGTGAAGAATGCAGCACACCCAGCTTACC | GGTAAGCTGGGTGTGCTGCATTCTTCACTGCTTCA |
| 94 | I2944F | c.8830A>T | AACTCTACCT | Class 1 | Class C0 | DNA binding domain | 0.027 | Outside SPiCE Interpretation | AATGATAAGAAACAAGCTCAGTTCCAGTTGGAAATTAGGAAGG | CCTTCCTAATTTCCAACTGGAACTGAGCTTGTTTCTTATCATT |
| 95 | I3044V | c.9130A>G | CTGCTGTGGC | Unclassified | Class C0 | DNA binding domain | 0.027 | Outside SPiCE Interpretation | ACAACTACCGGTTTCAGATGAAGTTTTATTTCAGATTTACCAGCC | GGCTGGTAAATCTGAAATAAAACTTCATCTGAAACCGGTAGTTGT |
| 96 | R3052W | c.9154C>T | TCTGTTGACC | Class 5 | Class C65 | DNA binding domain | 0.027 | Outside SPiCE Interpretation | GAAGGGGCTCCCATGGCTGGTAAATCTGAAATAAAA | TTTTATTTCAGATTTACCAGCCATGGGAGCCCCTTC |
| 97 | P3054S | c.9160C>T | TTGTTCTGGT | Unclassified | Class C0 | DNA binding domain | 0.027 | Outside SPiCE Interpretation | CCAGCCACGGGAGTCCCTTCACTTCAG | CTGAAGTGAAGGGACTCCCGTGGCTGG |
| 98 | D3095E | c.9285C>A | GGACATCATT | Class 5 | Class C35 | DNA binding domain | 0.027 | Outside SPiCE Interpretation | CCCTTTCGTCTATTTGTCAGAAGAATGTTACAATTTACTGGCA | TGCCAGTAAATTGTAACATTCTTCTGACAAATAGACGAAAGGG |
| 99 | E3096K | c.9286G>A | AGGCGTTTCC | Class 2 | Class C0 | DNA binding domain | 0.027 | Outside SPiCE Interpretation | CCCTTTCGTCTATTTGTCAGACAAATGTTACAATTTACTGGCAAT | ATTGCCAGTAAATTGTAACATTTGTCTGACAAATAGACGAAAGGG |
| 100 | Y3098H | c.9292T>C | ACTTAGTCAT | Class 1 | Class C0 | DNA binding domain | 0.027 | Outside SPiCE Interpretation | CTTTCGTCTATTTGTCAGACGAATGTCACAATTTACTGGCAATAAA | TTTATTGCCAGTAAATTGTGACATTCGTCTGACAAATAGACGAAAG |
| 101 | N3124I | c.9371A>T | TTTTCTTTAT | Class 5 | Class C65 | DNA binding domain | 0.027 | Outside SPiCE Interpretation | CATATGTTAATTGCTGCAAGCATCCTCCAGTGGCG | CGCCACTGGAGGATGCTTGCAGCAATTAACATATG |
| 102 | I3183V | c.9547A>G | CTTTTTTACT | Unclassified | Class C0 | DNA binding domain | 0.027 | Outside SPiCE Interpretation | CAATGAAGCAGAAAACAAGCTTATGCATGTACTGCATGCAAATGA | TCATTTGCATGCAGTACATGCATAAGCTTGTTTTCTGCTTCATTG |
| 103 | N3187K | c.9561T>A | CATTGTGTTT | Unclassified | Class C0 | - | 0.027 | Outside SPiCE Interpretation | CTTATGCATATACTGCATGCAAAAGATCCCAAGTGGTCC | GGACCACTTGGGATCTTTTGCATGCAGTATATGCATAAG |
| 104 | S3291C | c.9872C>G | TCTTGTTGGG | Unclassified | Class C65 | TR2 RAD51 binding domain | 0.027 | Outside SPiCE Interpretation | TAGTCCCATTTGTACATTTGTTTGTCCGGCTGCACA | TGTGCAGCCGGACAAACAAATGTACAAATGGGACTA |
| 105 | P3292L | c.9875C>T | GCTGGTCCTC | Class 1 | Class C0 | TR2 RAD52 binding domain | 0.027 | Outside SPiCE Interpretation | CATTTGTACATTTGTTTCTCTGGCTGCACAGAAGGCATTTC | GAAATGCCTTCTGTGCAGCCAGAGAAACAAATGTACAAATG |
| 106 | S3319F | c.9956C>T | AGGTGGTCAG | Unclassified | Class C0 | - | 0.027 | Outside SPiCE Interpretation | AACACCCATAAAGAAAAAAGAACTGAATTTTCCTCAGATGACTCC | GGAGTCATCTGAGGAAAATTCAGTTCTTTTTTCTTTATGGGTGTT |
| 107 | R3385H | c.10154G>A | ATCATGCCGT | Unclassified | Class C0 | - | 0.027 | Outside SPiCE Interpretation | GATTATCTCAGACTGAAACGACATTGTACTACATCTCTGATCAAA | TTTGATCAGAGATGTAGTACAATGTCGTTTCAGTCTGAGATAATC |
| 108 | T3387A | c.10159A>G | CGCACAGCCT | Unclassified | Class C0 | - | 0.027 | Outside SPiCE Interpretation | CAGACTGAAACGACGTTGTGCTACATCTCTGATCAAAGA | TCTTTGATCAGAGATGTAGCACAACGTCGTTTCAGTCTG |
| 109 | R18C | c.52C>T | TCAGCTGGGA | Class 2 | Class C15 | PALB2 interaction domain | 0.027 | Outside SPiCE Interpretation | CTGCTTTGTTGCAGCATGTCTTAAAAATTTCAAAAAATGTTGGCCTC | GAGGCCAACATTTTTTGAAATTTTTAAGACATGCTGCAACAAAGCAG |
| 110 | G25R | c.73G>A | CGCGGGCTTT | Unclassified | Class C65 | PALB2 interaction domain | 0.027 | Outside SPiCE Interpretation | GACACGCTGCAACAAAGCAGATTTAAGACCAATAAGTCTTAATTGG | CCAATTAAGACTTATTGGTCTTAAATCTGCTTTGTTGCAGCGTGTC |
| 111 | N30H | c.88A>C | GAGCGTCAGG | Unclassified | Class C15 | PALB2 interaction domain | 0.027 | Outside SPiCE Interpretation | CAGATTTAGGACCAATAAGTCTTCATTGGTTTGAAGAACTTTCTTCAG | CTGAAGAAAGTTCTTCAAACCAATGAAGACTTATTGGTCCTAAATCTG |
| 112 | W31G | c.91T>G | TCCGCGCGCG | Unclassified | Class C65 | PALB2 interaction domain | 0.027 | Outside SPiCE Interpretation | CAAAGCAGATTTAGGACCAATAAGTCTTAATGGGTTTGAAGAACTTTC | GAAAGTTCTTCAAACCCATTAAGACTTATTGGTCCTAAATCTGCTTTG |
| 113 | W31L | c.92G>T | GGGCATATCG | Unclassified | Class C55 | PALB2 interaction domain | 0.027 | Outside SPiCE Interpretation | CAAAGCAGATTTAGGACCAATAAGTCTTAATTTGTTTGAAGAACTTTCTTC | GAAGAAAGTTCTTCAAACAAATTAAGACTTATTGGTCCTAAATCTGCTTTG |
| 114 | F32L | c.94T>C | TGTTGTGGGG | Unclassified | Class C15 | PALB2 interaction domain | 0.027 | Outside SPiCE Interpretation | GATTTAGGACCAATAAGTCTTAATTGGCTTGAAGAACTTTCTTCAGAAGC | GCTTCTGAAGAAAGTTCTTCAAGCCAATTAAGACTTATTGGTCCTAAATC |
| 115 | T77I | c.230C>T | TTTGTGTTTG | Unclassified | Class C65 | - | 0.027 | Outside SPiCE Interpretation | GAAACCATCTTATAATCAGCTGGCTTCAATTCCAATAATATTCAAAGAGC | GCTCTTTGAATATTATTGGAATTGAAGCCAGCTGATTATAAGATGGTTTC |

| | | | | | | | | | | | |
|---|---|---|---|---|---|---|---|---|---|---|---|
| 116 | G185V | c.554G>T | GGCTTGCTGG | Unclassified | Class C65 | - | | 0.027 | Outside SPiCE Interpretation | CATATTTCTGAAAGTCTAGTAGCTGAGGTGGATCCTGATATG | CATATCAGGATCCACCTCAGCTACTAGACTTTCAGAAATATG |
| 117 | D191V | c.572A>T | GTTGCGGTGG | Unclassified | Class C35 | - | | 0.027 | Outside SPiCE Interpretation | CTAGGAGCTGAGGTGGATCCTGTTATGTCTTGGTCAAGTTCTTTAGC | GCTAAAGAACTTGACCAAGACATAACAGGATCCACCTCAGCTCCTAG |
| 118 | T200I | c.599C>T | GTTGAGGCCT | Unclassified | Class C65 | - | | 0.027 | Outside SPiCE Interpretation | GATATGTCTTGGTCAAGTTCTTTAGCTATACCACCCACCCTTAGTTC | GAACTAAGGGTGGGTGGTATAGCTAAAGAACTTGACCAAGACATATC |
| 119 | T207I | c.620C>T | TCTGTTCTTT | Unclassified | Class C65 | - | | 0.027 | Outside SPiCE Interpretation | CTACACCACCCACCCTTAGTTCTATTGTGCTCATAGTCAGAAATG | CATTTCTGACTATGAGCACAATAGAACTAAGGGTGGGTGGTGTAG |
| 19-2 | S445Y | c.1334C>A | GGCCCGGTTC | Unclassified | Class C0 | - | | 0.027 | Outside SPiCE Interpretation | GAAAGATTTTCTTACTTCAGAGAATTATTTGCCACGTATTTCTAGCCTAC | GTAGGCTAGAAATACGTGGCAAATAATTCTCTGAAGTAAGAAAATCTTTC |
| 120 | F590C | c.1769T>G | CCGGTAGGCC | Unclassified | Class C65 | - | | 0.027 | Outside SPiCE Interpretation | CCACTTTGAAAAAGAAAACAAATAAGTGTATTTATGCTATACATGATGAAACATC | GATGTTTCATCATGTATAGCATAAATACACTTATTTGTTTTCTTTTTCAAAGTGG |
| 121 | Y592C | c.1775A>G | CGCGTTGTCT | Unclassified | Class C65 | - | | 0.027 | Outside SPiCE Interpretation | CTTTGAAAAAGAAAACAAATAAGTTTATTTGTGCTATACATGATGAAACATCTTATAAAG | CTTTATAAGATGTTTCATCATGTATAGCACAAATAAACTTATTTGTTTTCTTTTTCAAAG |
| 122 | T1128I | c.3383C>T | TTCTCGGACC | Unclassified | Class C65 | - | | 0.027 | Outside SPiCE Interpretation | GAAGAATCAGGAAGTCAGTTTGAATTTATTCAGTTTAGAAAGCCAAGCTAC | GTAGCTTGGCTTTCTAAACTGAATAAATTCAAACTGACTTCCTGATTCTTC |
| 123 | G1224V | c.3671G>T | CCGTCTGTAT | Unclassified | Class C65 | - | | 0.027 | Outside SPiCE Interpretation | GTTTAGGGGCTTTTATTCTGCTCATGTCACAAAACTGAATGTTTCTACTG | CAGTAGAAACATTCAGTTTTGTGACATGAGCAGAATAAAAGCCCCTAAAC |
| 50-2 | S1970L | c.5909C>T | CTTTCGGACA | Unclassified | Class C0 | - | | 0.027 | Outside SPiCE Interpretation | GGAAGCTTCATAAGTCAGTCTTATCTGCAAATACTTGTGGGATTTTTAGC | GCTAAAAATCCCACAAGTATTTGCAGATAAGACTGACTTATGAAGCTTCC |
| 124 | T1980I | c.5939C>T | TCGTGATAGG | Unclassified | Class C65 | - | | 0.027 | Outside SPiCE Interpretation | CTGCAAATACTTGTGGGATTTTTAGCATAGCAAGTGGAAAATCTGTC | GACAGATTTTCCACTTGCTATGCTAAAAATCCCACAAGTATTTGCAG |
| 125 | S2006R | c.6016A>C | TTGTGCCATT | Unclassified | Class C0 | - | | 0.027 | Outside SPiCE Interpretation | CAAGTGTTTTCTGAAATAGAAGATCGTACCAAGCAAGTCTTTTCCAAAG | CTTTGGAAAAGACTTGCTTGGTACGATCTTCTATTTCAGAAAACACTTG |
| 126 | G2057E | c.6170G>A | GTGGTGGTGC | Unclassified | Class C65 | - | | 0.027 | Outside SPiCE Interpretation | GGTAAATTCATCTGCTTTCTCTGAATTTAGTACAGCAAGTGGAAAGC | GCTTTCCACTTGCTGTACTAAATTCAGAGAAAGCAGATGAATTTACC |
| 127 | R2488G | c.7462A>G | CCTCTCTTAG | Unclassified | Class C45 | DNA binding domain | | 0.027 | Outside SPiCE Interpretation | GATTTAATTACAAGTCTTCAGAATGCCGGAGATATACAGGATATGCGAATTAAG | CTTAATTCGCATATCCTGTATATCTCCGGCATTCTGAAGACTTGTAATTAAATC |
| 128 | Q2491R | c.7472A>G | ATCCCTGTTG | Unclassified | Class C35 | DNA binding domain | | 0.027 | Outside SPiCE Interpretation | CTTCAGAATGCCAGAGATATACGGGATATGCGAATTAAGAAGAAAC | GTTTCTTCTTAATTCGCATATCCCGTATATCTCTGGCATTCTGAAG |
| 129 | V2503D | c.7508T>A | CTCGCCGGTG | Unclassified | Class C45 | DNA binding domain | | 0.027 | Outside SPiCE Interpretation | GAAGAAACAAAGGCAACGCGACTTTCCACAGCCAGGCAGTCTG | CAGACTGCCTGGCTGTGGAAAGTCGCGTTGCCTTTGTTTCTTC |
| 130 | S2509R | c.7525A>C | TGCTGGAGGT | Unclassified | Class C65 | DNA binding domain | | 0.027 | Outside SPiCE Interpretation | CGTCTTTCCACAGCCAGGCCGTCTGTATCTTGCAAAAACATCC | GGATGTTTTTGCAAGATACAGACGGCCTGGCTGTGGAAAGACG |
| 131 | S2509N | c.7526G>A | TCGCGTGTAG | Unclassified | Class C45 | DNA binding domain | | 0.027 | Outside SPiCE Interpretation | CGTCTTTCCACAGCCAGGCAATCTGTATCTTGCAAAAACATCC | GGATGTTTTTGCAAGATACAGATTGCCTGGCTGTGGAAAGACG |
| 132 | L2510P | c.7529T>C | GAAATGATTG | Unclassified | Class C65 | DNA binding domain | | 0.027 | Outside SPiCE Interpretation | GTCTTTCCACAGCCAGGCAGTCCGTATCTTGCAAAAACATCCACTC | GAGTGGATGTTTTTGCAAGATACGGACTGCCTGGCTGTGGAAAGAC |
| 133 | K2514T | c.7541A>C | GTTGCCTTAT | Unclassified | Class C25 | DNA binding domain | | 0.027 | Outside SPiCE Interpretation | CAGGCAGTCTGTATCTTGCAACAACATCCACTCTGCCTCGAATC | GATTCGAGGCAGAGTGGATGTTGTTGCAAGATACAGACTGCCTG |
| 134 | T2515I | c.7544C>T | GGATTCGATT | Class 1 | Class C0 | DNA binding domain | | 0.027 | Outside SPiCE Interpretation | CAGGCAGTCTGTATCTTGCAAAAATATCCACTCTGCCTCGAATC | GATTCGAGGCAGAGTGGATATTTTTGCAAGATACAGACTGCCTG |
| 135 | S2516C | c.7547C>G | GTGTTTTTCC | Unclassified | Class C15 | DNA binding domain | | 0.027 | Outside SPiCE Interpretation | CAGTCTGTATCTTGCAAAAACATGCACTCTGCCTCGAATCTC | GAGATTCGAGGCAGAGTGCATGTTTTTGCAAGATACAGACTG |
| 136 | R2520Q | c.7559G>A | TGAGACTACG | Unclassified | Class C35 | DNA binding domain | | 0.027 | Outside SPiCE Interpretation | GCAAAAACATCCACTCTGCCTCAAATCTCTCTGAAAGCAGCAG | CTGCTGCTTTCAGAGAGATTTGAGGCAGAGTGGATGTTTTTGC |
| 137 | R2520P | c.7559G>C | GCATCAAGGT | Unclassified | Class C65 | DNA binding domain | | 0.027 | Outside SPiCE Interpretation | GCAAAAACATCCACTCTGCCTCCAATCTCTCTGAAAGCAGCAG | CTGCTGCTTTCAGAGAGATTGGAGGCAGAGTGGATGTTTTTGC |
| 138 | V2527I | c.7579G>A | CCATGTTCAC | Unclassified | Class C25 | DNA binding domain | | 0.027 | Outside SPiCE Interpretation | GAATCTCTCTGAAAGCAGCAATAGGAGGCCAAGTTCCCTCTG | CAGAGGGAACTTGGCCTCCTATTGCTGCTTTCAGAGAGATTC |
| 139 | P2532H | c.7595C>A | GTGATAGTTG | Unclassified | Class C65 | DNA binding domain | | 0.027 | Outside SPiCE Interpretation | CAGCAGTAGGAGGCCAAGTTCACTCTGCGTGTTCTCATAAACAG | CTGTTTATGAGAACACGCAGAGTGAACTTGGCCTCCTACTGCTG |
| 140 | G2544D | c.7631G>A | TCGCTTCCTT | Unclassified | Class C65 | DNA binding domain | | 0.027 | Outside SPiCE Interpretation | CTCATAAACAGCTGTATACGTATGACGTTTCTAAACATTGCATAAAAATTAACAG | CTGTTAATTTTTATGCAATGTTTAGAAACGTCATACGTATACAGCTGTTTATGAG |
| 141 | F2562V | c.7684T>G | CCTGTTTTCC | Unclassified | Class C45 | DNA binding domain | | 0.027 | Outside SPiCE Interpretation | CAGCAAAAATGCAGAGTCTTTTCAGGTTCACACTGAAGATTATTTTGGTAAG | CTTACCAAAATAATCTTCAGTGTGAACCTGAAAAGACTCTGCATTTTTGCTG |
| 142 | L2581W | c.7742T>G | CATTTAGTTG | Unclassified | Class C55 | DNA binding domain | | 0.027 | Outside SPiCE Interpretation | GGACTGGAAAAGGAATACAGTGGGCTGATGGCGGATGGCTC | GAGCCATCCGCCATCAGCCCACTGTATTCCTTTTCCAGTCC |
| 143 | G2584C | c.7750G>T | GCTTTTTGGT | Unclassified | Class C65 | DNA binding domain | | 0.027 | Outside SPiCE Interpretation | GAAAAGGAATACAGTTGGCTGATTGCGGATGGCTCATACCCTC | GAGGGTATGAGCCATCCGCAATCAGCCAACTGTATTCCTTTTC |
| 144 | K2597N | c.7791A>C | CTCTGCTGGG | Unclassified | Class C65 | DNA binding domain | | 0.027 | Outside SPiCE Interpretation | CAATGATGGAAAGGCTGGAAACGAAGAATTTTATAGGGCTCTGTG | CACAGAGCCCTATAAAATTCTTCGTTTCCAGCCTTTCCATCATTG |
| 145 | A2603T | c.7807G>A | GAAAGTCACT | Unclassified | Class C55 | DNA binding domain | | 0.023 | low | CTGGAAAAGAAGAATTTTATAGGACTCTGTGTGACACTCCAGGTGTG | CACACCTGGAGTGTCACACAGAGTCCTATAAAATTCTTCTTTTCCAG |
| 146 | T2607P | c.7819A>C | GGCGCTTTCC | Class 4 | Class C35 | DNA binding domain | | 0.027 | Outside SPiCE Interpretation | GAATTTTATAGGGCTCTGTGTGACCCTCCAGGTGTGGATCCAAAGC | GCTTTGGATCCACACCTGGAGGGTCACACAGAGCCCTATAAAATTC |
| 147 | D2611G | c.7832A>G | CGTGTTGTTA | Unclassified | Class C65 | DNA binding domain | | 0.027 | Outside SPiCE Interpretation | GTGTGACACTCCAGGTGTGGGTCCAAAGCTTATTTCTAGAATTTGG | CCAAATTCTAGAAATAAGCTTTGGACCCACACCTGGAGTGTCACAC |
| 148 | W2619C | c.7857G>C | TGTGTGGTCG | Unclassified | Class C65 | DNA binding domain | | 0.027 | Outside SPiCE Interpretation | GATCCAAAGCTTATTTCTAGAATTTGCGTTTATAATCACTATAGATGGATC | GATCCATCTATAGTGATTATAAACGCAAATTCTAGAAATAAGCTTTGGATC |
| 149 | I2627F | c.7879A>T | TTTCGTTGAT | Class 5 | Class C15 | DNA binding domain | | 0.027 | Outside SPiCE Interpretation | GGTTTATAATCACTATAGATGGTTCATATGGAAACTGGCAGCTATG | CATAGCTGCCAGTTTCCATATGAACCATCTATAGTGATTATAAACC |
| 150 | F2642C | c.7925T>G | TGGACTGTAC | Unclassified | Class C45 | DNA binding domain | | 0.027 | Outside SPiCE Interpretation | GGAATGTGCCTTTCCTAAGGAATGTGCTAATAGATGCCTAAGC | GCTTAGGCATCTATTAGCACATTCCTTAGGAAAGGCACATTCC |
| 151 | L2647P | c.7940T>C | CATGCTGTGG | Class 4 | Class C65 | DNA binding domain | | 0.027 | Outside SPiCE Interpretation | GGAATTTGCTAATAGATGCCCAAGCCCAGAAAGGGTGCTTC | GAAGCACCCTTTCTGGGCTTGGGCATCTATTAGCAAATTCC |
| 152 | V2652G | c.7955T>G | CGGTTGAACA | Unclassified | Class C65 | DNA binding domain | | 0.027 | Outside SPiCE Interpretation | GATGCCTAAGCCCAGAAAGGGGGCTTCTTCAACTAAAATACAG | CTGTATTTTAGTTGAAGAAGCCCCCTTTCTGGGCTTAGGCATC |
| 153 | L2653P | c.7958T>C | ACCCCGCTAT | Class 5 | Class C65 | DNA binding domain | | 0.027 | Outside SPiCE Interpretation | GATGCCTAAGCCCAGAAAGGGTGCCTCTTCAACTAAAATACAGATATG | CATATCTGTATTTTAGTTGAAGAGGCACCCTTTCTGGGCTTAGGCATC |
| 154 | L2654P | c.7961T>C | CGTATCGGCT | Unclassified | Class C65 | DNA binding domain | | 0.027 | Outside SPiCE Interpretation | CTAAGCCCAGAAAGGGTGCTTCCTCAACTAAAATACAGATATG | CATATCTGTATTTTAGTTGAGGAAGCACCCTTTCTGGGCTTAG |
| 155 | Y2660D | c.7978T>G | GCGATCGCGC | Unclassified | Class C65 | DNA binding domain | | 0.054 | low | GTGCTTCTTCAACTAAAATACAGAGATGATACGGAAATTGATAGAAGC | GCTTCTATCAATTTCCGTATCATCTCTGTATTTTAGTTGAAGAAGCAC |
| 156 | S2670L | c.8009C>T | GTGCTCTGTA | Class 4 | Class C15 | DNA binding domain | | 0.027 | Outside SPiCE Interpretation | GATACGGAAATTGATAGAAGCAGAAGATTGGCTATAAAAAAGATAATGGAAAGG | CCTTTCCATTATCTTTTTTATAGCCAATCTTCTGCTTCTATCAATTTCCGTATC |
| 157 | S2670W | c.8009C>G | ATGGCCGGGC | Unclassified | Class C25 | DNA binding domain | | 0.027 | Outside SPiCE Interpretation | GATACGGAAATTGATAGAAGCAGAAGATGGGCTATAAAAAAGATAATGG | CCATTATCTTTTTTATAGCCCATCTTCTGCTTCTATCAATTTCCGTATC |
| 158 | K2673N | c.8019A>T | CACGAGTGGT | Unclassified | Class C35 | DNA binding domain | | 0.027 | Outside SPiCE Interpretation | GATAGAAGCAGAAGATCGGCTATAAATAAGATAATGGAAAGGGATGAC | GTCATCCCTTTCCATTATCTTATTTATAGCCGATCTTCTGCTTCTATC |
| 159 | R2678G | c.8032A>G | GGGCGGGTGA | Unclassified | Class C45 | DNA binding domain | | 0.027 | Outside SPiCE Interpretation | GATCGGCTATAAAAAAGATAATGGAAGGGGATGACACAGCTGCAAAAACAC | GTGTTTTTGCAGCTGTGTCATCCCCTTCCATTATCTTTTTTATAGCCGATC |
| 160 | A2682V | c.8045C>T | TACTGTTCTA | Unclassified | Class C25 | DNA binding domain | | 0.027 | Outside SPiCE Interpretation | GATAATGGAAAGGGATGACACAGTTGCAAAAACACTTGTTCTCTGTG | CACAGAGAACAAGTGTTTTTGCAACTGTGTCATCCCTTTCCATTATC |
| 161 | L2686P | c.8057T>C | TTTTTTTTCC | Unclassified | Class C45 | DNA binding domain | | 0.027 | Outside SPiCE Interpretation | GATGACACAGCTGCAAAAACACCTGTTCTCTGTGTTTCTGAC | GTCAGAAACACAGAGAACAGGTGTTTTTGCAGCTGTGTCATC |
| 162 | A2730V | c.8189C>T | TTTCTTTGTG | Unclassified | Class C0 | DNA binding domain | | 0.027 | Outside SPiCE Interpretation | GATGGGTGGTATGCTGTTAAGGTCCAGTTAGATCCTCCCCTCTTAG | CTAAGAGGGGAGGATCTAACTGGACCTTAACAGCATACCACCCATC |
| 163 | R2744G | c.8230A>G | CTTTGCTCTC | Unclassified | Class C45 | DNA binding domain | | 0.027 | Outside SPiCE Interpretation | CTTAGCTGTCTTAAAGAATGGCGGACTGACAGTTGGTCAGAAG | CTTCTGACCAACTGTCAGTCCGCCATTCTTTAAGACAGCTAAG |
| 164 | L2745V | c.8233C>G | GGTGCCTTGT | Unclassified | Class C15 | DNA binding domain | | 0.027 | Outside SPiCE Interpretation | CTTAGCTGTCTTAAAGAATGGCAGAGTGACAGTTGGTCAGAAG | CTTCTGACCAACTGTCACTCTGCCATTCTTTAAGACAGCTAAG |
| 165 | G2755V | c.8264G>T | TGCTCGGGTA | Unclassified | Class C65 | DNA binding domain | | 0.027 | Outside SPiCE Interpretation | GTCAGAAGATTATTCTTCATGTAGCAGAACTGGTGGGCTCTC | GAGAGCCCACCAGTTCTGCTACATGAAGAATAATCTTCTGAC |
| 166 | A2764V | c.8291C>T | TGAGTGCGTG | Unclassified | Class C15 | DNA binding domain | | 0.027 | Outside SPiCE Interpretation | CTGGTGGGCTCTCCTGATGTCTGTACACCTCTTGAAGC | GCTTCAAGAGGTGTACAGACATCAGGAGAGCCCACCAG |
| 167 | P2771L | c.8312C>T | GCATGGGGAG | Unclassified | Class C65 | DNA binding domain | | 0.027 | Outside SPiCE Interpretation | CTGTACACCTCTTGAAGCCCTAGAATCTCTTATGTTAAAGATTTC | GAAATCTTTAACATAAGAGATTCTAGGGCTTCAAGAGGTGTACAG |
| 168 | S2773C | c.8318C>G | GTATGGCTGG | Unclassified | Class C25 | DNA binding domain | | 0.027 | Outside SPiCE Interpretation | GTACACCTCTTGAAGCCCCAGAATGTCTTATGTTAAAGATTTCTG | CAGAAATCTTTAACATAAGACATTCTGGGGCTTCAAGAGGTGTAC |
| 169 | K2777E | c.8329A>G | ACCTTTGGTG | Unclassified | Class C55 | DNA binding domain | | 0.109 | low | GAAGCCCCAGAATCTCTTATGTTAGAGATTTCTGCTAACAGTACTC | GAGTACTGTTAGCAGAAATCTCTAACATAAGAGATTCTGGGGCTTC |
| 170 | R2784W | c.8350C>T | CCGTCCTTGG | Unclassified | Class C65 | DNA binding domain | | 0.027 | Outside SPiCE Interpretation | GTTAAAGATTTCTGCTAACAGTACTTGGCCTGCTCGCTGGTATAC | GTATACCAGCGAGCAGGCCAAGTACTGTTAGCAGAAATCTTTAAC |
| 171 | R2784Q | c.8351G>A | GACGGGCTAT | Class 1 | Class C35 | DNA binding domain | | 0.027 | Outside SPiCE Interpretation | GATTTCTGCTAACAGTACTCAGCCTGCTCGCTGGTATACC | GGTATACCAGCGAGCAGGCTGAGTACTGTTAGCAGAAATC |
| 172 | W2788C | c.8364G>C | GGTTTTGGTG | Unclassified | Class C45 | DNA binding domain | | 0.027 | Outside SPiCE Interpretation | CAGTACTCGGCCTGCTCGCTGCTATACCAAACTTGGGTTC | GAACCCAAGTTTGGTATAGCAGCGAGCAGGCCGAGTACTG |
| 173 | L2792P | c.8375T>C | TGTTAGTGGC | Unclassified | Class C65 | DNA binding domain | | 0.027 | Outside SPiCE Interpretation | CTGCTCGCTGGTATACCAAACCTGGGTTCTTTCCTGACCCTAG | CTAGGGTCAGGAAAGAACCCAGGTTTGGTATACCAGCGAGCAG |
| 174 | G2793V | c.8378G>T | GCTCGCCCTG | Unclassified | Class C65 | DNA binding domain | | 0.027 | Outside SPiCE Interpretation | CTCGCTGGTATACCAAACTTGTGTTCTTTCCTGACCCTAGAC | GTCTAGGGTCAGGAAAGAACACAAGTTTGGTATACCAGCGAG |
| 175 | P2800T | c.8398C>A | GATAGTGGTG | Unclassified | Class C35 | DNA binding domain | | 0.027 | Outside SPiCE Interpretation | GGTTCTTTCCTGACCCTAGAACTTTTCCTCTGCCCTTATCATC | GATGATAAGGGCAGAGGAAAAGTTCTAGGGTCAGGAAAGAACC |
| 176 | P2800R | c.8399C>G | CATTGTTGCG | Unclassified | Class C65 | DNA binding domain | | 0.027 | Outside SPiCE Interpretation | GTTCTTTCCTGACCCTAGACGTTTTCCTCTGCCCTTATCATC | GATGATAAGGGCAGAGGAAAACGTCTAGGGTCAGGAAAGAAC |
| 177 | G2813E | c.8438G>A | GTACTTTGGG | Unclassified | Class C65 | DNA binding domain | | 0.027 | Outside SPiCE Interpretation | CATCGCTTTTCAGTGATGGAGAAAATGTTGGTTGTGTTGATG | CATCAACACAACCAACATTTTCTCCATCACTGAAAAGCGATG |
| 178 | Y2826C | c.8477A>G | TGATGCGACT | Unclassified | Class C65 | DNA binding domain | | 0.027 | Outside SPiCE Interpretation | GTGTTGATGTAATTATTCAAAGAGCATGCCCTATACAGTGGATGGAG | CTCCATCCACTGTATAGGGCATGCTCTTTGAATAATTACATCAACAC |
| 179 | P2827A | c.8479C>G | CACGCGGGCT | Unclassified | Class C25 | DNA binding domain | | 0.027 | Outside SPiCE Interpretation | GTAATTATTCAAAGAGCATACGCTATACAGTGGATGGAGAAGAC | GTCTTCTCCATCCACTGTATAGCGTATGCTCTTTGAATAATTAC |
| 180 | K2833N | c.8499G>C | GGTCTCTTTC | Unclassified | Class C35 | DNA binding domain | | 0.027 | Outside SPiCE Interpretation | CATACCCTATACAGTGGATGGAGAACACATCATCTGGATTATACATATTTC | GAAATATGTATAATCCAGATGATGTGTTCTCCATCCACTGTATAGGGTATG |
| 181 | R2842C | c.8524C>T | CCGATGCAAT | Unclassified | Class C65 | DNA binding domain | | 0.027 | Outside SPiCE Interpretation | GAAGACATCATCTGGATTATACATATTTTGCAATGAAAGAGAGGAAG | CTTCCTCTCTTTCATTGCAAAATATGTATAATCCAGATGATGTCTTC |
| 182 | R2842L | c.8525G>T | GTTTTTGGTA | Unclassified | Class C65 | DNA binding domain | | 0.027 | Outside SPiCE Interpretation | CATCATCTGGATTATACATATTTCTCAATGAAAGAGAGGAAGAAAAGG | CCTTTTCTTCCTCTCTTTCATTGAGAAATATGTATAATCCAGATGATG |
| 183 | E2847K | c.8539G>A | TGTATATTTC | Unclassified | Class C55 | DNA binding domain | | 0.027 | Outside SPiCE Interpretation | CATATTTCGCAATGAAAGAGAGAAAGAAAAGGAAGCAGCAAAATATG | CATATTTTGCTGCTTCCTTTTCTTTCTCTCTTTCATTGCGAAATATG |
| 184 | L2862Q | c.8585T>A | ATCTGTCATA | Unclassified | Class C55 | DNA binding domain | | 0.027 | Outside SPiCE Interpretation | GTGGAGGCCCAACAAAAGAGACAAGAAGCCTTATTCACTAAAATTC | GAATTTTAGTGAATAAGGCTTCTTGTCTCTTTTGTTGGGCCTCCAC |
| 185 | L2865V | c.8593T>G | GTTTGTTTTT | Unclassified | Class C25 | DNA binding domain | | 0.027 | Outside SPiCE Interpretation | CAACAAAAGAGACTAGAAGCCGTATTCACTAAAATTCAGGAGG | CCTCCTGAATTTTAGTGAATACGGCTTCTAGTCTCTTTTGTTG |
| 186 | I2869N | c.8606T>A | TTTGTCATTG | Unclassified | Class C45 | DNA binding domain | | 0.027 | Outside SPiCE Interpretation | CAAAAGAGACTAGAAGCCTTATTCACTAAAAATCAGGAGGAATTTGAAGAAC | GTTCTTCAAATTCCTCCTGATTTTTAGTGAATAAGGCTTCTAGTCTCTTTTG |
| 187 | F2873C | c.8618T>G | GTACGGGGCT | Unclassified | Class C55 | DNA binding domain | | 0.027 | Outside SPiCE Interpretation | CTTATTCACTAAAATTCAGGAGGAATGTGAAGAACATGAAGAAAACACAAC | GTTGTGTTTTCTTCATGTTCTTCACATTCCTCCTGAATTTTAGTGAATAAG |
| 188 | D2900V | c.8699A>T | TCTTTTGTGA | Unclassified | Class C25 | DNA binding domain | | 0.027 | Outside SPiCE Interpretation | CAGCAAGTTCGTGCTTTGCAAGTTGGTGCAGAGCTTTATGAAGC | GCTTCATAAAGCTCTGCACCAACTTGCAAAGCACGAACTTGCTG |
| 189 | G2901V | c.8702G>T | TAATAGATGG | Unclassified | Class C65 | DNA binding domain | | 0.027 | Outside SPiCE Interpretation | CAAGTTCGTGCTTTGCAAGATGTTGCAGAGCTTTATGAAGCAGTG | CACTGCTTCATAAAGCTCTGCAACATCTTGCAAAGCACGAACTTG |
| 190 | Y2905H | c.8713T>C | GTTGCAGTCT | Unclassified | Class C35 | DNA binding domain | | 0.027 | Outside SPiCE Interpretation | CTTTGCAAGATGGTGCAGAGCTTCATGAAGCAGTGAAGAATGCAG | CTGCATTCTTCACTGCTTCATGAAGCTCTGCACCATCTTGCAAAG |
| 191 | Y2905C | c.8714A>G | GTTGTTTCAG | Unclassified | Class C55 | DNA binding domain | | 0.027 | Outside SPiCE Interpretation | CTTTGCAAGATGGTGCAGAGCTTTGTGAAGCAGTGAAGAATGCAG | CTGCATTCTTCACTGCTTCACAAAGCTCTGCACCATCTTGCAAAG |
| 192 | D2913V | c.8738A>T | TGTCGTCGCT | Unclassified | Class C65 | DNA binding domain | | 0.027 | Outside SPiCE Interpretation | GAAGCAGTGAAGAATGCAGCAGTCCCAGCTTACCTTGAGGGTTATTTC | GAAATAACCCTCAAGGTAAGCTGGGACTGCTGCATTCTTCACTGCTTC |
| 193 | S2922R | c.8764A>C | TCCATTGGTC | Unclassified | Class C15 | DNA binding domain | | 0.027 | Outside SPiCE Interpretation | GCTTACCTTGAGGGTTATTTCCGTGAAGAGCAGTTAAGAGCCTTG | CAAGGCTCTTAACTGCTCTTCACGGAAATAACCCTCAAGGTAAGC |
| 194 | Q2925R | c.8774A>G | CGTATCTACA | Unclassified | Class C35 | DNA binding domain | | 0.027 | Outside SPiCE Interpretation | GAGGGTTATTTCAGTGAAGAGCGGTTAAGAGCCTTGAATAATCAC | GTGATTATTCAAGGCTCTTAACCGCTCTTCACTGAAATAACCCTC |
| 195 | Q2925H | c.8775G>C | CTTGTATCTA | Unclassified | Class C15 | DNA binding domain | | 0.027 | Outside SPiCE Interpretation | GAGGGTTATTTCAGTGAAGAGCACTTAAGAGCCTTGAATAATCACAG | CTGTGATTATTCAAGGCTCTTAAGTGCTCTTCACTGAAATAACCCTC |
| 196 | R2933T | c.8798G>C | AAGTGGGGGC | Unclassified | Class C25 | DNA binding domain | | 0.027 | Outside SPiCE Interpretation | GCAGTTAAGAGCCTTGAATAATCACACGCAAATGTTGAATGATAAG | CTTATCATTCAACATTTGCGTGTGATTATTCAAGGCTCTTAACTGC |
| 197 | V2966A | c.8897T>C | GTGGCTAGAC | Unclassified | Class C65 | DNA binding domain | | 0.027 | Outside SPiCE Interpretation | GAACAAGGTTTATCAAGGGATGCCACAACCGTGTGGAAGTTGC | GCAACTTCCACACGGTTGTGGCATCCCTTGATAAACCTTGTTC |
| 198 | L2972W | c.8915T>G | TGGGTTGGTT | Unclassified | Class C15 | DNA binding domain | | 0.027 | Outside SPiCE Interpretation | GATGTCACAACCGTGTGGAAGTGGCGTATTGTAAGCTATTC | GAATAGCTTACAATACGCCACTTCCACACGGTTGTGACATC |
| 199 | S2998Y | c.8993C>A | CGCAGACCGG | Unclassified | Class C15 | DNA binding domain | | 0.027 | Outside SPiCE Interpretation | GAGTATTTGGCGTCCATCATCAGATTTATATTATCTGTTAACAGAAGGAAAGAG | CTCTTTCCTTCTGTTAACAGATAATATAAATCTGATGATGGACGCCAAATACTC |
| 200 | E3002D | c.9006A>T | AGTTCGTGTG | Unclassified | Class C35 | DNA binding domain | | 0.027 | Outside SPiCE Interpretation | CATCAGATTTATATTCTCTGTTAACAGATGGAAAGAGATACAGAATTTATC | GATAAATTCTGTATCTCTTTCCATCTGTTAACAGAGAATATAAATCTGATG |
| 201 | Y3006D | c.9016T>G | CGCGTGCCTG | Unclassified | Class C55 | DNA binding domain | | 0.027 | Outside SPiCE Interpretation | GATTTATATTCTCTGTTAACAGAAGGAAAGAGAGACAGAATTTATCATCTTGCAAC | GTTGCAAGATGATAAATTCTGTCTCTCTTTCCTTCTGTTAACAGAGAATATAAATC |
| 202 | R3007G | c.9019A>G | TCTTCTGGGG | Unclassified | Class C45 | DNA binding domain | | 0.027 | Outside SPiCE Interpretation | CTGTTAACAGAAGGAAAGAGATACGGAATTTATCATCTTGCAACTTC | GAAGTTGCAAGATGATAAATTCCGTATCTCTTTCCTTCTGTTAACAG |
| 203 | L3011P | c.9032T>C | GTATTGAGGG | Unclassified | Class C25 | DNA binding domain | | 0.027 | Outside SPiCE Interpretation | GAAGGAAAGAGATACAGAATTTATCATCCTGCAACTTCAAAATCTAAAAGTAAATC | GATTTACTTTTAGATTTTGAAGTTGCAGGATGATAAATTCTGTATCTCTTTCCTTC |
| 204 | T3030I | c.9089C>T | TTGCAGCTTG | Unclassified | Class C65 | DNA binding domain | | 0.027 | Outside SPiCE Interpretation | GAGCTAACATACAGTTAGCAGCGATAAAAAAAACTCAGTATCAACAAC | GTTGTTGATACTGAGTTTTTTTTATCGCTGCTAACTGTATGTTAGCTC |
| 205 | T3033I | c.9098C>T | TTCTGCCATG | Unclassified | Class C65 | DNA binding domain | | 0.027 | Outside SPiCE Interpretation | CATACAGTTAGCAGCGACAAAAAAAATTCAGTATCAACAACTACCGGTTTC | GAAACCGGTAGTTGTTGATACTGAATTTTTTTTGTCGCTGCTAACTGTATG |
| 206 | Y3035C | c.9104A>G | TTAGGGGAAA | Unclassified | Class C55 | DNA binding domain | | 0.027 | Outside SPiCE Interpretation | CAGTTAGCAGCGACAAAAAAAACTCAGTGTCAACAACTACCGGTTTCAG | CTGAAACCGGTAGTTGTTGACACTGAGTTTTTTTTGTCGCTGCTAACTG |
| 207 | F3060C | c.9179T>G | CGGATAGGCC | Unclassified | Class C55 | DNA binding domain | | 0.027 | Outside SPiCE Interpretation | GAGCCCCTTCACTTCAGCAAATGTTTAGATCCAGACTTTCAGC | GCTGAAAGTCTGGATCTAAACATTTGCTGAAGTGAAGGGGCTC |
| 208 | V3072E | c.9215T>A | TACTTCGCGC | Unclassified | Class C35 | DNA binding domain | | 0.027 | Outside SPiCE Interpretation | CAGCCATCTTGTTCTGAGGAGGACCTAATAGGATTTGTC | GACAAATCCTATTAGGTCCTCCTCAGAACAAGATGGCTG |
| 209 | G3076E | c.9227G>A | TCCCTTTTTG | Unclassified | Class C65 | DNA binding domain | | 0.027 | Outside SPiCE Interpretation | GTTCTGAGGTGGACCTAATAGAATTTGTCGTTTCTGTTGTG | CACAACAGAAACGACAAATTCTATTAGGTCCACCTCAGAAC |
| 210 | L3101R | c.9302T>G | CTGCCTCACG | Unclassified | Class C35 | DNA binding domain | | 0.027 | Outside SPiCE Interpretation | GTCAGACGAATGTTACAATTTACGGGCAATAAAGTTTTGGATAGAC | GTCTATCCAAAACTTTATTGCCCGTAAATTGTAACATTCGTCTGAC |
| 211 | L3101P | c.9302T>C | TCGGACCTGC | Unclassified | Class C25 | DNA binding domain | | 0.027 | Outside SPiCE Interpretation | GTCAGACGAATGTTACAATTTACCGGCAATAAAGTTTTGGATAGAC | GTCTATCCAAAACTTTATTGCCGGTAAATTGTAACATTCGTCTGAC |
| 212 | S3123G | c.9367A>G | ACTTTCCGTG | Unclassified | Class C55 | DNA binding domain | | 0.027 | Outside SPiCE Interpretation | CCTCATATGTTAATTGCTGCAGGCAACCTCCAGTGGCGACCAG | CTGGTCGCCACTGGAGGTTGCCTGCAGCAATTAACATATGAGG |
| 213 | S3123R | c.9369C>A | GCAGCTATGC | Unclassified | Class C65 | DNA binding domain | | 0.027 | Outside SPiCE Interpretation | CATATGTTAATTGCTGCAAGAAACCTCCAGTGGCGACCAGAATC | GATTCTGGTCGCCACTGGAGGTTTCTTGCAGCAATTAACATATG |
| 214 | Q3126P | c.9377A>C | TTGTCACGAC | Unclassified | Class C15 | DNA binding domain | | 0.027 | Outside SPiCE Interpretation | GTTAATTGCTGCAAGCAACCTCCCGTGGCGACCAGAATCCAAATC | GATTTGGATTCTGGTCGCCACGGGAGGTTGCTTGCAGCAATTAAC |
| 215 | R3269G | c.9805A>G | ACCTGGTCTG | Unclassified | Class C45 | TR2 RAD51 binding domain | | 0.027 | Outside SPiCE Interpretation | CCAAAAGAACTGCAAAAAGAGAGGAGCCTTGGATTTCTTGAGTAG | CTACTCAAGAAATCCAAGGCTCCTCTCTTTTTGCAGTTCTTTTGG |
| 216 | R3269T | c.9806G>C | TAGGGGCGTC | Unclassified | Class C25 | TR2 RAD52 binding domain | | 0.027 | Outside SPiCE Interpretation | GACCAAAAGAACTGCAAAAAGAGAACAGCCTTGGATTTCTTGAG | CTCAAGAAATCCAAGGCTGTTCTCTTTTTGCAGTTCTTTTGGTC |
| 217 | L3274W | c.9821T>G | CTGCGTTTTT | Unclassified | Class C55 | TR2 RAD53 binding domain | | 0.027 | Outside SPiCE Interpretation | GAGAAGAGCCTTGGATTTCTGGAGTAGACTGCCTTTACCTCCAC | GTGGAGGTAAAGGCAGTCTACTCCAGAAATCCAAGGCTCTTCTC |
| 218 | P3280R | c.9839C>G | TCATGTTCTG | Unclassified | Class C65 | TR2 RAD54 binding domain | | 0.027 | Outside SPiCE Interpretation | CTTGAGTAGACTGCCTTTACGTCCACCTGTTAGTCCCATTTG | CAAATGGGACTAACAGGTGGACGTAAAGGCAGTCTACTCAAG |
| 219 | I3286N | c.9857T>A | TGTGTCTAGC | Unclassified | Class C35 | TR2 RAD55 binding domain | | 0.027 | Outside SPiCE Interpretation | CCTCCACCTGTTAGTCCCAATTGTACATTTGTTTCTCC | GGAGAAACAAATGTACAATTGGGACTAACAGGTGGAGG |
| 220 | Q3295P | c.9884A>C | GCTGGCGGGG | Unclassified | Class C15 | TR2 RAD56 binding domain | | 0.027 | Outside SPiCE Interpretation | CATTTGTTTCTCCGGCTGCACCGAAGGCATTTCAGCCACCAAG | CTTGGTGGCTGAAATGCCTTCGGTGCAGCCGGAGAAACAAATG |
| 221 | P3301L | c.9902C>T | CTCGATCTCT | Unclassified | Class C65 | TR2 RAD57 binding domain | | 0.027 | Outside SPiCE Interpretation | CAGAAGGCATTTCAGCCACTAAGGAGTTGTGGCACCAAATAC | GTATTTGGTGCCACAACTCCTTAGTGGCTGAAATGCCTTCTG |
| 106-2 | S3319F | c.9956C>T | GTGATGCTAG | Unclassified | Class C0 | - | | 0.027 | Outside SPiCE Interpretation | GAAACACCCATAAAGAAAAAAGAACTGAATTTTCCTCAGATGACTCC | GGAGTCATCTGAGGAAAATTCAGTTCTTTTTTCTTTATGGGTGTTTC |
| 222 | Y3035S | c.9104A>C | ATTCTTGTTG | Class 2 | Class C55 | DNA binding domain | | 0.027 | Outside SPiCE Interpretation | GCAGCGACAAAAAAAACTCAGTCTCAACAACTACCGGTTTCAG | CTGAAACCGGTAGTTGTTGAGACTGAGTTTTTTTTGTCGCTGC |

Supplementary Table 1: Reclassifications of IARC Class 1/2/4/5 BRCA2 variants by ClinVar, ACMG, and MANO-B method.

| Variant | IARC Classification | ClinVar classification | ACMG clasification | MANO-B classification | Type of evidence | ACMG edidence: PVS1 | PS1 | PS2 | PS3 | PS4 | PM1 | PM2 | PM3 | PM4 | PM5 | PM6 | PP1 | PP2 | PP3 | PP4 | PP5 | BA1 | BS1 | BS2 | BS3 | BS4 | BP1 | BP2 | BP3 | BP4 | BP5 | BP6 | BP7 |
|---|---|---|---|---|---|---|---|---|---|---|---|---|---|---|---|---|---|---|---|---|---|---|---|---|---|---|---|---|---|---|---|---|---|
| R18H | Class 2 | Benign | Benign | fClass 3 | BS3, BS4, BP4, BP6 | - | - | - | - | - | - | - | - | - | - | - | - | - | - | - | - | - | - | - | + | + | - | - | - | + | - | + | - |
| R18C | Class 2 | VUS | Likely benign | fClass 1 | PM2, BS4, BP2 | - | - | - | - | - | - | + | - | - | - | - | - | - | - | - | - | - | - | - | - | + | - | + | - | - | - | - | - |
| Y42C | Class 1 | Benign | Benign | fClass 1 | BS1, BS3, BS4, BP2, BP6 | - | - | - | - | - | - | - | - | - | - | - | - | - | - | - | - | - | + | - | + | + | - | + | - | - | - | + | - |
| P168T | Class 1 | Benign | Likely benign | fClass 1 | BS3, BP6 | - | - | - | - | - | - | - | - | - | - | - | - | - | - | - | - | - | - | - | + | - | - | - | - | - | - | + | - |
| S326R | Class 1 | Benign | Likely benign | fClass 1 | BP2, BP6 | - | - | - | - | - | - | - | - | - | - | - | - | - | - | - | - | - | - | - | - | - | - | + | - | - | - | + | - |
| E462G | Class 1 | Benign | Benign | fClass 1 | BS3, BS4, BP6 | - | - | - | - | - | - | - | - | - | - | - | - | - | - | - | - | - | - | - | + | + | - | - | - | - | - | + | - |
| N588D | Class 2 | Likely benign | Uncertain significance | fClass 1 | - | - | - | - | - | - | - | - | - | - | - | - | - | - | - | - | - | - | - | - | - | - | - | - | - | - | - | - | - |
| T598A | Class 2 | Benign/Likely benign | Benign | fClass 1 | BS1, BS2, BS4, BP6 | - | - | - | - | - | - | - | - | - | - | - | - | - | - | - | - | - | + | - | + | + | - | - | - | - | - | + | - |
| G602R | Class 1 | Benign | Likely benign | fClass 1 | BS4, BP6 | - | - | - | - | - | - | - | - | - | - | - | - | - | - | - | - | - | - | - | - | + | - | - | - | - | - | + | - |
| L1019V | Class 1 | Benign | Likely benign | fClass 2 | BS4, BP4, BP6 | - | - | - | - | - | - | - | - | - | - | - | - | - | - | - | - | - | - | - | - | + | - | - | - | + | - | + | - |
| S1172L | Class 1 | Benign | Likely benign | fClass 2 | BS1, BP6 | - | - | - | - | - | - | - | - | - | - | - | - | - | - | - | - | - | + | - | + | - | - | - | - | - | - | + | - |
| N1228D | Class 1 | Benign | Likely benign | fClass 2 | BS4, BP6 | - | - | - | - | - | - | - | - | - | - | - | - | - | - | - | - | - | - | - | - | + | - | - | - | - | - | + | - |
| D1420Y | Class 1 | Benign | Likely benign | fClass 1 | BS1, BP2, BP6 | - | - | - | - | - | - | - | - | - | - | - | - | - | - | - | - | - | + | - | - | - | - | + | - | - | - | + | - |
| F1524V | Class 1 | Benign | Uncertain significance | fClass 1 | PP3, BP6 | - | - | - | - | - | - | - | - | - | - | - | - | - | + | - | - | - | - | - | - | - | - | - | - | - | - | + | - |
| K1690N | Class 1 | Benign | Uncertain significance | fClass 1 | BP6 | - | - | - | - | - | - | - | - | - | - | - | - | - | - | - | - | - | - | - | - | - | - | - | - | - | - | + | - |
| N1878K | Class 1 | Conflicting | Uncertain significance | fClass 1 | - | - | - | - | - | - | - | - | - | - | - | - | - | - | - | - | - | - | - | - | - | - | - | - | - | - | - | - | - |
| H1918R | Class 1 | Conflicting | Uncertain significance | fClass 3 | BP2 | - | - | - | - | - | - | - | - | - | - | - | - | - | - | - | - | - | - | - | - | - | - | + | - | - | - | - | - |
| I1929V | Class 1 | Benign | Benign | fClass 1 | BS1, BS4, BP2, BP4, BP6 | - | - | - | - | - | - | - | - | - | - | - | - | - | - | - | - | - | + | - | - | + | - | + | - | + | - | + | - |
| E2020K | Class 2 | VUS | Uncertain significance | fClass 2 | PM2, BP4 | - | - | - | - | - | - | + | - | - | - | - | - | - | - | - | - | - | - | - | - | - | - | - | - | + | - | - | - |
| H2074N | Class 1 | Benign | Likely benign | fClass 1 | BP4, BP6 | - | - | - | - | - | - | - | - | - | - | - | - | - | - | - | - | - | - | - | - | - | - | - | - | + | - | + | - |
| K2411T | Class 1 | Conflicting | Uncertain significance | fClass 1 | BS3 | - | - | - | - | - | - | - | - | - | - | - | - | - | - | - | - | - | - | - | + | - | - | - | - | - | - | - | - |
| R2418G | Class 2 | Conflicting | Uncertain significance | fClass 3 | PM2, BP4 | - | - | - | - | - | - | + | - | - | - | - | - | - | - | - | - | - | - | - | - | - | - | - | - | + | - | - | - |
| N2436I | Class 1 | Benign | Likely benign | fClass 3 | BP2, BP4, BP6 | - | - | - | - | - | - | - | - | - | - | - | - | - | - | - | - | - | - | - | + | - | - | + | - | + | - | + | - |
| G2508S | Class 4 | Conflicting | Uncertain significance | fClass 4 | PP3, BS3 | - | - | - | - | - | - | - | - | - | - | - | - | - | + | - | - | - | - | - | + | - | - | - | - | - | - | - | - |
| T2515I | Class 1 | Benign | Likely benign | fClass 2 | BS3, BP2, BP6 | - | - | - | - | - | - | - | - | - | - | - | - | - | - | - | - | - | - | - | + | - | - | + | - | - | - | + | - |
| T2607P | Class 4 | VUS | Uncertain significance | fClass 5 | PM2 | - | - | - | - | - | - | + | - | - | - | - | - | - | - | - | - | - | - | - | - | - | - | - | - | - | - | - | - |
| G2609D | Class 4 | Likely pathogenic | Pathogenic | fClass 5 | PS3, PM2, PP3 | - | - | - | + | - | - | + | - | - | - | - | - | - | + | - | - | - | - | - | - | - | - | - | - | - | - | - | - |
| W2626C | Class 5 | Pathogenic | Pathogenic | fClass 5 | PS3, PM2, PP3, PP5 | - | - | - | + | - | - | - | - | - | - | - | - | - | + | - | + | - | - | - | - | - | - | - | - | - | - | - | - |
| I2627F | Class 5 | Pathogenic | Likely pathogenic | fClass 5 | PS3, PP3, PP5 | - | - | - | + | - | - | + | - | - | - | - | - | - | + | - | + | - | - | - | - | - | - | - | - | - | - | - | - |
| L2647P | Class 4 | Conflicting | Pathogenic | fClass 5 | PS3, PM2, PP3 | - | - | - | + | - | - | + | - | - | - | - | - | - | + | - | - | - | - | - | - | - | - | - | - | - | - | - | - |
| L2653P | Class 5 | Pathogenic | Pathogenic | fClass 5 | PS3, PM2, PP5 | - | - | - | + | - | - | + | - | - | - | - | - | - | - | - | + | - | - | - | - | - | - | - | - | - | - | - | - |
| R2659G | Class 5 | Conflicting | Likely pathogenic | fClass 5 | PS3, PM2, PP1, PP3 | - | - | - | + | - | - | + | - | - | - | - | + | - | + | - | - | - | - | - | - | - | - | - | - | - | - | - | - |
| S2670L | Class 4 | Conflicting | Likely pathogenic | fClass 5 | PS3, PM2, PP3 | - | - | - | + | - | - | + | - | - | - | - | - | - | + | - | - | - | - | - | - | - | - | - | - | - | - | - | - |
| M2676T | Class 2 | Conflicting | Uncertain significance | fClass 1 | BS3 | - | - | - | - | - | - | - | - | - | - | - | - | - | - | - | - | - | - | - | + | - | - | - | - | - | - | - | - |
| L2688P | Class 4 | Conflicting | Pathogenic | fClass 5 | PS3, PM2, PP3 | - | - | - | + | - | - | + | - | - | - | - | - | - | + | - | - | - | - | - | - | - | - | - | - | - | - | - | - |
| S2697N | Class 2 | Conflicting | Benign | fClass 1 | BS3, BS4, BP4 | - | - | - | - | - | - | - | - | - | - | - | - | - | - | - | - | - | - | - | + | + | - | - | - | + | - | - | - |
| A2717S | Class 1 | Benign | Benign | fClass 1 | BS1, BS3, BP2, BP6 | - | - | - | - | - | - | - | - | - | - | - | - | - | - | - | - | - | + | - | + | - | - | + | - | - | - | + | - |
| T2722R | Class 5 | Pathogenic | Pathogenic | fClass 5 | PS3, PM2, PP3, PP5 | - | - | - | + | - | - | + | - | - | - | - | - | - | + | - | + | - | - | - | - | - | - | - | - | - | - | - | - |
| D2723H | Class 5 | Pathogenic | Pathogenic | fClass 5 | PS3, PM2, PP1, PP3, PP5 | - | - | - | + | - | - | + | - | - | - | - | + | - | + | - | + | - | - | - | - | - | - | - | - | - | - | - | - |
| K2729N | Class 1 | Benign | Benign | fClass 2 | BS1, BS3, BS4, BP6 | - | - | - | - | - | - | - | - | - | - | - | - | - | - | - | - | - | + | - | + | + | - | - | - | - | - | + | - |
| G2748D | Class 5 | Pathogenic | Uncertain significance | fClass 5 | PS3, PP5 | - | - | - | + | - | - | - | - | - | - | - | - | - | - | - | + | - | - | - | - | - | - | - | - | - | - | - | - |
| R2784Q | Class 1 | Conflicting | Uncertain significance | fClass 5 | PS3, PP3, BS4, BP2 | - | - | - | + | - | - | - | - | - | - | - | - | - | + | - | - | - | - | - | - | + | - | + | - | - | - | - | - |
| R2842H | Class 1 | Benign | Uncertain significance | fClass 3 | BP6 | - | - | - | - | - | - | - | - | - | - | - | - | - | - | - | - | - | - | - | - | - | - | - | - | - | - | + | - |
| V2908G | Class 2 | Likely benign | Uncertain significance | fClass 4 | PP3, BS3 | - | - | - | - | - | - | - | - | - | - | - | - | - | + | - | - | - | - | - | + | - | - | - | - | - | - | - | - |
| I2944F | Class 1 | Benign | Benign | fClass 1 | BS1, BS3, BP6 | - | - | - | - | - | - | - | - | - | - | - | - | - | - | - | - | - | + | - | + | - | - | - | - | - | - | + | - |
| Y3035S | Class 2 | Conflicting | Uncertain significance | fClass 1 | - | - | - | - | - | - | - | - | - | - | - | - | - | - | - | - | - | - | - | - | - | - | - | - | - | - | - | - | - |
| R3052W | Class 5 | Pathogenic | Likely pathogenic | fClass 5 | PS3, PP1, PP3, PP5 | - | - | - | + | - | - | - | - | - | - | - | + | - | + | - | + | - | - | - | - | - | - | - | - | - | - | - | - |
| D3095E | Class 5 | Pathogenic | Likely pathogenic | fClass 5 | PS3, PP3, PP5 | - | - | - | + | - | - | - | - | - | - | - | - | - | + | - | + | - | - | - | - | - | - | - | - | - | - | - | - |
| E3096K | Class 2 | VUS | Uncertain significance | fClass 1 | - | - | - | - | - | - | - | - | - | - | - | - | - | - | - | - | - | - | - | - | - | - | - | - | - | - | - | - | - |
| Y3098H | Class 1 | Benign | Likely benign | fClass 1 | BS3, BP4, BP6 | - | - | - | - | - | - | - | - | - | - | - | - | - | - | - | - | - | - | - | + | - | - | - | - | + | - | + | - |
| N3124I | Class 5 | Pathogenic/likely pathogenic | Likely pathogenic | fClass 5 | PS3, PP1, PP3 | - | - | - | + | - | - | - | - | - | - | - | + | - | + | - | - | - | - | - | - | - | - | - | - | - | - | - | - |
| P3292L | Class 1 | Conflicting | Uncertain significance | fClass 1 | PP3 | - | - | - | - | - | - | - | - | - | - | - | - | - | + | - | - | - | - | - | - | - | - | - | - | - | - | - | - |
| Wild-type | Class 1 | Benign | Benign | fClass 1 | BA1, BS3, BP4, BP6 | - | - | - | - | - | - | - | - | - | - | - | - | - | - | - | - | + | - | - | + | - | - | - | - | + | - | + | - |
| Empty-vector | Class 5 | Pathogenic | Pathogenic | fClass 5 | PVS1, PS3, PP5 | + | - | - | + | - | - | - | - | - | - | - | - | - | - | - | + | - | - | - | - | - | - | - | - | - | - | - | - |
| K485X | Class 5 | Pathogenic | Pathogenic | fClass 5 | PVS1, PS3, PP5 | + | - | - | + | - | - | - | - | - | - | - | - | - | - | - | + | - | - | - | - | - | - | - | - | - | - | - | - |
| L997X | Class 5 | Pathogenic | Pathogenic | fClass 5 | PVS1, PS3, PP5 | + | - | - | + | - | - | - | - | - | - | - | - | - | - | - | + | - | - | - | - | - | - | - | - | - | - | - | - |
| Q1502X | Class 5 | Pathogenic | Pathogenic | fClass 5 | PVS1, PS3, PP5 | + | - | - | + | - | - | - | - | - | - | - | - | - | - | - | + | - | - | - | - | - | - | - | - | - | - | - | - |
| K1984X | Class 5 | Pathogenic | Pathogenic | fClass 5 | PVS1, PS3, PP5 | + | - | - | + | - | - | - | - | - | - | - | - | - | - | - | + | - | - | - | - | - | - | - | - | - | - | - | - |
| C2535X | Class 5 | Pathogenic | Pathogenic | fClass 4 | PVS1, PS3, PP5 | + | - | - | + | - | - | - | - | - | - | - | - | - | - | - | + | - | - | - | - | - | - | - | - | - | - | - | - |
| N3187K | Unclassified | Likely pathogenic | Uncertain significance | fClass 1 | PM2, PM6, BS3 | - | - | - | - | - | - | + | - | - | - | + | - | - | - | - | - | - | - | - | + | - | - | - | - | - | - | - | - |

| Variant | Epidemiological data: Co-occurrence likelihood | Segregation | GnomAD Exomes | HDR assay: Fold-change | HDR assay: Classification | MANO-B method: score | MANO-B method: Classification | HGVS mutation | Align-GVGD | HumanSplicingFinder: New ESS Site | ESE Site Broken | New Donor Site | Broken WT Donor Site | New Acceptor Site | Computational predictions: Mutationtaster | Polyphen2Hdiv | Provean | SIFT |
|---|---|---|---|---|---|---|---|---|---|---|---|---|---|---|---|---|---|---|
| R18H | 1.1329 | 0.0199 | 2.53E-05 | - | - | -1.38 | fClass 3 | c.53G>A | Class C0 | - | - | - | - | - | polymorphism | 0.962 | -1.17 | 0.256 |
| R18C | 0.009 | 0.0366 | 0.00E+00 | - | - | -0.23 | fClass 1 | c.52C>T | Class C15 | - | - | - | - | - | polymorphism | 1 | -1.94 | 0 |
| Y42C | 8.90E-11 | 0.00000067 | 0.001609 | - | - | -0.64 | fClass 1 | c.125A>G | Class C0 | - | - | + | - | - | disease causing | 0.09 | -1.32 | 0.113 |
| P168T | 0.08128 | 1 | 4.23E-05 | 5.59 | neutral | -0.66 | fClass 1 | c.502C>A | Class C0 | - | - | - | - | - | disease causing | 1 | -2.02 | 0 |
| S326R | 0.001 | - | 0.0007933 | - | - | -0.44 | fClass 1 | c.978C>A | Class C0 | + | - | - | - | - | polymorphism | 0.004 | -0.74 | 0.048 |
| E462G | 0.22909 | 0.01514 | 0.0002437 | - | - | -0.28 | fClass 1 | c.1385A>G | Class C0 | - | + | - | - | - | polymorphism | 0.998 | -1.01 | 0.067 |
| N588D | - | - | 2.13E-05 | - | - | -0.62 | fClass 1 | c.1762A>G | Class C0 | - | - | + | - | + | polymorphism | 0.255 | -0.56 | 0.001 |
| T598A | - | 0.000296622 | 2.25E-03 | 5.24 | neutral | -0.38 | fClass 1 | c.1792A>G | Class C0 | - | + | - | - | + | polymorphism | 0.01 | -0.12 | 0.798 |
| G602R | 1.479 | 0.00026 | 4.45E-05 | - | - | -0.19 | fClass 1 | c.1804G>A | Class C0 | - | - | - | - | + | polymorphism | 0.355 | -1.18 | 0.132 |
| L1019V | 3.46737 | 0.00066 | 0.0001144 | - | - | -0.82 | fClass 2 | c.3055C>G | Class C0 | + | + | + | - | - | polymorphism | 0.013 | -1.73 | 0.141 |
| S1172L | 0.27542 | 0.12882 | 0.001451 | 4.7 | neutral | -1.05 | fClass 2 | c.3515C>T | Class C0 | + | + | - | - | - | polymorphism | 0.97 | -4.89 | 0.035 |
| N1228D | 0.06607 | 0.00912 | 2.98E-05 | - | - | -1.14 | fClass 2 | c.3682A>G | Class C0 | + | - | - | - | - | polymorphism | 0.896 | -3.61 | 0.026 |
| D1420Y | 0.001 | - | 0.006514 | - | - | -0.75 | fClass 1 | c.4258G>T | Class C0 | + | + | - | - | - | polymorphism | 0.03 | -6.6 | 0.03 |
| F1524V | 0.06607 | 1.47911 | 2.12E-05 | - | - | -0.78 | fClass 1 | c.4570T>G | Class C45 | - | - | - | - | - | disease causing | 1 | -6.84 | 0.002 |
| K1690N | 0.29512 | 1 | 0.0001813 | - | - | 0.42 | fClass 1 | c.5070A>C | Class C0 | + | - | - | - | - | polymorphism | 0.989 | -3.66 | 0.007 |
| N1878K | 1.549157977 | 0.468988544 | 1.28E-05 | - | - | -0.11 | fClass 1 | c.5634C>G | Class C0 | - | + | - | - | + | polymorphism | 0.015 | -0.96 | 0.034 |
| H1918R | 0.043027149 | - | 2.12E-05 | - | - | -1.48 | fClass 3 | c.5753A>G | Class C0 | - | - | + | - | - | polymorphism | 0.001 | -3.25 | 0.292 |
| I1929V | 0.01175 | 0.05012 | 0.0009277 | - | - | 0.38 | fClass 1 | c.5785A>G | Class C0 | + | + | - | - | - | polymorphism | 0.004 | -0.49 | 0.307 |
| E2020K | 1.075678821 | 0.1762 | 0 | - | - | -0.82 | fClass 2 | c.6058G>A | Class C0 | - | + | - | - | - | polymorphism | 0.032 | -3.13 | 0.023 |
| H2074N | 0.48978 | 1 | 0.0007275 | - | - | -0.31 | fClass 1 | c.6220C>A | Class C0 | - | + | - | - | - | polymorphism | 0.001 | -1.8 | 0.195 |
| K2411T | 0.05754 | 1 | 6.34E-05 | - | - | -0.46 | fClass 1 | c.7232A>C | Class C0 | - | - | - | - | - | disease causing | 1 | -1.8 | 0 |
| R2418G | - | - | 0 | - | - | -1.64 | fClass 3 | c.7252A>G | Class C0 | + | - | - | - | - | polymorphism | 0.037 | -0.98 | 0.176 |
| N2436I | 0.04786 | 0.38905 | 1.27E-05 | 4.99 | neutral | -1.34 | fClass 3 | c.7307A>T | Class C0 | - | + | - | - | - | polymorphism | 0.062 | -1.95 | 0.158 |
| G2508S | 2.215 | - | 1.69E-04 | 2.559 | neutral | -2.42 | fClass 4 | c.7522G>A | Class C55 | + | - | - | - | + | disease causing | 1 | -1.8 | 0 |
| T2515I | 1.86E-03 | 0.932 | 0.0005363 | 4.15 | neutral | -0.86 | fClass 2 | c.7544C>T | Class C0 | - | + | - | - | - | polymorphism | 0.9 | -1.64 | 0.064 |
| T2607P | 1.02E+00 | - | 0 | - | - | -2.70 | fClass 5 | c.7819A>C | Class C35 | - | - | - | - | - | disease causing | 1 | -1.6 | 0 |
| G2609D | - | - | 0 | 1.35 | pathogenic | -2.81 | fClass 5 | c.7826G>A | Class C65 | + | + | - | - | - | disease causing | 1 | -1.86 | 0 |
| W2626C | 2.138 | 1 | 4.22E-06 | 1.16 | pathogenic | -2.98 | fClass 5 | c.7878G>C | Class C65 | - | + | - | - | - | disease causing | 1 | -3.47 | 0 |
| I2627F | 1.288 | 1.995 | 0 | 1.01 | pathogenic | -3.17 | fClass 5 | c.7879A>T | Class C15 | - | + | + | - | - | disease causing | 1 | -1.06 | 0.001 |
| L2647P | 1.18 | - | 0 | 0.93 | pathogenic | -2.57 | fClass 5 | c.7940T>C | Class C65 | + | + | - | - | - | disease causing | 1 | -1.86 | 0 |
| L2653P | 1.148 | 0.575 | 0 | 0.75 | pathogenic | -2.78 | fClass 5 | c.7958T>C | Class C65 | - | - | - | - | - | disease causing | 1 | -1.86 | 0 |
| R2659G | 1.102157073 | 1950 | 0 | 1.57 | pathogenic | -2.65 | fClass 5 | c.7975A>G | Class C65 | - | + | - | + | - | disease causing | 1 | -1.86 | 0 |
| S2670L | 1.225 | 2 | 0 | 0.93 | pathogenic | -3.08 | fClass 5 | c.8009C>T | Class C15 | + | - | - | - | - | disease causing | 1 | -1.78 | 0.001 |
| M2676T | 1.157084926 | - | 1.69E-05 | 5.9 | neutral | -0.49 | fClass 1 | c.8027T>C | Class C0 | - | + | - | - | - | polymorphism | 0.248 | -1.31 | 0.458 |
| L2688P | - | - | 0 | 1.02 | pathogenic | -2.92 | fClass 5 | c.8063T>C | Class C65 | + | - | - | - | - | disease causing | 1 | -2.08 | 0 |
| S2697N | 1.549157977 | 0.040326 | 3.38E-05 | 5.41 | neutral | -0.75 | fClass 1 | c.8090G>A | Class C0 | - | - | - | - | - | polymorphism | 0.001 | -0.17 | 0.435 |
| A2717S | 1.28E-03 | 0.1211 | 0.00106 | 4.08 | neutral | -0.67 | fClass 1 | c.8149G>T | Class C0 | - | + | - | - | - | polymorphism | 0.955 | -0.44 | 0.118 |
| T2722R | 1.259 | 10.715 | 0 | 1.08 | pathogenic | -2.67 | fClass 5 | c.8165C>G | Class C65 | - | + | - | - | + | disease causing | 1 | -1.78 | 0 |
| D2723H | 2 | 13731 | 0 | 1 | pathogenic | -2.52 | fClass 5 | c.8167G>C | Class C65 | - | + | - | - | - | disease causing | 1 | -2.08 | 0 |
| K2729N | 6.30957 | 0.04169 | 0.0007961 | 3.99 | neutral | -0.89 | fClass 2 | c.8187G>T | Class C0 | + | - | - | - | - | polymorphism | 0.998 | -1.31 | 0.001 |
| G2748D | 1.445 | 6.761 | 8.53E-06 | 0.68 | pathogenic | -2.57 | fClass 5 | c.8243G>A | Class C65 | - | - | - | - | - | disease causing | 1 | -2.08 | 0 |
| R2784Q | 0.048590003 | 0.000106872 | 1.27E-05 | 1.27 | pathogenic | -2.74 | fClass 5 | c.8351G>A | Class C35 | - | - | - | - | + | disease causing | 1 | -1.19 | 0 |
| R2842H | 0.05888 | 0.07586 | 6.34E-05 | 2.36 | indeterminate | -2.02 | fClass 3 | c.8525G>A | Class C25 | - | - | - | - | - | disease causing | 1 | -1.49 | 0 |
| V2908G | 0.05754 | 1 | 1.69E-05 | 2.44 | neutral | -2.30 | fClass 4 | c.8723T>G | Class C35 | + | + | + | - | - | disease causing | 1 | -1.69 | 0 |
| I2944F | - | - | 0.002928 | 4.72 | neutral | -0.68 | fClass 1 | c.8830A>T | Class C0 | + | + | - | - | - | disease causing | 0.986 | -1.12 | 0.002 |
| Y3035S | 1.666396398 | 1.070769443 | 5.11E-05 | 2.32 | indeterminate | -0.63 | fClass 1 | c.9104A>C | Class C55 | - | - | - | - | - | disease causing | 1 | -2.38 | 0 |
| R3052W | 1.48 | 3981 | 1.27E-05 | 0.86 | pathogenic | -2.91 | fClass 5 | c.9154C>T | Class C65 | + | + | - | - | - | disease causing | 1 | -2.06 | 0 |
| D3095E | 1.66 | 0.427 | 4.23E-06 | 1.07 | pathogenic | -3.00 | fClass 5 | c.9285C>A | Class C35 | - | - | - | - | + | disease causing | 1 | -1.16 | 0 |
| E3096K | - | - | 4.23E-06 | - | - | 0.00 | fClass 1 | c.9286G>A | Class C0 | - | + | - | - | - | disease causing | 1 | -1.16 | 0 |
| Y3098H | 0.195 | 5.012 | 0.0002919 | 5.64 | neutral | -0.76 | fClass 1 | c.9292T>C | Class C0 | - | - | - | - | - | polymorphism | 0 | 0.39 | 0.775 |
| N3124I | 2.074083157 | 402919.0213 | 4.22E-06 | 0.92 | pathogenic | -3.18 | fClass 5 | c.9371A>T | Class C65 | - | + | - | - | - | disease causing | 1 | -2.61 | 0 |
| P3292L | 0.059024729 | 0.704751102 | 7.18E-05 | - | - | -0.40 | fClass 1 | c.9875C>T | Class C0 | + | + | - | - | - | disease causing | 0.999 | -2.85 | 0.001 |
| Wild-type | - | - | 0.9949 | 4.46 | neutral | 0.00 | fClass 1 | - | Class C0 | - | - | - | - | - | polymorphism | - | 0.62 | 1 |
| Empty-vector | - | - | - | - | - | -2.54 | fClass 5 | - | Non-sense | - | - | - | - | - | - | - | - | - |
| K485X | - | - | - | - | - | -3.03 | fClass 5 | c.1453A>T | Non-sense | + | + | - | - | - | - | - | - | - |
| L997X | - | - | - | - | - | -3.50 | fClass 5 | c.2990T>A | Non-sense | - | - | - | - | - | - | - | - | - |
| Q1502X | - | - | - | - | - | -2.99 | fClass 5 | c.4504C>T | Non-sense | - | + | - | - | - | - | - | - | - |
| K1984X | - | - | - | - | - | -2.96 | fClass 5 | c.5950A>T | Non-sense | - | + | - | - | - | - | - | - | - |
| C2535X | - | - | - | - | - | -2.18 | fClass 4 | c.7604_7605delinsAG | Non-sense | - | - | - | - | - | - | - | - | - |
| N3187K | - | - | 0 | 5.22 | neutral | -0.07 | fClass 1 | c.9561T>A | Class C0 | + | - | - | - | - | polymorphism | 0.111 | -1.26 | 0.004 |

PP3, BP4, deleterious threshold: PolyPhen-2 score > 0.15, Align-GVGD class > C15, SIFT score < 0.05, PROVEAN score < -2.5

BP2, neutral likelihood threshold:Co-occurrence < 0.053

BS4, neutral likelihood threshold: Segregation < 0.053

PP1, deleteriousl likelihood threshold: Segregation > 18.7

BS1, Maximal expected allele frequency for a non-common pathogenic BRCA2 variant: 0.075%

PS3, BS3: based on HDR assays
