## Supplementary_Table_2 for "High-Throughput Functional Evaluation of *BRCA2* Variants of Unknown Significance"

Supplementary Table 2: Primers used in digital droplet PCR (ddPCR) and real-time qRT-PCR.

| Primer | Sequense |
|---|---|
| qRT-PCR, BRCA2 exon 11, forward | 5′-AGCCAGTGGTAAAATCGTTTGTG-3′ |
| qRT-PCR, BRCA2 exon 11, reverse | 5′-CCTCTGAATCATCCAATGCCTCGTA-3′ |
| qRT-PCR, ACTB, forward | 5′-AGAGCTACGAGCTGCCTGAC-3′ |
| qRT-PCR, ACTB, reverse | 5′-AGCACTGTGTTGGCGTACAG-3′ |
| ddPCR, BRCA2 cDNA, forward | 5′-CCACCCACCCTTAG-3′ |
| ddPCR, BRCA2 cDNA, reverse | 5′-CATGAGGAAATACAGTTTCA-3′ |
| ddPCR, BRCA2 cDNA, probe | 5′-FAM-ACTGTGCTCATAGTCAGAAATG-BHQ1-3′ |
| ddPCR, BRCA2 intron forward | 5′-GTGTGATACATGCTTTGAA-3′ |
| ddPCR, BRCA2 intron reverse | 5′-TGCCCATTACCTATATTCC-3′ |
| ddPCR, BRCA2 intron, probe | 5′-HEX-TTCATGGCTATAGGGAGTGCA-3′ |

Supplementary Table 2: Raw data of Supplementary Figure S1A.

| Variant | Copy number | 95% CI | |
|---|---|---|---|
| | | Lower limit | Upper limit |
| Wild-type | 10.6 | 10.04 | 11.14 |
| P168T | 10.3 | 9.82 | 10.72 |
| T598A | 10.3 | 9.94 | 10.7 |
| S1172L | 8.9 | 8.52 | 9.3 |
| K1530N | 10.3 | 9.92 | 10.72 |
| E1695V | 8.7 | 8.38 | 9.1 |
| G1696V | 8.6 | 8.26 | 9 |
| R2336G | 12.6 | 11.2 | 14 |
| N2436I | 9 | 8.62 | 9.4 |
| L2604P | 10.7 | 10.06 | 11.28 |
| P2639L | 9.4 | 8.92 | 9.84 |
| D2723H | 8 | 7.54 | 8.38 |
| R2842H | 10 | 9.46 | 10.56 |
| R2896C | 12 | 11.46 | 12.5 |
| A2911E | 13.3 | 12.78 | 13.86 |
| D2913H | 9.2 | 8.78 | 9.68 |
| R3052W | 11.6 | 10 | 13.2 |
| D3095E | 9.8 | 9.3 | 10.22 |
| N3124I | 12.9 | 12.12 | 13.7 |
| S3291C | 11.9 | 11.28 | 12.52 |
| Empty-vector | 0 | 0 | 0 |

CI: confidence interval

Supplementary Table 2: Raw data of Supplementary Figure S1B.

| Variant | BRCA2 mRNA expression | |
|---|---|---|
| | Mean | SD |
| Wild-type | 0.577 | 0.082 |
| P168T | 0.853 | 0.321 |
| T598A | 0.844 | 0.204 |
| S1172L | 0.627 | 0.191 |
| K1530N | 0.884 | 0.461 |
| E1695V | 0.690 | 0.228 |
| G1696V | 0.601 | 0.248 |
| R2336G | 0.758 | 0.146 |
| N2436I | 1.017 | 0.083 |
| L2604P | 0.769 | 0.184 |
| P2639L | 0.860 | 0.322 |
| D2723H | 0.954 | 0.636 |
| R2842H | 0.881 | 0.387 |
| R2896C | 1.522 | 0.990 |
| A2911E | 1.746 | 0.725 |
| D2913H | 0.919 | 0.467 |
| R3052W | 0.955 | 0.213 |
| D3095E | 0.614 | 0.177 |
| N3124I | 1.046 | 0.275 |
| S3291C | 0.959 | 0.218 |
| Empty-vector | 0.000 | 0.000 |

SD: standard deviation

Supplementary Table 2: Raw data of Supplementary Figure S2A.

| Drug | Concentration (nmol/L) | Wild-type | | P168T | | T598A | | S1172L | | K1530N | | E1695V | | G1696V | | R2336G | |
|---|---|---|---|---|---|---|---|---|---|---|---|---|---|---|---|---|---|
| | | Cell viability (%) | SEM | Cell viability (%) | SEM | Cell viability (%) | SEM | Cell viability (%) | SEM | Cell viability (%) | SEM | Cell viability (%) | SEM | Cell viability (%) | SEM | Cell viability (%) | SEM |
| Olaparib | 0 | 100.00 | 1.67 | 100.00 | 2.20 | 100.00 | 1.18 | 100.00 | 2.89 | 100.00 | 1.24 | 100.00 | 2.02 | 100.00 | 1.62 | 100.00 | 0.54 |
| Olaparib | 10 | 96.71 | 0.78 | 98.90 | 2.84 | 104.33 | 1.56 | 99.37 | 3.29 | 102.95 | 2.18 | 104.18 | 2.49 | 100.61 | 2.45 | 105.23 | 2.09 |
| Olaparib | 50 | 88.52 | 1.53 | 88.87 | 2.08 | 98.21 | 0.57 | 100.42 | 2.01 | 98.03 | 1.33 | 96.49 | 1.00 | 94.47 | 0.77 | 107.23 | 1.78 |
| Olaparib | 100 | 85.93 | 0.12 | 88.06 | 0.86 | 97.13 | 1.69 | 100.66 | 1.55 | 97.09 | 1.68 | 96.48 | 1.16 | 89.63 | 1.40 | 104.70 | 2.40 |
| Olaparib | 500 | 80.32 | 2.11 | 80.97 | 0.98 | 86.98 | 1.68 | 84.84 | 2.41 | 88.11 | 2.29 | 86.55 | 0.64 | 81.79 | 0.58 | 95.40 | 1.55 |
| Olaparib | 1000 | 77.65 | 1.32 | 75.23 | 0.97 | 82.05 | 0.76 | 75.84 | 1.00 | 82.48 | 1.73 | 77.65 | 2.02 | 68.56 | 2.09 | 84.65 | 0.79 |
| Olaparib | 5000 | 36.40 | 0.89 | 25.54 | 0.83 | 34.79 | 0.78 | 15.96 | 0.91 | 23.56 | 0.39 | 22.00 | 1.21 | 18.29 | 1.14 | 48.98 | 1.75 |
| Niraparib | 0 | 100.00 | 1.11 | 100.00 | 1.44 | 100.00 | 2.41 | 100.00 | 1.66 | 100.00 | 2.95 | 100.00 | 2.62 | 100.00 | 0.95 | 100.00 | 3.78 |
| Niraparib | 5 | 94.71 | 1.86 | 99.33 | 2.49 | 100.37 | 0.96 | 102.03 | 2.53 | 102.58 | 5.09 | 97.90 | 1.37 | 97.32 | 1.37 | 103.15 | 3.85 |
| Niraparib | 10 | 97.88 | 1.50 | 100.60 | 2.31 | 105.84 | 1.22 | 104.89 | 3.26 | 103.20 | 3.74 | 95.73 | 2.53 | 95.95 | 1.54 | 105.42 | 2.03 |
| Niraparib | 50 | 98.42 | 1.35 | 96.90 | 1.45 | 103.13 | 2.24 | 100.15 | 0.89 | 108.05 | 2.05 | 97.52 | 0.44 | 94.96 | 1.51 | 103.91 | 2.37 |
| Niraparib | 100 | 91.08 | 1.45 | 95.83 | 0.51 | 100.63 | 2.16 | 103.55 | 2.02 | 105.56 | 3.30 | 97.69 | 1.96 | 95.55 | 1.67 | 105.50 | 2.91 |
| Niraparib | 500 | 75.10 | 1.27 | 78.05 | 1.08 | 84.63 | 1.44 | 84.42 | 0.80 | 80.59 | 4.90 | 78.09 | 0.79 | 62.70 | 2.22 | 84.72 | 2.37 |
| Niraparib | 1000 | 56.84 | 2.67 | 47.42 | 2.50 | 55.78 | 2.22 | 46.58 | 3.04 | 51.07 | 0.90 | 35.24 | 1.96 | 28.27 | 0.80 | 69.73 | 1.12 |
| Rucaparib | 0 | 100.00 | 2.00 | 100.00 | 1.87 | 100.00 | 1.07 | 100.00 | 1.77 | 100.00 | 1.27 | 100.00 | 1.02 | 100.00 | 2.00 | 100.00 | 0.54 |
| Rucaparib | 10 | 87.59 | 1.36 | 99.45 | 1.45 | 104.97 | 1.47 | 106.25 | 3.11 | 110.62 | 1.78 | 99.68 | 1.74 | 100.75 | 1.31 | 99.43 | 2.39 |
| Rucaparib | 50 | 83.23 | 0.78 | 94.94 | 0.69 | 102.24 | 1.11 | 106.11 | 2.50 | 103.38 | 4.64 | 98.86 | 2.41 | 93.28 | 2.37 | 95.09 | 1.16 |
| Rucaparib | 100 | 81.32 | 0.73 | 96.05 | 3.93 | 99.44 | 2.24 | 107.95 | 2.87 | 103.51 | 4.12 | 97.98 | 3.04 | 88.45 | 2.31 | 91.78 | 0.96 |
| Rucaparib | 500 | 77.04 | 1.28 | 88.07 | 1.27 | 90.68 | 2.00 | 95.01 | 2.18 | 91.29 | 2.35 | 88.06 | 1.74 | 77.11 | 3.74 | 86.83 | 0.40 |
| Rucaparib | 1000 | 69.38 | 0.81 | 80.60 | 0.97 | 83.79 | 1.65 | 87.28 | 1.66 | 85.06 | 1.16 | 80.83 | 2.19 | 69.95 | 1.75 | 80.03 | 1.37 |
| Rucaparib | 5000 | 34.94 | 1.48 | 28.26 | 1.40 | 40.64 | 1.46 | 23.82 | 1.87 | 28.39 | 1.12 | 19.82 | 0.48 | 16.45 | 0.37 | 41.54 | 2.21 |
| CBDCA | 0 | 100.00 | 1.30 | 100.00 | 2.33 | 100.00 | 0.94 | 100.00 | 2.42 | 100.00 | 0.59 | 100.00 | 2.11 | 100.00 | 1.30 | 100.00 | 2.12 |
| CBDCA | 100 | 98.63 | 0.81 | 101.70 | 2.37 | 102.80 | 0.57 | 102.51 | 1.95 | 102.65 | 2.54 | 100.60 | 1.83 | 100.50 | 3.19 | 102.10 | 1.90 |
| CBDCA | 500 | 88.27 | 1.84 | 96.54 | 1.07 | 95.72 | 1.10 | 99.28 | 1.49 | 95.50 | 2.55 | 93.31 | 1.16 | 88.52 | 3.72 | 92.34 | 1.88 |
| CBDCA | 1000 | 79.01 | 0.48 | 80.91 | 0.60 | 84.36 | 1.11 | 83.54 | 2.23 | 84.95 | 1.01 | 83.18 | 2.11 | 75.91 | 1.94 | 81.13 | 2.06 |
| CBDCA | 5000 | 18.23 | 0.62 | 15.31 | 0.50 | 13.11 | 0.47 | 11.01 | 0.42 | 14.75 | 0.67 | 10.26 | 0.33 | 8.75 | 0.33 | 20.37 | 0.68 |
| CBDCA | 10000 | 5.21 | 0.29 | 4.29 | 0.24 | 4.27 | 0.11 | 4.50 | 0.21 | 5.24 | 0.22 | 4.47 | 0.16 | 3.77 | 0.17 | 6.21 | 0.31 |

| Drug | Concentration (nmol/L) | N2436I | | L2604P | | P2639L | | D2723H | | R2842H | | R2896C | | A2911E | | D2913H | |
|---|---|---|---|---|---|---|---|---|---|---|---|---|---|---|---|---|---|
| | | Cell viability (%) | SEM | Cell viability (%) | SEM | Cell viability (%) | SEM | Cell viability (%) | SEM | Cell viability (%) | SEM | Cell viability (%) | SEM | Cell viability (%) | SEM | Cell viability (%) | SEM |
| Olaparib | 0 | 100.00 | 0.41 | 100.00 | 0.74 | 100.00 | 0.48 | 100.00 | 1.54 | 100.00 | 2.04 | 100.00 | 0.91 | 100.00 | 0.66 | 100.00 | 3.60 |
| Olaparib | 10 | 99.88 | 1.39 | 94.60 | 1.96 | 100.20 | 0.46 | 90.10 | 1.33 | 98.19 | 1.91 | 100.63 | 2.25 | 100.81 | 0.76 | 108.10 | 2.16 |
| Olaparib | 50 | 103.69 | 2.12 | 85.12 | 1.03 | 97.82 | 0.96 | 73.86 | 0.56 | 88.06 | 1.28 | 100.08 | 1.55 | 97.56 | 1.35 | 99.83 | 3.76 |
| Olaparib | 100 | 109.70 | 0.80 | 80.49 | 0.76 | 94.39 | 1.28 | 63.86 | 1.70 | 77.73 | 0.44 | 97.12 | 2.78 | 101.18 | 1.81 | 100.50 | 1.99 |
| Olaparib | 500 | 102.43 | 1.76 | 46.89 | 1.78 | 71.53 | 1.24 | 26.40 | 0.94 | 41.81 | 1.94 | 92.26 | 2.56 | 93.36 | 1.64 | 79.64 | 2.51 |
| Olaparib | 1000 | 91.96 | 1.36 | 26.06 | 0.58 | 44.50 | 1.64 | 13.29 | 0.77 | 23.49 | 1.20 | 86.37 | 1.32 | 82.88 | 2.63 | 51.56 | 1.37 |
| Olaparib | 5000 | 33.79 | 0.77 | 5.95 | 0.22 | 8.70 | 0.29 | 3.73 | 0.08 | 5.18 | 0.25 | 59.54 | 2.33 | 36.43 | 0.99 | 9.71 | 0.47 |
| Niraparib | 0 | 100.00 | 1.05 | 100.00 | 1.67 | 100.00 | 2.66 | 100.00 | 2.39 | 100.00 | 1.12 | 100.00 | 2.79 | 100.00 | 1.69 | 100.00 | 3.46 |
| Niraparib | 5 | 99.17 | 1.48 | 94.88 | 1.86 | 95.28 | 1.56 | 100.45 | 1.77 | 96.66 | 0.83 | 101.12 | 3.36 | 102.93 | 1.87 | 104.87 | 4.36 |
| Niraparib | 10 | 104.84 | 0.77 | 94.91 | 1.52 | 94.55 | 2.02 | 98.50 | 1.24 | 97.43 | 1.43 | 99.64 | 2.74 | 100.78 | 1.65 | 108.01 | 2.87 |
| Niraparib | 50 | 104.26 | 1.51 | 88.27 | 1.08 | 99.63 | 1.41 | 90.93 | 1.52 | 91.86 | 1.37 | 100.84 | 1.06 | 98.64 | 1.73 | 107.52 | 3.03 |
| Niraparib | 100 | 104.18 | 2.01 | 83.52 | 1.82 | 95.63 | 2.45 | 77.53 | 1.15 | 86.57 | 1.85 | 95.99 | 2.40 | 97.14 | 2.28 | 103.69 | 1.15 |
| Niraparib | 500 | 93.30 | 0.61 | 20.67 | 1.35 | 51.70 | 1.96 | 17.17 | 0.80 | 23.84 | 1.17 | 87.67 | 1.60 | 81.07 | 1.07 | 61.56 | 1.91 |
| Niraparib | 1000 | 61.87 | 1.62 | 7.55 | 0.36 | 15.78 | 0.40 | 6.92 | 0.20 | 9.53 | 0.12 | 81.05 | 2.48 | 55.59 | 1.47 | 21.25 | 0.78 |
| Rucaparib | 0 | 100.00 | 0.62 | 100.00 | 1.93 | 100.00 | 0.42 | 100.00 | 3.03 | 100.00 | 0.88 | 100.00 | 0.80 | 100.00 | 1.14 | 100.00 | 1.07 |
| Rucaparib | 10 | 94.16 | 1.74 | 97.35 | 1.07 | 96.23 | 1.41 | 90.30 | 3.30 | 91.86 | 1.39 | 99.04 | 2.60 | 100.54 | 2.20 | 105.58 | 2.30 |
| Rucaparib | 50 | 96.72 | 0.84 | 82.39 | 1.03 | 95.18 | 1.22 | 71.50 | 1.77 | 77.31 | 1.45 | 100.47 | 1.97 | 96.95 | 2.06 | 99.73 | 1.33 |
| Rucaparib | 100 | 101.29 | 2.01 | 72.88 | 1.84 | 90.45 | 1.46 | 59.64 | 1.90 | 70.70 | 1.70 | 95.07 | 3.06 | 97.46 | 0.84 | 97.64 | 1.46 |
| Rucaparib | 500 | 94.01 | 0.80 | 41.17 | 1.12 | 70.38 | 2.03 | 27.19 | 0.51 | 42.25 | 1.75 | 92.33 | 1.22 | 85.35 | 2.55 | 77.17 | 2.12 |
| Rucaparib | 1000 | 91.62 | 1.07 | 20.34 | 0.77 | 51.14 | 1.20 | 14.82 | 0.63 | 23.96 | 1.02 | 84.29 | 2.64 | 78.40 | 2.08 | 56.84 | 1.79 |
| Rucaparib | 5000 | 34.70 | 1.50 | 5.87 | 0.18 | 7.19 | 0.22 | 5.22 | 0.25 | 5.98 | 0.18 | 56.71 | 1.85 | 34.40 | 1.18 | 11.89 | 0.55 |
| CBDCA | 0 | 100.00 | 1.72 | 100.00 | 1.33 | 100.00 | 1.47 | 100.00 | 3.38 | 100.00 | 0.77 | 100.00 | 1.54 | 100.00 | 0.77 | 100.00 | 5.57 |
| CBDCA | 100 | 98.13 | 2.33 | 93.57 | 2.01 | 95.12 | 1.87 | 106.11 | 2.28 | 101.57 | 1.88 | 91.79 | 1.98 | 89.73 | 2.47 | 91.83 | 9.31 |
| CBDCA | 500 | 101.22 | 2.07 | 73.26 | 1.98 | 85.12 | 0.26 | 68.80 | 1.73 | 81.47 | 0.82 | 85.52 | 3.14 | 88.53 | 1.43 | 89.76 | 6.75 |
| CBDCA | 1000 | 87.69 | 2.55 | 30.89 | 1.00 | 55.85 | 2.68 | 26.40 | 0.63 | 43.21 | 1.56 | 78.22 | 2.60 | 80.62 | 2.55 | 53.16 | 3.17 |
| CBDCA | 5000 | 12.37 | 0.55 | 4.12 | 0.11 | 4.87 | 0.15 | 3.78 | 0.12 | 4.84 | 0.07 | 39.05 | 1.72 | 17.64 | 0.21 | 16.19 | 0.83 |
| CBDCA | 10000 | 4.48 | 0.18 | 2.98 | 0.09 | 2.80 | 0.12 | 2.55 | 0.04 | 3.33 | 0.10 | 8.02 | 0.28 | 4.69 | 0.10 | 9.65 | 0.44 |

| Drug | Concentration (nmol/L) | R3052W | | D3095E | | N3124I | | S3291C | | Empty vector | | Parental | | Trunsduction (-) | |
|---|---|---|---|---|---|---|---|---|---|---|---|---|---|---|---|
| | | Cell viability (%) | SEM | Cell viability (%) | SEM | Cell viability (%) | SEM | Cell viability (%) | SEM | Cell viability (%) | SEM | Cell viability (%) | SEM | Cell viability (%) | SEM |
| Olaparib | 0 | 100.00 | 2.09 | 100.00 | 0.34 | 100.00 | 0.55 | 100.00 | 1.78 | 100.00 | 2.67 | 98.44 | 2.02 | 100.00 | 2.47 |
| Olaparib | 10 | 97.06 | 1.74 | 96.34 | 2.17 | 97.82 | 1.37 | 94.92 | 0.84 | 96.83 | 0.60 | 98.26 | 1.01 | 89.77 | 3.61 |
| Olaparib | 50 | 78.86 | 2.56 | 87.19 | 0.70 | 88.85 | 1.63 | 88.06 | 0.62 | 85.08 | 1.75 | 98.37 | 0.81 | 81.03 | 2.07 |
| Olaparib | 100 | 68.91 | 3.69 | 82.01 | 1.74 | 78.57 | 1.09 | 83.29 | 1.40 | 77.31 | 1.03 | 99.60 | 0.25 | 71.23 | 0.92 |
| Olaparib | 500 | 26.96 | 1.09 | 48.34 | 0.65 | 46.17 | 1.48 | 68.94 | 1.31 | 41.49 | 0.89 | 98.74 | 1.34 | 30.24 | 0.68 |
| Olaparib | 1000 | 14.06 | 0.82 | 25.70 | 1.05 | 23.12 | 0.42 | 46.55 | 0.65 | 18.68 | 0.74 | 97.66 | 0.79 | 15.71 | 0.62 |
| Olaparib | 5000 | 3.66 | 0.10 | 4.67 | 0.14 | 5.20 | 0.14 | 9.43 | 0.25 | 3.97 | 0.16 | 100.00 | 1.40 | 4.62 | 0.11 |
| Niraparib | 0 | 100.00 | 1.46 | 100.00 | 2.39 | 100.00 | 0.65 | 100.00 | 1.12 | 100.00 | 1.77 | 100.00 | 0.88 | 100.00 | 0.75 |
| Niraparib | 5 | 96.34 | 2.11 | 101.17 | 1.82 | 99.34 | 1.21 | 95.38 | 1.01 | 94.46 | 2.29 | 95.27 | 0.86 | 98.13 | 3.57 |
| Niraparib | 10 | 95.39 | 2.97 | 98.74 | 2.30 | 98.56 | 1.03 | 94.06 | 1.52 | 95.68 | 1.22 | 95.58 | 1.01 | 99.70 | 2.28 |
| Niraparib | 50 | 85.31 | 3.25 | 96.60 | 1.59 | 93.39 | 0.90 | 92.71 | 1.87 | 92.37 | 1.67 | 95.73 | 0.81 | 92.33 | 2.92 |
| Niraparib | 100 | 69.68 | 3.92 | 92.69 | 1.82 | 86.69 | 1.27 | 89.51 | 0.94 | 83.89 | 1.19 | 92.10 | 0.56 | 80.37 | 2.13 |
| Niraparib | 500 | 12.13 | 0.46 | 28.79 | 0.69 | 30.89 | 1.43 | 47.07 | 2.00 | 22.08 | 1.09 | 94.58 | 0.50 | 19.30 | 0.62 |
| Niraparib | 1000 | 4.81 | 0.25 | 9.53 | 0.14 | 10.35 | 0.32 | 16.62 | 1.13 | 6.89 | 0.35 | 95.23 | 0.61 | 7.27 | 0.24 |
| Rucaparib | 0 | 100.00 | 1.98 | 100.00 | 1.40 | 100.00 | 0.83 | 100.00 | 1.14 | 100.00 | 2.08 | 100.00 | 0.53 | 100.00 | 1.34 |
| Rucaparib | 10 | 93.78 | 1.60 | 99.89 | 1.34 | 94.64 | 0.26 | 97.16 | 0.99 | 95.20 | 1.51 | 95.30 | 0.85 | 90.68 | 0.96 |
| Rucaparib | 50 | 69.86 | 2.31 | 86.85 | 1.04 | 83.67 | 1.09 | 89.57 | 1.55 | 79.00 | 2.02 | 94.23 | 0.53 | 74.77 | 1.45 |
| Rucaparib | 100 | 60.36 | 1.74 | 80.24 | 1.31 | 73.72 | 1.08 | 83.72 | 1.23 | 73.27 | 1.23 | 95.32 | 0.62 | 63.61 | 1.94 |
| Rucaparib | 500 | 22.59 | 0.42 | 47.80 | 1.03 | 42.71 | 1.33 | 66.66 | 1.30 | 43.73 | 1.41 | 95.84 | 0.85 | 29.38 | 1.79 |
| Rucaparib | 1000 | 13.33 | 0.36 | 29.00 | 1.84 | 26.92 | 0.50 | 50.78 | 1.10 | 22.89 | 0.66 | 95.25 | 0.88 | 16.75 | 0.71 |
| Rucaparib | 5000 | 4.36 | 0.11 | 6.53 | 0.14 | 6.33 | 0.24 | 10.38 | 0.41 | 4.54 | 0.12 | 97.31 | 0.95 | 5.68 | 0.29 |
| CBDCA | 0 | 100.00 | 5.04 | 100.00 | 0.83 | 100.00 | 1.37 | 100.00 | 1.97 | 100.00 | 1.35 | 100.00 | 0.58 | 100.00 | 1.48 |
| CBDCA | 100 | 113.37 | 6.95 | 100.51 | 0.66 | 98.61 | 1.05 | 90.89 | 1.28 | 93.03 | 1.31 | 94.69 | 1.18 | 92.23 | 1.28 |
| CBDCA | 500 | 88.77 | 11.56 | 86.47 | 2.26 | 80.66 | 0.86 | 85.11 | 1.75 | 73.65 | 1.20 | 95.36 | 0.97 | 66.23 | 2.20 |
| CBDCA | 1000 | 52.96 | 5.66 | 51.55 | 0.51 | 40.80 | 1.41 | 67.19 | 1.77 | 32.56 | 1.05 | 94.43 | 0.99 | 21.78 | 0.89 |
| CBDCA | 5000 | 24.50 | 4.00 | 4.78 | 0.11 | 4.61 | 0.08 | 7.21 | 0.10 | 3.51 | 0.15 | 91.89 | 1.43 | 3.98 | 0.12 |
| CBDCA | 10000 | 11.73 | 0.45 | 3.05 | 0.07 | 3.05 | 0.12 | 3.59 | 0.08 | 2.80 | 0.11 | 89.05 | 0.83 | 2.53 | 0.11 |

Supplementary Table 2: Raw data of Supplementary Figure 2B.

| Drug | | Wild-type | | P168T | | T598A | | S1172L | | K1530N | | E1695V | | G1696V | | R2336G | |
|---|---|---|---|---|---|---|---|---|---|---|---|---|---|---|---|---|---|
| | Concentration (nmol/L) | Cell viability (%) | SEM | Cell viability (%) | SEM | Cell viability (%) | SEM | Cell viability (%) | SEM | Cell viability (%) | SEM | Cell viability (%) | SEM | Cell viability (%) | SEM | Cell viability (%) | SEM |
| Olaparib | 1000 | 100.00 | 1.70 | 96.89 | 1.24 | 105.66 | 0.98 | 97.67 | 1.29 | 106.21 | 2.22 | 99.99 | 2.60 | 88.28 | 2.70 | 109.01 | 1.01 |
| Niraparib | 500 | 100.00 | 1.70 | 103.92 | 1.44 | 112.69 | 1.91 | 112.41 | 1.06 | 107.30 | 6.53 | 103.98 | 1.05 | 83.48 | 2.96 | 112.81 | 3.16 |
| Rucaparib | 1000 | 100.00 | 1.16 | 116.17 | 1.39 | 120.78 | 2.38 | 125.81 | 2.39 | 122.60 | 1.67 | 116.51 | 3.15 | 100.83 | 2.53 | 115.35 | 1.98 |
| CBDCA | 1000 | 100.00 | 0.61 | 102.40 | 0.75 | 106.77 | 1.40 | 105.73 | 2.83 | 107.52 | 1.27 | 105.28 | 2.67 | 96.08 | 2.45 | 102.68 | 2.61 |

| Drug | N2436I | | L2604P | | P2639L | | D2723H | | R2842H | | R2896C | | A2911E | | D2913H | |
|---|---|---|---|---|---|---|---|---|---|---|---|---|---|---|---|---|
| | Cell viability (%) | SEM | Cell viability (%) | SEM | Cell viability (%) | SEM | Cell viability (%) | SEM | Cell viability (%) | SEM | Cell viability (%) | SEM | Cell viability (%) | SEM | Cell viability (%) | SEM |
| Olaparib | 118.42 | 1.75 | 33.55 | 0.75 | 57.30 | 2.11 | 17.12 | 1.00 | 30.25 | 1.55 | 111.22 | 1.70 | 106.73 | 3.39 | 66.40 | 1.76 |
| Niraparib | 124.23 | 0.81 | 27.52 | 1.79 | 68.84 | 2.61 | 22.86 | 1.07 | 31.75 | 1.55 | 116.73 | 2.14 | 107.94 | 1.43 | 81.97 | 2.55 |
| Rucaparib | 132.06 | 1.54 | 29.31 | 1.12 | 73.71 | 1.73 | 21.36 | 0.90 | 34.54 | 1.47 | 121.49 | 3.80 | 113.00 | 2.99 | 81.92 | 2.58 |
| CBDCA | 110.98 | 3.23 | 39.09 | 1.26 | 70.68 | 3.39 | 33.41 | 0.80 | 54.68 | 1.97 | 99.00 | 3.29 | 102.03 | 3.22 | 67.28 | 4.02 |

| Drug | R3052W | | D3095E | | N3124I | | S3291C | | Empty vector | | Parental | | W | |
|---|---|---|---|---|---|---|---|---|---|---|---|---|---|---|
| | Cell viability (%) | SEM | Cell viability (%) | SEM | Cell viability (%) | SEM | Cell viability (%) | SEM | Cell viability (%) | SEM | Cell viability (%) | SEM | Cell viability (%) | SEM |
| Olaparib | 18.11 | 1.05 | 33.10 | 1.35 | 29.77 | 0.54 | 59.95 | 0.84 | 24.06 | 0.96 | 125.76 | 1.02 | 20.23 | 0.80 |
| Niraparib | 16.15 | 0.61 | 38.33 | 0.91 | 41.13 | 1.91 | 62.68 | 2.66 | 29.40 | 1.46 | 125.93 | 0.67 | 25.69 | 0.82 |
| Rucaparib | 19.22 | 0.52 | 41.79 | 2.65 | 38.80 | 0.73 | 73.19 | 1.58 | 32.99 | 0.96 | 137.29 | 1.26 | 24.15 | 1.03 |
| CBDCA | 67.03 | 7.16 | 65.25 | 0.64 | 51.63 | 1.79 | 85.04 | 2.24 | 41.21 | 1.33 | 119.50 | 1.25 | 27.57 | 1.13 |

Supplementary Table 2: Raw data of Supplementary Figure 11

| Variant | GFP positive fraction | |
|---|---|---|
| | Mean | SD |
| Wild-type | 5 | - |
| P168T | 5.59 | 0.12 |
| T598A | 5.24 | 0.38 |
| S1172L | 4.70 | 0.14 |
| M1168I | 4.75 | 0.15 |
| K1530N | 4.33 | 0.05 |
| E1695V | 4.44 | 0.06 |
| G1696V | 4.70 | 0.06 |
| R2336G | 5.51 | 0.14 |
| T2412I | 4.59 | 0.42 |
| N2436I | 4.99 | 0.27 |
| R2896C | 5.08 | 0.23 |
| A2911E | 4.25 | 0.51 |
| P2639L | 3.15 | 0.05 |
| N3187K | 5.22 | 0.82 |
| D2913H | 2.19 | 0.34 |
| S3291C | 3.34 | 0.25 |
| L2604P | 0.91 | 0.07 |
| D2723H | 1 | - |
| R2842H | 2.21 | 0.31 |
| V2908G | 1.71 | 0.06 |
| R3052W | 0.92 | 0.01 |
| D3095E | 0.86 | 0.09 |
| N3124I | 0.81 | 0.02 |
| Empty-vector | 1.13 | 0.19 |

Values are normalized to the standard values of 1 for D2723H and 5 for wild-type.
