## Supplementary_Table_3 for "High-Throughput Functional Evaluation of *BRCA2* Variants of Unknown Significance"

Supplementary Table 3: Primers and the amplified barcode sequence used for deep sequencing in the MANO-B method.

| Primer | Sequense |
|---|---|
| MANO-B_501 | AATGATACGGCGACCACCGAGATCTACACTATAGCCTTCGCAGTGCATGTGCCTGAGTAAG |
| MANO-B_502 | AATGATACGGCGACCACCGAGATCTACACATAGAGGCTCGCAGTGCATGTGCCTGAGTAAG |
| MANO-B_503 | AATGATACGGCGACCACCGAGATCTACACCCTATCCTTCGCAGTGCATGTGCCTGAGTAAG |
| MANO-B_504 | AATGATACGGCGACCACCGAGATCTACACGGCTCTGATCGCAGTGCATGTGCCTGAGTAAG |
| MANO-B_505 | AATGATACGGCGACCACCGAGATCTACACAGGCGAAGTCGCAGTGCATGTGCCTGAGTAAG |
| MANO-B_506 | AATGATACGGCGACCACCGAGATCTACACTAATCTTATCGCAGTGCATGTGCCTGAGTAAG |
| MANO-B_507 | AATGATACGGCGACCACCGAGATCTACACCAGGACGTTCGCAGTGCATGTGCCTGAGTAAG |
| MANO-B_508 | AATGATACGGCGACCACCGAGATCTACACGTACTGACTCGCAGTGCATGTGCCTGAGTAAG |
| MANO-B_701 | CAAGCAGAAGACGGCATACGAGATCGAGTAATCTCGTTTAGTGAACCGTCAGATCG |
| MANO-B_702 | CAAGCAGAAGACGGCATACGAGATTCTCCGGACTCGTTTAGTGAACCGTCAGATCG |
| MANO-B_703 | CAAGCAGAAGACGGCATACGAGATAATGAGCGCTCGTTTAGTGAACCGTCAGATCG |
| MANO-B_704 | CAAGCAGAAGACGGCATACGAGATGGAATCTCCTCGTTTAGTGAACCGTCAGATCG |
| MANO-B_705 | CAAGCAGAAGACGGCATACGAGATTTCTGAATCTCGTTTAGTGAACCGTCAGATCG |
| MANO-B_706 | CAAGCAGAAGACGGCATACGAGATACGAATTCCTCGTTTAGTGAACCGTCAGATCG |
| MANO-B_707 | CAAGCAGAAGACGGCATACGAGATAGCTTCAGCTCGTTTAGTGAACCGTCAGATCG |
| MANO-B_708 | CAAGCAGAAGACGGCATACGAGATGCGCATTACTCGTTTAGTGAACCGTCAGATCG |
| MANO-B_709 | CAAGCAGAAGACGGCATACGAGATCATAGCCGCTCGTTTAGTGAACCGTCAGATCG |
| MANO-B_710 | CAAGCAGAAGACGGCATACGAGATTTCGCGGACTCGTTTAGTGAACCGTCAGATCG |
| MANO-B_711 | CAAGCAGAAGACGGCATACGAGATGCGCGAGACTCGTTTAGTGAACCGTCAGATCG |
| MANO-B_712 | CAAGCAGAAGACGGCATACGAGATCTATCGCTCTCGTTTAGTGAACCGTCAGATCG |
| MANO-B_index-1 | CAGGCGATCTGACGGTTCACTAAACGAG |
| MANO-B_index-2 | AATGATACGGCGACCACCGAGATCTACAC |
| MANO-B_read-1 | CGATTCGCTTGCAGTGTTACACCTGAGTC |
| MANO-B_read-2 | CATCCACGCTGTTTTGACCTCCATAGAAG |
| Amplified sequence | CTCGTTTAGTGAACCGTCAGATCGCCTGGAGACGCCATCCACGCTGTTTTGACCTCCATAGAA GATTCTAGTTCCTAGGTTCCAGTCAGGAGAGGTANNNNNNNNNNTCAGCAGGTAGGACTCAGG TGTAACACTGCAAGCGAATCGATGAAGAGTCGCTTACTCAGGCACATGCACTGCGA |

Index primers and custom read primers.
N stands for A, T, G, or C.

Supplementary Table 3: Raw data of Figure 2.

| # | | 1 | 2 | 3 | 4 | 5 | 6 | 7 | 8 | 9 | 10 | 11 | 12 | 13 | 14 | 15 | 16 | 17 | 18 | 19 | 20 |
|---|---|---|---|---|---|---|---|---|---|---|---|---|---|---|---|---|---|---|---|---|---|
| Drug | Concentration (nmol/L) | Wild-type | Empty-vector | R18H | K21R | W31C | P41L | Y42C | N55S | P59A | N108S | Q147R | D156G | V159E | P168T | V208G | F266L | R324T | S326R | S445Y | E462G |
| CBDCA | 0 | 1 | 1 | 1 | 1 | 1 | 1 | 1 | 1 | 1 | 1 | 1 | 1 | 1 | 1 | 1 | 1 | 1 | 1 | 1 | 1 |
| CBDCA | 20 | 1 | 0.9262 | 1.0886 | 0.9430 | 0.8442 | 1.4917 | 1.3256 | 0.7577 | 1.0586 | 1.0970 | 0.9481 | 1.0344 | 1.0453 | 0.9362 | 1.0262 | 1.0055 | 1.1172 | 1.0854 | 0.9602 | 1.1606 |
| CBDCA | 50 | 1 | 0.8981 | 0.9196 | 1.1034 | 0.7730 | 1.4979 | 0.9441 | 1.5449 | 1.0209 | 1.0906 | 0.9935 | 1.0275 | 0.9399 | 1.1198 | 1.3705 | 1.1557 | 1.0804 | 1.3853 | 1.0197 | 1.0640 |
| CBDCA | 100 | 1 | 0.8298 | 0.9991 | 0.9266 | 0.7707 | 1.3096 | 1.1652 | 0.8835 | 1.0718 | 1.0441 | 0.8281 | 0.9181 | 0.9882 | 0.9228 | 1.1380 | 0.9837 | 1.0009 | 1.0197 | 0.9947 | 1.1293 |
| CBDCA | 200 | 1 | 0.6058 | 0.9547 | 1.1583 | 0.5104 | 1.2308 | 0.9033 | 0.6018 | 1.1446 | 0.9445 | 0.9890 | 1.0296 | 0.8235 | 1.0756 | 1.3199 | 0.9768 | 0.8686 | 0.9422 | 0.9853 | 1.2164 |
| CBDCA | 500 | 1 | 0.1762 | 0.6108 | 1.0110 | 0.1845 | 0.7362 | 0.4758 | 0.3829 | 1.2111 | 0.6974 | 0.6798 | 1.0104 | 0.7225 | 0.9374 | 1.3403 | 0.9686 | 0.5976 | 0.7466 | 1.0276 | 1.1185 |
| CBDCA | 1000 | 1 | 0.0118 | 0.2587 | 0.8685 | 0.0114 | 0.2602 | 0.4404 | 0.1188 | 1.0876 | 0.3100 | 0.4153 | 0.8637 | 0.6096 | 0.6660 | 1.0578 | 0.5893 | 0.4229 | 0.4101 | 1.0003 | 0.6564 |
| CBDCA | 2000 | 1 | 0.0033 | 0.1112 | 0.6356 | 0.0030 | 0.2523 | 0.3592 | 0.0583 | 1.0167 | 0.3286 | 0.4688 | 1.1594 | 0.8229 | 0.5012 | 0.7311 | 0.4906 | 0.4114 | 0.2483 | 0.9627 | 0.6046 |
| Niraparib | 0 | 1 | 1 | 1 | 1 | 1 | 1 | 1 | 1 | 1 | 1 | 1 | 1 | 1 | 1 | 1 | 1 | 1 | 1 | 1 | 1 |
| Niraparib | 5 | 1 | 1.0435 | 0.9279 | 1.0829 | 0.8440 | 1.3043 | 1.4462 | 1.3518 | 1.3064 | 1.0393 | 0.9675 | 1.2271 | 1.1163 | 1.1877 | 1.0856 | 1.1080 | 1.0604 | 1.1623 | 1.0762 | 0.9798 |
| Niraparib | 10 | 1 | 0.7599 | 1.1106 | 1.1357 | 0.8751 | 1.2394 | 1.0013 | 0.8615 | 1.0618 | 1.1873 | 0.7615 | 0.9663 | 0.8656 | 1.0235 | 1.1710 | 1.0970 | 1.1488 | 0.9548 | 1.1256 | 1.0018 |
| Niraparib | 20 | 1 | 0.6853 | 0.9503 | 1.1197 | 0.5568 | 1.1840 | 1.3761 | 1.8228 | 1.0725 | 0.9442 | 0.8659 | 0.9370 | 1.0103 | 1.1765 | 1.0888 | 0.9032 | 0.9629 | 0.8272 | 1.1295 | 1.1028 |
| Niraparib | 50 | 1 | 0.4259 | 0.8165 | 1.1714 | 0.4710 | 0.9080 | 0.8804 | 0.7382 | 1.1580 | 0.9559 | 0.8437 | 1.0239 | 0.9732 | 1.1359 | 1.5907 | 1.0141 | 1.2518 | 1.0273 | 0.8449 | 1.0965 |
| Niraparib | 100 | 1 | 0.2432 | 0.6691 | 0.7792 | 0.2023 | 0.6305 | 0.6297 | 0.3559 | 0.7470 | 0.7864 | 0.5368 | 0.9319 | 0.7424 | 0.8982 | 1.2816 | 0.9892 | 0.6303 | 0.7102 | 0.8492 | 0.8076 |
| Niraparib | 200 | 1 | 0.0996 | 0.4151 | 0.9788 | 0.1240 | 0.5151 | 0.7719 | 0.4791 | 0.8348 | 0.6561 | 0.5959 | 0.8335 | 0.6106 | 0.8638 | 1.1578 | 0.6635 | 0.5181 | 0.6593 | 0.7262 | 0.8360 |
| Niraparib | 500 | 1 | 0.0060 | 0.4160 | 0.7305 | 0.0063 | 0.1671 | 0.3303 | 0.3486 | 0.8828 | 0.4289 | 0.6693 | 0.7340 | 0.4648 | 0.5495 | 1.2371 | 0.6067 | 0.4040 | 0.3054 | 0.6408 | 0.6417 |
| Olaparib | 0 | 1 | 1 | 1 | 1 | 1 | 1 | 1 | 1 | 1 | 1 | 1 | 1 | 1 | 1 | 1 | 1 | 1 | 1 | 1 | 1 |
| Olaparib | 10 | 1 | 0.6248 | 0.9221 | 1.0675 | 0.6488 | 1.1918 | 1.4548 | 0.8134 | 1.1254 | 1.0920 | 0.9814 | 1.0445 | 0.9061 | 1.2510 | 1.1473 | 0.9547 | 0.7769 | 1.1139 | 0.9654 | 1.0488 |
| Olaparib | 20 | 1 | 0.5461 | 1.1214 | 1.2045 | 0.6893 | 1.3235 | 0.8595 | 0.7675 | 0.9494 | 0.9563 | 0.9431 | 1.1814 | 1.1127 | 1.1450 | 1.2816 | 0.9612 | 0.8847 | 1.1964 | 0.9874 | 1.3610 |
| Olaparib | 50 | 1 | 0.2040 | 0.9402 | 1.0904 | 0.2874 | 1.1152 | 0.6664 | 0.5440 | 1.4013 | 0.9530 | 0.9736 | 0.9461 | 0.8793 | 1.2671 | 1.5403 | 0.8166 | 0.7693 | 0.9269 | 1.0536 | 1.2721 |
| Olaparib | 100 | 1 | 0.1558 | 0.6160 | 0.8831 | 0.2181 | 0.9128 | 0.8795 | 0.5368 | 0.9763 | 0.6224 | 0.6739 | 0.8464 | 0.6455 | 0.8860 | 0.9084 | 0.7551 | 0.6401 | 0.7510 | 0.9268 | 0.9517 |
| Olaparib | 200 | 1 | 0.0927 | 0.6857 | 1.1714 | 0.1092 | 0.8280 | 0.5942 | 0.5795 | 1.1494 | 0.5963 | 0.4447 | 0.8013 | 0.5811 | 0.9065 | 1.3906 | 0.8355 | 0.5743 | 0.8310 | 0.8163 | 0.8223 |
| Olaparib | 500 | 1 | 0.0399 | 0.4984 | 0.9462 | 0.0408 | 0.5610 | 0.4165 | 0.2299 | 0.7417 | 0.5081 | 0.2598 | 0.6757 | 0.4642 | 0.7618 | 1.0974 | 0.5759 | 0.5868 | 0.8448 | 0.6314 | 0.6538 |
| Olaparib | 1000 | 1 | 0.0185 | 0.3168 | 1.6075 | 0.0218 | 0.6109 | 0.2658 | 0.1248 | 1.2562 | 0.9810 | 0.8344 | 1.0411 | 0.4945 | 0.8707 | 1.4383 | 0.9196 | 0.5442 | 0.9064 | 1.0069 | 0.9113 |
| Olaparib | 2000 | 1 | 0.0022 | 0.3088 | 0.9695 | 0.0066 | 0.4990 | 0.0945 | 0.1918 | 0.9552 | 0.5569 | 1.1303 | 0.8180 | 0.3054 | 0.4672 | 1.4238 | 0.5906 | 0.9575 | 0.7837 | 0.9537 | 0.4580 |
| Rucaparib | 0 | 1 | 1 | 1 | 1 | 1 | 1 | 1 | 1 | 1 | 1 | 1 | 1 | 1 | 1 | 1 | 1 | 1 | 1 | 1 | 1 |
| Rucaparib | 5 | 1 | 0.6548 | 0.8822 | 1.0485 | 0.6406 | 1.0840 | 1.1006 | 0.7059 | 1.1643 | 0.9090 | 0.8131 | 1.0414 | 0.8244 | 1.0048 | 1.1526 | 1.0059 | 0.8510 | 0.9526 | 0.8663 | 1.0668 |
| Rucaparib | 10 | 1 | 0.5646 | 0.7348 | 0.9106 | 0.4899 | 1.2080 | 0.9439 | 1.0618 | 1.0328 | 0.9433 | 0.6319 | 1.0232 | 0.7852 | 0.8868 | 1.1731 | 1.0414 | 0.8208 | 0.8448 | 1.0585 | 0.9560 |
| Rucaparib | 20 | 1 | 0.4172 | 0.7008 | 1.0243 | 0.3013 | 0.8343 | 0.6984 | 0.7987 | 0.9315 | 0.7317 | 0.5914 | 0.9945 | 0.7320 | 0.9018 | 1.1635 | 0.8378 | 0.8679 | 0.8704 | 1.0587 | 1.0006 |
| Rucaparib | 50 | 1 | 0.2821 | 0.8080 | 1.0532 | 0.3089 | 0.8789 | 1.1431 | 0.5409 | 1.0554 | 0.8837 | 0.9011 | 1.0173 | 0.9469 | 1.0907 | 1.3361 | 0.8933 | 0.6899 | 0.8258 | 1.0133 | 1.0420 |
| Rucaparib | 100 | 1 | 0.1620 | 0.6007 | 1.0042 | 0.1983 | 0.7780 | 0.4594 | 0.4597 | 1.1152 | 0.7154 | 0.5703 | 0.9940 | 0.9693 | 1.0671 | 1.2672 | 0.8544 | 0.7796 | 0.9549 | 1.0210 | 0.9328 |
| Rucaparib | 200 | 1 | 0.0783 | 0.4527 | 1.0845 | 0.0778 | 0.5010 | 0.4486 | 0.3860 | 1.2263 | 0.5602 | 0.4906 | 0.9536 | 0.7172 | 1.0421 | 1.5990 | 0.7131 | 0.8722 | 0.6521 | 0.8571 | 0.8525 |
| Rucaparib | 500 | 1 | 0.0126 | 0.2997 | 0.7622 | 0.0154 | 0.3196 | 0.2241 | 0.1946 | 0.6219 | 0.4301 | 0.5641 | 0.6623 | 0.5127 | 0.5189 | 1.1140 | 0.7316 | 0.3389 | 0.6199 | 0.6535 | 0.5375 |
| Rucaparib | 1000 | 1 | 0.0072 | 0.3403 | 1.2977 | 0.0068 | 0.3004 | 0.2132 | 0.0906 | 1.0181 | 0.6356 | 0.4555 | 0.8497 | 0.3919 | 0.7497 | 1.8806 | 0.5064 | 0.4704 | 0.5281 | 0.9093 | 0.5666 |
| Rucaparib | 2000 | 1 | 0.0027 | 0.3572 | 0.8915 | 0.0022 | 0.1571 | 0.1000 | 0.1300 | 0.5355 | 0.4239 | 0.6254 | 0.6043 | 0.3833 | 0.4656 | 1.4964 | 0.5420 | 0.2679 | 0.4225 | 0.7177 | 0.6349 |

| # | | 21 | 22 | 23 | 24 | 25 | 26 | 27 | 28 | 29 | 30 | 31 | 32 | 33 | 34 | 35 | 36 | 37 | 38 | 39 | 40 |
|---|---|---|---|---|---|---|---|---|---|---|---|---|---|---|---|---|---|---|---|---|---|
| Drug | Concentration (nmol/L) | N588D | H595Y | T598A | Y600C | G602R | G602V | E747G | S755C | N854S | L1019V | S1074C | I1167V | M1168I | A1170V | S1172L | N1228D | R1329S | D1420Y | F1524V | K1530N |
| CBDCA | 0 | 1 | 1 | 1 | 1 | 1 | 1 | 1 | 1 | 1 | 1 | 1 | 1 | 1 | 1 | 1 | 1 | 1 | 1 | 1 | 1 |
| CBDCA | 20 | 0.9532 | 0.9903 | 1.0339 | 1.1265 | 1.0926 | 1.2323 | 1.0060 | 0.9848 | 1.3519 | 0.9169 | 1.1161 | 0.9978 | 1.0045 | 1.0028 | 0.9995 | 1.1441 | 1.1415 | 1.1727 | 1.2882 | 1.2088 |
| CBDCA | 50 | 0.9400 | 1.0178 | 1.0226 | 1.1428 | 1.1042 | 1.3023 | 0.8786 | 1.1068 | 1.2690 | 1.0011 | 0.9703 | 1.0277 | 0.9315 | 0.9975 | 1.2138 | 0.9847 | 1.0695 | 1.0745 | 1.2835 | 1.0506 |
| CBDCA | 100 | 0.8830 | 1.1444 | 0.8885 | 1.0009 | 1.1375 | 1.2543 | 0.9709 | 1.0535 | 1.1587 | 0.8645 | 1.0469 | 0.8962 | 0.9879 | 1.0604 | 0.9834 | 0.9196 | 1.2402 | 1.0718 | 1.0165 | 1.0065 |
| CBDCA | 200 | 0.8466 | 0.8800 | 1.0122 | 0.9670 | 1.3477 | 1.1324 | 0.9120 | 1.0138 | 1.1466 | 1.0311 | 0.9059 | 0.9310 | 1.2933 | 0.9609 | 0.8348 | 0.8920 | 1.3158 | 1.0203 | 1.0456 | 1.2473 |
| CBDCA | 500 | 0.4225 | 0.8511 | 1.0779 | 0.9807 | 1.0237 | 1.0587 | 0.9052 | 1.1548 | 1.1867 | 0.7279 | 0.7885 | 0.8128 | 0.8869 | 1.2618 | 0.6316 | 0.6433 | 1.0068 | 0.7697 | 0.5744 | 1.1904 |
| CBDCA | 1000 | 0.2960 | 0.7742 | 0.8170 | 0.8730 | 1.2684 | 0.4579 | 0.9136 | 0.6642 | 0.9056 | 0.3599 | 0.5111 | 0.6389 | 0.9630 | 1.1056 | 0.3033 | 0.2144 | 0.6620 | 0.3766 | 0.1766 | 0.7242 |
| CBDCA | 2000 | 0.3175 | 0.6857 | 0.7291 | 0.9640 | 0.6031 | 0.3417 | 1.2350 | 0.1952 | 1.5244 | 0.2536 | 0.5501 | 0.6460 | 0.9005 | 1.1748 | 0.1225 | 0.0963 | 0.5982 | 0.3577 | 0.1454 | 0.6501 |
| Niraparib | 0 | 1 | 1 | 1 | 1 | 1 | 1 | 1 | 1 | 1 | 1 | 1 | 1 | 1 | 1 | 1 | 1 | 1 | 1 | 1 | 1 |
| Niraparib | 5 | 0.8946 | 1.0174 | 1.0455 | 0.9874 | 1.3751 | 1.3842 | 0.9890 | 0.9653 | 1.0573 | 1.1397 | 1.2263 | 0.9720 | 1.1008 | 1.1276 | 0.9155 | 1.0676 | 1.1634 | 1.0137 | 1.2733 | 1.1734 |
| Niraparib | 10 | 0.7701 | 0.9099 | 0.9906 | 0.9718 | 1.1888 | 1.2220 | 0.8849 | 0.9882 | 0.9854 | 1.0433 | 1.0942 | 0.9467 | 1.0476 | 1.0713 | 0.9444 | 1.0491 | 1.0975 | 0.9802 | 1.1541 | 1.0569 |
| Niraparib | 20 | 0.8992 | 1.0910 | 1.1339 | 1.1761 | 1.3226 | 1.0755 | 0.8616 | 0.8624 | 1.1958 | 0.9618 | 0.9182 | 1.0220 | 1.1549 | 0.8871 | 0.8266 | 0.9863 | 1.0159 | 1.0805 | 0.7987 | 0.9591 |
| Niraparib | 50 | 0.7310 | 1.0789 | 1.1345 | 1.0873 | 1.3467 | 0.9386 | 0.9256 | 0.9881 | 1.1954 | 0.8724 | 1.0996 | 1.0079 | 1.2207 | 1.0498 | 0.7261 | 0.9040 | 1.1989 | 1.0724 | 1.0382 | 1.1191 |
| Niraparib | 100 | 0.5812 | 0.8034 | 1.0957 | 0.8236 | 1.3726 | 1.0394 | 0.8715 | 0.9731 | 0.9192 | 0.8194 | 0.7999 | 0.8020 | 0.9178 | 0.7557 | 0.6077 | 0.6929 | 0.9749 | 0.7971 | 0.6601 | 0.9059 |
| Niraparib | 200 | 0.3876 | 0.8823 | 1.0886 | 0.8090 | 1.4019 | 0.7920 | 0.8789 | 0.8431 | 0.8677 | 0.5364 | 0.5921 | 0.6458 | 0.8510 | 0.8776 | 0.5285 | 0.4908 | 0.8042 | 0.6740 | 0.3607 | 0.7957 |
| Niraparib | 500 | 0.1047 | 0.6012 | 0.9852 | 0.6912 | 0.7252 | 0.2616 | 0.6063 | 0.3802 | 1.2312 | 0.2463 | 0.1852 | 0.5254 | 0.9218 | 0.7550 | 0.2321 | 0.1675 | 0.7772 | 0.4136 | 0.1729 | 0.7644 |
| Olaparib | 0 | 1 | 1 | 1 | 1 | 1 | 1 | 1 | 1 | 1 | 1 | 1 | 1 | 1 | 1 | 1 | 1 | 1 | 1 | 1 | 1 |
| Olaparib | 10 | 0.9665 | 1.1573 | 1.1035 | 0.9519 | 1.1192 | 1.3902 | 1.0062 | 0.9669 | 1.2903 | 0.8155 | 0.9674 | 0.9928 | 0.9937 | 1.1270 | 0.7920 | 1.0046 | 1.1192 | 0.9106 | 1.1160 | 1.2827 |
| Olaparib | 20 | 0.7677 | 1.0236 | 1.2432 | 1.0663 | 1.4929 | 1.1442 | 0.8961 | 1.1149 | 1.1672 | 1.0882 | 0.9668 | 1.1164 | 0.9333 | 1.1085 | 0.7981 | 1.0272 | 1.1662 | 0.9432 | 1.0832 | 1.0773 |
| Olaparib | 50 | 0.8131 | 1.3232 | 1.4150 | 1.1127 | 1.7143 | 1.2155 | 1.0774 | 1.1135 | 1.1987 | 1.0723 | 0.9936 | 1.1292 | 1.2076 | 1.1815 | 1.0040 | 0.9447 | 1.2225 | 0.9897 | 0.8714 | 1.2402 |
| Olaparib | 100 | 0.4520 | 1.0721 | 1.2841 | 0.8704 | 1.0578 | 0.9248 | 0.9602 | 0.7552 | 0.9923 | 0.6002 | 0.6812 | 0.7639 | 1.1624 | 0.9185 | 0.6171 | 0.6288 | 0.8063 | 0.7169 | 0.4581 | 1.0234 |
| Olaparib | 200 | 0.5093 | 0.9091 | 1.3012 | 0.9414 | 1.2192 | 0.9431 | 0.8830 | 0.7433 | 0.9873 | 0.6714 | 0.6311 | 0.6886 | 0.8591 | 1.2083 | 0.4727 | 0.5719 | 0.9059 | 0.7189 | 0.5324 | 1.0901 |
| Olaparib | 500 | 0.2892 | 0.8124 | 0.9465 | 0.6479 | 0.9503 | 0.5769 | 1.0079 | 0.3939 | 0.8861 | 0.4461 | 0.4468 | 0.6162 | 1.1430 | 0.7019 | 0.3857 | 0.2810 | 0.7231 | 0.4342 | 0.3351 | 0.6857 |
| Olaparib | 1000 | 0.2484 | 1.1979 | 1.5945 | 0.7442 | 1.8672 | 0.5512 | 1.3269 | 0.2877 | 1.1358 | 0.3775 | 0.5055 | 0.8280 | 1.1702 | 1.1858 | 0.2754 | 0.3044 | 0.6687 | 0.4060 | 0.2618 | 1.1602 |
| Olaparib | 2000 | 0.2073 | 0.9613 | 1.1831 | 0.7394 | 1.2704 | 0.4836 | 1.0814 | 0.2393 | 0.8949 | 0.1502 | 0.3706 | 0.6900 | 1.5491 | 0.7985 | 0.2199 | 0.1635 | 0.5476 | 0.3256 | 0.1922 | 0.6121 |
| Rucaparib | 0 | 1 | 1 | 1 | 1 | 1 | 1 | 1 | 1 | 1 | 1 | 1 | 1 | 1 | 1 | 1 | 1 | 1 | 1 | 1 | 1 |
| Rucaparib | 5 | 0.8325 | 1.0533 | 1.0351 | 0.8897 | 1.2392 | 1.1553 | 0.9812 | 0.8422 | 0.9537 | 0.9520 | 0.9387 | 0.8675 | 0.9978 | 1.1358 | 0.7679 | 0.8327 | 1.1342 | 1.0150 | 1.0371 | 1.0998 |
| Rucaparib | 10 | 0.7906 | 0.9529 | 1.0355 | 0.8710 | 1.0941 | 0.8375 | 0.8491 | 0.8741 | 0.9335 | 1.0293 | 0.8802 | 0.7827 | 0.9392 | 0.9525 | 0.7753 | 0.9053 | 0.9667 | 0.7767 | 0.8886 | 0.8108 |
| Rucaparib | 20 | 0.6274 | 0.9036 | 0.9709 | 0.8729 | 1.1198 | 1.0101 | 0.8524 | 1.0082 | 1.3118 | 0.9335 | 0.8788 | 0.7465 | 0.9238 | 0.8674 | 0.7573 | 0.8412 | 1.0102 | 0.8832 | 0.8594 | 0.9558 |
| Rucaparib | 50 | 0.7364 | 1.0768 | 1.2936 | 0.8572 | 1.7208 | 1.1200 | 0.9352 | 0.9227 | 1.1677 | 0.8452 | 0.8648 | 0.9071 | 1.1342 | 1.0179 | 0.7364 | 0.5568 | 1.1026 | 0.9151 | 0.6886 | 1.0516 |
| Rucaparib | 100 | 0.3592 | 0.9565 | 1.1849 | 0.8880 | 1.2576 | 1.1402 | 0.8472 | 0.8283 | 1.0850 | 0.8211 | 0.9746 | 0.8199 | 1.1172 | 1.1476 | 0.6240 | 0.7559 | 1.2157 | 1.0722 | 0.6391 | 1.0278 |
| Rucaparib | 200 | 0.3876 | 0.8887 | 1.3010 | 0.8364 | 1.2711 | 0.9815 | 0.8882 | 0.7813 | 1.3198 | 0.6178 | 0.6898 | 0.6624 | 0.9451 | 1.0764 | 0.5990 | 0.5881 | 0.8312 | 0.7466 | 0.4430 | 1.3954 |
| Rucaparib | 500 | 0.2833 | 0.6946 | 0.7717 | 0.5907 | 1.1575 | 0.3924 | 0.8449 | 0.4548 | 1.0362 | 0.2963 | 0.3848 | 0.5376 | 0.7353 | 0.7930 | 0.2363 | 0.2487 | 0.4857 | 0.3070 | 0.1177 | 0.6564 |
| Rucaparib | 1000 | 0.3320 | 0.6653 | 1.3455 | 0.9682 | 1.3539 | 0.4495 | 0.9691 | 0.4505 | 0.9772 | 0.4094 | 0.5379 | 0.7615 | 1.0301 | 1.1584 | 0.3354 | 0.2174 | 0.6376 | 0.4798 | 0.2897 | 1.1967 |
| Rucaparib | 2000 | 0.1469 | 0.7816 | 1.3496 | 0.6166 | 0.9812 | 0.2481 | 0.8668 | 0.1935 | 1.1672 | 0.3328 | 0.2985 | 0.5142 | 1.4058 | 0.9003 | 0.1144 | 0.1248 | 0.6347 | 0.2821 | 0.1366 | 0.6765 |

| # | | 41 | 42 | 43 | 44 | 45 | 46 | 47 | 48 | 49 | 50 | 51 | 52 | 53 | 54 | 55 | 56 | 57 | 58 | 59 | 60 |
|---|---|---|---|---|---|---|---|---|---|---|---|---|---|---|---|---|---|---|---|---|---|
| Drug | Concentration (nmol/L) | V1532F | K1690N | E1695V | G1696V | D1699N | D1728N | D1737V | N1878K | H1918R | S1970L | E2020K | H2074N | K2075N | V2138D | H2324R | P2329L | R2336G | T2337A | K2411T | T2412A |
| CBDCA | 0 | 1 | 1 | 1 | 1 | 1 | 1 | 1 | 1 | 1 | 1 | 1 | 1 | 1 | 1 | 1 | 1 | 1 | 1 | 1 | 1 |
| CBDCA | 20 | 1.2297 | 1.2095 | 0.9898 | 1.0271 | 0.9838 | 1.0130 | 1.3348 | 1.0483 | 0.9858 | 1.1400 | 1.2544 | 1.1874 | 1.0206 | 0.9449 | 1.1005 | 1.1557 | 1.1685 | 1.1256 | 0.9445 | 1.2095 |
| CBDCA | 50 | 1.2033 | 1.1299 | 0.9777 | 1.0723 | 0.9941 | 1.0437 | 1.2517 | 1.0262 | 1.1314 | 1.0473 | 1.1461 | 1.1065 | 1.0124 | 0.9328 | 1.1824 | 0.9501 | 1.2581 | 1.1219 | 1.0078 | 1.1740 |
| CBDCA | 100 | 0.9367 | 0.9101 | 1.0363 | 0.8509 | 0.9505 | 0.7838 | 1.1059 | 1.0282 | 0.9357 | 0.9163 | 1.2034 | 1.0806 | 0.9273 | 0.9161 | 1.0302 | 1.1239 | 1.0497 | 0.9619 | 0.9135 | 1.1230 |
| CBDCA | 200 | 1.0846 | 1.7679 | 0.9695 | 0.6768 | 0.8045 | 0.7872 | 1.1422 | 0.9824 | 0.8616 | 1.0222 | 1.0604 | 1.0530 | 0.8680 | 0.8837 | 0.7815 | 0.8805 | 0.9533 | 0.9423 | 0.9671 | 1.0504 |
| CBDCA | 500 | 0.8407 | 1.6749 | 0.6482 | 0.5230 | 0.6481 | 0.9255 | 0.9822 | 0.7919 | 0.5828 | 1.0358 | 0.9312 | 1.0581 | 0.7308 | 0.7590 | 0.4961 | 0.3775 | 1.3081 | 0.5407 | 0.6496 | 1.0139 |
| CBDCA | 1000 | 1.0021 | 1.9824 | 0.4795 | 0.1004 | 0.6600 | 0.6158 | 0.8658 | 0.5069 | 0.2618 | 0.7223 | 0.6005 | 0.8018 | 0.3604 | 0.5450 | 0.2636 | 0.0630 | 1.3357 | 0.2651 | 0.3492 | 0.5984 |
| CBDCA | 2000 | 1.5515 | 2.8214 | 0.3141 | 0.0946 | 0.3065 | 0.2364 | 0.7007 | 0.4199 | 0.2092 | 0.4085 | 0.6030 | 0.8925 | 0.2086 | 0.4668 | 0.1665 | 0.0163 | 2.5608 | 0.1709 | 0.1603 | 0.5526 |
| Niraparib | 0 | 1 | 1 | 1 | 1 | 1 | 1 | 1 | 1 | 1 | 1 | 1 | 1 | 1 | 1 | 1 | 1 | 1 | 1 | 1 | 1 |
| Niraparib | 5 | 1.3074 | 1.0750 | 1.0989 | 0.9298 | 1.1351 | 1.1087 | 1.1130 | 1.0047 | 1.0284 | 1.0682 | 1.1833 | 1.2289 | 1.0716 | 1.0715 | 1.0632 | 0.9922 | 1.0455 | 1.1617 | 1.0245 | 1.3369 |
| Niraparib | 10 | 1.1912 | 1.2727 | 1.0994 | 0.8593 | 0.9906 | 0.7548 | 1.0771 | 1.0044 | 1.0144 | 0.9474 | 1.0036 | 1.2777 | 1.0060 | 0.8688 | 0.9978 | 0.9177 | 1.1085 | 1.2752 | 0.9982 | 1.2924 |
| Niraparib | 20 | 0.8049 | 1.2092 | 0.9295 | 0.6819 | 1.0572 | 0.5683 | 0.8556 | 1.0894 | 0.7599 | 0.9022 | 0.8343 | 1.2363 | 1.0229 | 1.1171 | 0.9744 | 0.8839 | 1.1314 | 1.0851 | 0.8205 | 1.4051 |
| Niraparib | 50 | 1.1412 | 1.6586 | 1.1082 | 0.7890 | 0.9804 | 0.9267 | 1.0677 | 1.0560 | 0.7936 | 1.1944 | 1.0156 | 1.1922 | 1.0480 | 1.0678 | 0.7532 | 0.5043 | 1.3708 | 0.8526 | 0.8537 | 1.1431 |
| Niraparib | 100 | 0.7170 | 1.2639 | 0.6841 | 0.5190 | 0.8891 | 0.6724 | 0.7766 | 0.8069 | 0.6140 | 0.8943 | 0.7267 | 1.1788 | 0.6543 | 0.6326 | 0.6183 | 0.4536 | 1.0892 | 0.7704 | 0.6049 | 1.0787 |
| Niraparib | 200 | 0.6807 | 1.5872 | 0.6278 | 0.4250 | 0.8412 | 0.8638 | 0.6517 | 0.8047 | 0.4216 | 1.0439 | 0.6168 | 1.0451 | 0.5346 | 0.6503 | 0.4493 | 0.2676 | 1.0855 | 0.4979 | 0.5432 | 0.8044 |
| Niraparib | 500 | 0.5213 | 2.1557 | 0.3575 | 0.2071 | 0.5109 | 1.1131 | 0.6509 | 0.5916 | 0.1437 | 0.7503 | 0.2712 | 0.6498 | 0.3989 | 0.2736 | 0.1737 | 0.0421 | 1.4377 | 0.1570 | 0.2989 | 0.6970 |
| Olaparib | 0 | 1 | 1 | 1 | 1 | 1 | 1 | 1 | 1 | 1 | 1 | 1 | 1 | 1 | 1 | 1 | 1 | 1 | 1 | 1 | 1 |
| Olaparib | 10 | 1.2394 | 1.2430 | 1.0006 | 0.7941 | 0.8295 | 0.6800 | 1.2263 | 1.2633 | 0.8843 | 1.2045 | 1.1462 | 1.2971 | 0.8690 | 0.9351 | 0.9588 | 1.0163 | 0.9362 | 1.1761 | 0.8790 | 1.1958 |
| Olaparib | 20 | 1.3927 | 1.5145 | 0.9748 | 0.8789 | 1.0030 | 1.1596 | 1.1821 | 1.1548 | 1.2010 | 1.2558 | 1.0252 | 1.3681 | 0.9973 | 0.8348 | 0.8595 | 0.8872 | 1.2419 | 1.1317 | 0.9575 | 1.3706 |
| Olaparib | 50 | 1.0212 | 1.9709 | 0.8487 | 0.7193 | 1.2516 | 0.9278 | 1.3358 | 1.1534 | 0.9031 | 1.2864 | 0.9703 | 1.7079 | 1.2857 | 1.0316 | 0.7112 | 0.5336 | 1.0142 | 1.1361 | 0.9689 | 1.2511 |
| Olaparib | 100 | 1.0971 | 1.3038 | 0.7411 | 0.4534 | 0.6069 | 0.7368 | 0.8508 | 0.7466 | 0.6954 | 0.7029 | 0.6589 | 0.9502 | 0.5668 | 0.6328 | 0.5757 | 0.3844 | 0.8979 | 0.8056 | 0.6224 | 0.9573 |
| Olaparib | 200 | 0.8535 | 1.4766 | 0.4829 | 0.4060 | 0.8844 | 0.9586 | 0.8310 | 0.8836 | 0.5580 | 1.0499 | 0.8290 | 1.0722 | 0.9701 | 0.6790 | 0.4271 | 0.2524 | 1.3179 | 0.7832 | 0.6091 | 1.0898 |
| Olaparib | 500 | 1.0710 | 1.2875 | 0.2885 | 0.2862 | 0.5276 | 1.0474 | 0.7803 | 0.6343 | 0.3395 | 0.7481 | 0.6690 | 0.7529 | 0.5410 | 0.5082 | 0.2333 | 0.0868 | 0.8179 | 0.3196 | 0.3664 | 0.9449 |
| Olaparib | 1000 | 1.7125 | 2.2378 | 0.5250 | 0.2129 | 0.7517 | 0.9398 | 1.1704 | 0.5815 | 0.4462 | 0.9296 | 0.6022 | 1.2982 | 0.6782 | 0.5351 | 0.1222 | 0.0874 | 1.6152 | 0.5285 | 0.3602 | 0.9020 |
| Olaparib | 2000 | 1.0042 | 1.4466 | 0.6850 | 0.2310 | 0.4611 | 0.9947 | 0.8460 | 0.3936 | 0.1639 | 0.4300 | 0.7043 | 0.6692 | 0.4896 | 0.4531 | 0.1384 | 0.0173 | 1.7954 | 0.2430 | 0.1902 | 0.6730 |
| Rucaparib | 0 | 1 | 1 | 1 | 1 | 1 | 1 | 1 | 1 | 1 | 1 | 1 | 1 | 1 | 1 | 1 | 1 | 1 | 1 | 1 | 1 |
| Rucaparib | 5 | 1.0166 | 1.0855 | 0.8919 | 0.7213 | 0.8921 | 0.8198 | 0.9919 | 0.9589 | 0.9163 | 1.0168 | 1.0825 | 1.1193 | 0.9098 | 0.8820 | 0.8994 | 0.9299 | 0.8879 | 1.0218 | 1.0790 | 1.1529 |
| Rucaparib | 10 | 0.9640 | 1.2045 | 0.7398 | 0.7596 | 0.8577 | 0.6104 | 1.1287 | 0.9946 | 0.8096 | 0.9272 | 1.1820 | 1.0962 | 0.7739 | 0.9921 | 0.8658 | 0.6374 | 0.9869 | 0.7948 | 0.9124 | 0.9631 |
| Rucaparib | 20 | 0.7271 | 1.4050 | 0.6765 | 0.5542 | 0.6982 | 0.5954 | 0.8522 | 0.7758 | 0.6810 | 0.9954 | 1.0455 | 1.0048 | 0.8139 | 0.7601 | 0.7579 | 0.6036 | 0.8295 | 0.7245 | 0.7634 | 0.8690 |
| Rucaparib | 50 | 1.0683 | 1.7520 | 0.6224 | 0.6149 | 0.8117 | 0.6710 | 0.8956 | 1.0138 | 0.6322 | 1.0000 | 0.6628 | 1.1507 | 0.9763 | 0.8741 | 0.7335 | 0.6246 | 1.1712 | 0.9029 | 0.8124 | 1.1721 |
| Rucaparib | 100 | 0.8712 | 1.4929 | 0.5857 | 0.4675 | 0.8260 | 0.9194 | 0.9326 | 1.0002 | 0.5981 | 0.8801 | 0.8714 | 1.1244 | 0.6753 | 0.8176 | 0.4728 | 0.3397 | 1.0731 | 0.6966 | 0.6050 | 1.1177 |
| Rucaparib | 200 | 0.9496 | 1.7119 | 0.6129 | 0.4599 | 0.7373 | 1.0662 | 0.9622 | 0.8350 | 0.4760 | 0.9616 | 0.6942 | 1.2111 | 0.6671 | 0.6876 | 0.5121 | 0.2268 | 1.0797 | 0.5042 | 0.6864 | 0.9405 |
| Rucaparib | 500 | 0.6074 | 1.4056 | 0.4757 | 0.1792 | 0.5653 | 0.7419 | 0.6115 | 0.5681 | 0.2718 | 0.5806 | 0.4897 | 0.6925 | 0.2875 | 0.4876 | 0.2159 | 0.0431 | 1.1889 | 0.3082 | 0.2350 | 0.6707 |
| Rucaparib | 1000 | 0.8318 | 2.1014 | 0.5170 | 0.1804 | 0.8474 | 1.3114 | 0.7423 | 0.7700 | 0.3615 | 0.7846 | 0.6480 | 0.9716 | 0.4858 | 0.4974 | 0.2266 | 0.0369 | 1.7062 | 0.3030 | 0.5415 | 0.7985 |
| Rucaparib | 2000 | 0.3744 | 2.3526 | 0.4105 | 0.1058 | 0.6752 | 0.9412 | 0.2867 | 0.3438 | 0.0769 | 0.3869 | 0.3071 | 0.6351 | 0.5395 | 0.3519 | 0.1132 | 0.0156 | 1.4326 | 0.1212 | 0.1512 | 0.7618 |

| # | | 61 | 62 | 63 | 64 | 65 | 66 | 67 | 68 | 69 | 70 | 71 | 72 | 73 | 74 | 75 | 76 | 77 | 78 | 79 | 80 |
|---|---|---|---|---|---|---|---|---|---|---|---|---|---|---|---|---|---|---|---|---|---|
| Drug | Concentration (nmol/L) | T2412I | R2418G | N2436I | Y2601C | L2604P | G2609D | N2622S | H2623R | W2626R | W2626C | I2627V | P2639L | E2663K | I2664M | M2676T | L2688P | S2697N | A2717S | L2721H | T2722R |
| CBDCA | 0 | 1 | 1 | 1 | 1 | 1 | 1 | 1 | 1 | 1 | 1 | 1 | 1 | 1 | 1 | 1 | 1 | 1 | 1 | 1 | 1 |
| CBDCA | 20 | 1.0151 | 0.9735 | 0.9359 | 1.2078 | 1.1132 | 1.0210 | 0.9299 | 0.7739 | 1.4371 | 0.8242 | 1.0250 | 1.1965 | 0.8612 | 1.3652 | 1.1130 | 1.0017 | 0.9968 | 1.0337 | 0.8794 | 0.9791 |
| CBDCA | 50 | 1.2265 | 1.0013 | 0.8944 | 1.2310 | 1.0225 | 0.7230 | 0.8963 | 0.8778 | 0.9645 | 1.0030 | 1.0701 | 1.0746 | 1.0892 | 1.3236 | 1.1300 | 1.1656 | 1.3038 | 1.0614 | 0.8047 | 1.3766 |
| CBDCA | 100 | 1.0310 | 0.8621 | 0.9350 | 1.0685 | 0.7933 | 0.6831 | 0.8732 | 0.7018 | 0.9929 | 0.8770 | 0.9175 | 0.9475 | 0.9671 | 1.0891 | 1.0711 | 0.7568 | 1.3715 | 0.9906 | 0.8146 | 1.0797 |
| CBDCA | 200 | 0.9068 | 0.7916 | 0.9742 | 0.8385 | 0.4651 | 0.6041 | 0.4505 | 0.4256 | 0.6351 | 0.5403 | 0.9897 | 0.8448 | 0.4745 | 0.8429 | 1.0644 | 0.6241 | 1.3937 | 0.8709 | 0.4558 | 0.6379 |
| CBDCA | 500 | 0.8211 | 0.5946 | 0.7287 | 0.4848 | 0.1480 | 0.1634 | 0.1735 | 0.1203 | 0.2490 | 0.0780 | 0.7014 | 0.3443 | 0.1594 | 0.4956 | 0.8418 | 0.1395 | 1.1652 | 0.8057 | 0.1144 | 0.2230 |
| CBDCA | 1000 | 0.2866 | 0.2068 | 0.3788 | 0.2769 | 0.0165 | 0.0267 | 0.0124 | 0.0106 | 0.0299 | 0.0240 | 0.4980 | 0.0642 | 0.0094 | 0.1453 | 0.3623 | 0.0084 | 0.9394 | 0.6922 | 0.0080 | 0.0233 |
| CBDCA | 2000 | 0.2929 | 0.0844 | 0.2701 | 0.1266 | 0.0044 | 0.0055 | 0.0074 | 0.0054 | 0.0094 | 0.0091 | 0.2757 | 0.0109 | 0.0063 | 0.1537 | 0.2111 | 0.0105 | 1.1133 | 0.3434 | 0.0064 | 0.0052 |
| Niraparib | 0 | 1 | 1 | 1 | 1 | 1 | 1 | 1 | 1 | 1 | 1 | 1 | 1 | 1 | 1 | 1 | 1 | 1 | 1 | 1 | 1 |
| Niraparib | 5 | 1.2850 | 0.9778 | 0.9984 | 0.9988 | 0.9699 | 0.8068 | 0.8027 | 0.6901 | 0.8460 | 1.2116 | 0.8976 | 1.0683 | 1.0215 | 1.1247 | 1.1237 | 1.1500 | 1.1592 | 1.2758 | 0.7942 | 0.9911 |
| Niraparib | 10 | 1.1116 | 0.9160 | 0.8448 | 1.1602 | 0.7696 | 0.7327 | 0.7312 | 0.7255 | 0.7087 | 0.6628 | 1.0882 | 0.9830 | 0.9593 | 1.0481 | 1.1296 | 0.8743 | 1.0893 | 1.0657 | 0.6907 | 0.8706 |
| Niraparib | 20 | 0.8862 | 0.8127 | 0.9074 | 0.9074 | 0.6708 | 0.7367 | 0.5606 | 0.5928 | 0.7373 | 0.5369 | 1.0928 | 0.7952 | 0.7013 | 0.8331 | 1.1097 | 0.5924 | 1.2378 | 0.9283 | 0.4895 | 0.9797 |
| Niraparib | 50 | 0.9814 | 0.8386 | 1.0289 | 0.9016 | 0.5186 | 0.4307 | 0.3333 | 0.2326 | 0.4267 | 0.4727 | 1.0031 | 0.6957 | 0.2885 | 0.7480 | 1.0344 | 0.3196 | 1.1956 | 0.9392 | 0.3221 | 0.5805 |
| Niraparib | 100 | 0.7214 | 0.4781 | 0.8108 | 0.6602 | 0.2262 | 0.2521 | 0.1433 | 0.2261 | 0.2653 | 0.1848 | 0.9112 | 0.4718 | 0.2119 | 0.4987 | 0.7693 | 0.2009 | 1.4771 | 0.7000 | 0.2067 | 0.3203 |
| Niraparib | 200 | 0.8093 | 0.4157 | 0.6588 | 0.3417 | 0.0822 | 0.1824 | 0.0576 | 0.0428 | 0.0785 | 0.0828 | 0.9148 | 0.2329 | 0.0469 | 0.4433 | 0.5485 | 0.1175 | 1.1111 | 0.6834 | 0.0572 | 0.1180 |
| Niraparib | 500 | 0.4431 | 0.2370 | 0.2812 | 0.0617 | 0.0067 | 0.0043 | 0.0205 | 0.0029 | 0.0066 | 0.0032 | 0.3454 | 0.0593 | 0.0046 | 0.2397 | 0.2948 | 0.0060 | 0.9427 | 0.4080 | 0.0015 | 0.0112 |
| Olaparib | 0 | 1 | 1 | 1 | 1 | 1 | 1 | 1 | 1 | 1 | 1 | 1 | 1 | 1 | 1 | 1 | 1 | 1 | 1 | 1 | 1 |
| Olaparib | 10 | 1.0803 | 0.9228 | 1.0446 | 1.1447 | 0.7217 | 0.6389 | 0.5281 | 0.6006 | 0.7262 | 0.5123 | 1.0449 | 0.8025 | 0.5847 | 1.0050 | 1.0041 | 0.5468 | 1.0918 | 0.9746 | 0.4896 | 0.7086 |
| Olaparib | 20 | 1.1579 | 0.7773 | 1.0662 | 0.7481 | 0.5333 | 0.6131 | 0.5732 | 0.3610 | 0.8436 | 0.5404 | 1.4743 | 0.7488 | 0.4037 | 0.8379 | 0.9059 | 0.3153 | 1.2086 | 0.9878 | 0.3110 | 0.6271 |
| Olaparib | 50 | 0.9708 | 0.7284 | 0.9239 | 0.9138 | 0.2455 | 0.3304 | 0.3113 | 0.1994 | 0.2405 | 0.2191 | 1.3666 | 0.4981 | 0.1352 | 0.5482 | 0.7046 | 0.2170 | 1.4370 | 0.9582 | 0.2019 | 0.4545 |
| Olaparib | 100 | 0.7428 | 0.5141 | 0.6852 | 0.6501 | 0.1046 | 0.2360 | 0.1387 | 0.1322 | 0.1626 | 0.1110 | 0.8628 | 0.2543 | 0.1461 | 0.4394 | 0.7766 | 0.1603 | 1.1813 | 0.7878 | 0.1482 | 0.1995 |
| Olaparib | 200 | 0.8726 | 0.4366 | 0.7486 | 0.3729 | 0.1384 | 0.2094 | 0.0706 | 0.1237 | 0.0755 | 0.1106 | 0.6373 | 0.2295 | 0.1277 | 0.4035 | 0.7136 | 0.0607 | 1.1388 | 0.6775 | 0.0481 | 0.1145 |
| Olaparib | 500 | 0.4330 | 0.2417 | 0.5318 | 0.2216 | 0.0300 | 0.1550 | 0.0055 | 0.0240 | 0.0469 | 0.0227 | 0.3530 | 0.1002 | 0.0082 | 0.3534 | 0.5148 | 0.0290 | 1.0327 | 0.4508 | 0.0055 | 0.0140 |
| Olaparib | 1000 | 0.4023 | 0.3973 | 0.5020 | 0.2757 | 0.0055 | 0.0410 | 0.0283 | 0.0068 | 0.0195 | 0.0157 | 0.7908 | 0.0672 | 0.0167 | 0.2802 | 0.5153 | 0.0062 | 1.0574 | 0.9731 | 0.0047 | 0.0193 |
| Olaparib | 2000 | 0.3182 | 0.2369 | 0.2868 | 0.1827 | 0.0097 | 0.0094 | 0.0043 | 0.0037 | 0.0118 | 0.0068 | 0.6939 | 0.0143 | 0.0161 | 0.1043 | 0.2715 | 0.0048 | 0.7665 | 0.9938 | 0.0074 | 0.0032 |
| Rucaparib | 0 | 1 | 1 | 1 | 1 | 1 | 1 | 1 | 1 | 1 | 1 | 1 | 1 | 1 | 1 | 1 | 1 | 1 | 1 | 1 | 1 |
| Rucaparib | 5 | 1.1945 | 0.9002 | 0.9749 | 1.0282 | 0.6818 | 0.7027 | 0.5199 | 0.4887 | 0.5608 | 0.6124 | 1.1390 | 0.8456 | 0.6675 | 0.8801 | 1.0151 | 0.6891 | 0.9104 | 1.0576 | 0.5309 | 0.8211 |
| Rucaparib | 10 | 0.9035 | 0.5966 | 0.8656 | 0.8738 | 0.5135 | 0.4556 | 0.4624 | 0.3736 | 0.4575 | 0.4859 | 0.7752 | 0.6246 | 0.5590 | 0.7336 | 0.9042 | 0.3946 | 0.9954 | 0.8504 | 0.3526 | 0.6039 |
| Rucaparib | 20 | 0.8833 | 0.6388 | 0.8605 | 1.1434 | 0.4668 | 0.3873 | 0.4259 | 0.2805 | 0.2872 | 0.3897 | 1.0566 | 0.5837 | 0.3137 | 0.6645 | 0.9422 | 0.3585 | 1.1106 | 0.7287 | 0.2792 | 0.5678 |
| Rucaparib | 50 | 1.0305 | 0.5900 | 0.9915 | 0.7667 | 0.2592 | 0.3136 | 0.1695 | 0.1639 | 0.1953 | 0.3222 | 1.0409 | 0.5604 | 0.2091 | 0.5394 | 0.8726 | 0.1172 | 1.6921 | 0.7707 | 0.1689 | 0.3775 |
| Rucaparib | 100 | 0.7829 | 0.6118 | 0.9195 | 0.5837 | 0.1627 | 0.2725 | 0.1571 | 0.0807 | 0.1108 | 0.1454 | 0.8966 | 0.3646 | 0.1009 | 0.4997 | 0.6581 | 0.1271 | 1.2216 | 0.8320 | 0.0809 | 0.1844 |
| Rucaparib | 200 | 0.6769 | 0.4263 | 0.7031 | 0.3686 | 0.0638 | 0.1352 | 0.0623 | 0.0648 | 0.0630 | 0.0675 | 0.7881 | 0.2174 | 0.0630 | 0.4794 | 0.6764 | 0.0902 | 1.3830 | 0.8940 | 0.0364 | 0.0784 |
| Rucaparib | 500 | 0.3390 | 0.2227 | 0.4515 | 0.1583 | 0.0175 | 0.0224 | 0.0238 | 0.0152 | 0.0050 | 0.0089 | 0.3800 | 0.0639 | 0.0183 | 0.1451 | 0.4218 | 0.0060 | 0.8129 | 0.5480 | 0.0098 | 0.0101 |
| Rucaparib | 1000 | 0.4272 | 0.3601 | 0.4340 | 0.0933 | 0.0090 | 0.0182 | 0.0192 | 0.0117 | 0.0085 | 0.0114 | 0.4355 | 0.0528 | 0.0112 | 0.1923 | 0.4960 | 0.0071 | 1.4726 | 0.9085 | 0.0061 | 0.0152 |
| Rucaparib | 2000 | 0.5072 | 0.1776 | 0.1826 | 0.0715 | 0.0018 | 0.0017 | 0.0038 | 0.0030 | 0.0146 | 0.0038 | 0.3404 | 0.0116 | 0.0062 | 0.2468 | 0.3296 | 0.0049 | 1.5160 | 0.6585 | 0.0054 | 0.0025 |

| # | | 81 | 82 | 83 | 84 | 85 | 86 | 87 | 88 | 89 | 90 | 91 | 92 | 93 | 94 | 95 | 96 | 97 | 98 | 99 | 100 |
|---|---|---|---|---|---|---|---|---|---|---|---|---|---|---|---|---|---|---|---|---|---|
| Drug | Concentration (nmol/L) | T2722K | D2723H | D2723E | G2724W | K2729N | V2747I | G2748D | R2842H | R2896C | V2908G | A2911E | A2911V | D2913H | I2944F | I3044V | R3052W | P3054S | D3095E | E3096K | Y3098H |
| CBDCA | 0 | 1 | 1 | 1 | 1 | 1 | 1 | 1 | 1 | 1 | 1 | 1 | 1 | 1 | 1 | 1 | 1 | 1 | 1 | 1 | 1 |
| CBDCA | 20 | 0.9769 | 0.8412 | 0.7827 | 1.0855 | 1.0437 | 1.0374 | 0.9856 | 0.9003 | 1.1447 | 1.2407 | 1.4134 | 1.0132 | 0.9892 | 1.0262 | 1.0576 | 1.0378 | 0.8697 | 0.8338 | 1.0733 | 1.1680 |
| CBDCA | 50 | 0.9646 | 0.9507 | 1.0141 | 1.1323 | 0.8584 | 1.0995 | 0.9435 | 0.5977 | 1.2473 | 1.1818 | 1.9062 | 0.8775 | 0.9535 | 1.0249 | 1.1411 | 1.0924 | 0.9788 | 0.8578 | 0.9906 | 1.0210 |
| CBDCA | 100 | 0.7931 | 0.8114 | 0.9666 | 0.6581 | 0.7834 | 0.9774 | 0.9446 | 0.6126 | 1.2345 | 0.9207 | 1.4559 | 0.8959 | 0.7420 | 0.9849 | 1.0797 | 0.6291 | 0.9616 | 0.7326 | 1.0249 | 1.2845 |
| CBDCA | 200 | 0.5682 | 0.5489 | 0.5939 | 0.5630 | 0.9778 | 1.0260 | 0.5800 | 0.4436 | 1.3674 | 0.8638 | 1.9014 | 0.9439 | 0.5803 | 0.8948 | 0.9863 | 0.4547 | 0.8557 | 0.4419 | 0.8899 | 1.1806 |
| CBDCA | 500 | 0.1402 | 0.1361 | 0.3148 | 0.1401 | 1.2829 | 0.7371 | 0.1755 | 0.1985 | 1.3664 | 0.4233 | 1.7178 | 0.6888 | 0.1987 | 0.7175 | 1.1917 | 0.0916 | 0.6534 | 0.1212 | 1.0933 | 1.3957 |
| CBDCA | 1000 | 0.0065 | 0.0216 | 0.0229 | 0.0094 | 0.8836 | 0.3610 | 0.0067 | 0.0364 | 1.8102 | 0.0981 | 1.5230 | 0.2398 | 0.0703 | 0.3331 | 0.7952 | 0.0067 | 0.2474 | 0.0084 | 0.7550 | 1.5639 |
| CBDCA | 2000 | 0.0034 | 0.0025 | 0.0033 | 0.0056 | 1.6746 | 0.2372 | 0.0055 | 0.0052 | 2.9358 | 0.0071 | 1.2057 | 0.1451 | 0.0257 | 0.1790 | 0.6495 | 0.0032 | 0.1707 | 0.0039 | 0.9144 | 2.2778 |
| Niraparib | 0 | 1 | 1 | 1 | 1 | 1 | 1 | 1 | 1 | 1 | 1 | 1 | 1 | 1 | 1 | 1 | 1 | 1 | 1 | 1 | 1 |
| Niraparib | 5 | 0.7779 | 0.7052 | 1.0213 | 0.8913 | 0.8875 | 1.1329 | 0.9712 | 0.7975 | 1.2787 | 1.1836 | 1.4022 | 1.0537 | 0.8081 | 1.0634 | 1.1173 | 0.9088 | 1.0078 | 0.7048 | 1.2315 | 1.1536 |
| Niraparib | 10 | 0.7687 | 0.6647 | 0.9457 | 0.8274 | 0.7996 | 1.0572 | 0.9216 | 0.6818 | 1.1240 | 1.0301 | 1.5235 | 0.8768 | 0.8258 | 0.9076 | 1.0298 | 0.6022 | 0.9402 | 0.7464 | 0.9666 | 1.0422 |
| Niraparib | 20 | 0.6473 | 0.6970 | 0.7095 | 0.6301 | 0.9500 | 0.9574 | 0.7326 | 0.8527 | 1.4284 | 0.9202 | 1.7207 | 1.0551 | 0.7787 | 1.1118 | 1.3110 | 0.5672 | 0.7802 | 0.7324 | 1.1494 | 1.5789 |
| Niraparib | 50 | 0.3429 | 0.3628 | 0.4183 | 0.3659 | 1.4179 | 1.1922 | 0.4289 | 0.3363 | 1.2954 | 0.7080 | 1.9327 | 1.0333 | 0.6906 | 0.9255 | 1.1525 | 0.2828 | 0.8798 | 0.2865 | 0.9649 | 0.9753 |
| Niraparib | 100 | 0.1974 | 0.1884 | 0.4158 | 0.2477 | 0.9192 | 0.8552 | 0.2385 | 0.2411 | 1.4117 | 0.4233 | 1.2064 | 0.5839 | 0.3393 | 0.7041 | 0.9480 | 0.1391 | 0.6585 | 0.1555 | 0.7796 | 1.0679 |
| Niraparib | 200 | 0.0751 | 0.0840 | 0.1375 | 0.0943 | 0.7293 | 0.7854 | 0.0692 | 0.0872 | 1.4453 | 0.3095 | 1.4983 | 0.4639 | 0.2045 | 0.7066 | 0.9405 | 0.0248 | 0.4755 | 0.0376 | 1.0641 | 1.2341 |
| Niraparib | 500 | 0.0029 | 0.0062 | 0.0115 | 0.0027 | 1.2410 | 0.2987 | 0.0063 | 0.0587 | 1.3166 | 0.0470 | 1.1802 | 0.2123 | 0.0838 | 0.2631 | 0.5339 | 0.0074 | 0.1194 | 0.0015 | 0.9279 | 1.5952 |
| Olaparib | 0 | 1 | 1 | 1 | 1 | 1 | 1 | 1 | 1 | 1 | 1 | 1 | 1 | 1 | 1 | 1 | 1 | 1 | 1 | 1 | 1 |
| Olaparib | 10 | 0.6496 | 0.6366 | 0.6750 | 0.6099 | 1.1489 | 1.1683 | 0.6381 | 0.4687 | 1.0818 | 0.9234 | 1.9359 | 0.8220 | 0.8352 | 0.9995 | 1.1529 | 0.5473 | 0.7547 | 0.4833 | 1.2073 | 0.9797 |
| Olaparib | 20 | 0.4122 | 0.4443 | 0.6755 | 0.4236 | 1.0441 | 1.0608 | 0.3746 | 0.5885 | 1.2734 | 0.8282 | 1.7472 | 0.9717 | 0.5737 | 0.9698 | 1.3764 | 0.3715 | 0.8972 | 0.3487 | 1.0896 | 1.2777 |
| Olaparib | 50 | 0.2492 | 0.2183 | 0.4525 | 0.1936 | 1.4604 | 1.0342 | 0.1994 | 0.4126 | 1.6701 | 0.4604 | 2.4188 | 0.6889 | 0.4467 | 1.0650 | 1.2786 | 0.2449 | 0.8435 | 0.2418 | 1.5458 | 1.4536 |
| Olaparib | 100 | 0.1303 | 0.1767 | 0.4679 | 0.1010 | 1.1199 | 0.7655 | 0.1900 | 0.1609 | 1.2887 | 0.4080 | 1.8183 | 0.5971 | 0.2516 | 0.6986 | 1.0147 | 0.1409 | 0.6699 | 0.1034 | 0.8783 | 1.0514 |
| Olaparib | 200 | 0.0912 | 0.0999 | 0.3790 | 0.0651 | 1.1357 | 0.7519 | 0.0990 | 0.1898 | 1.4077 | 0.3268 | 1.9180 | 0.6459 | 0.2216 | 0.7209 | 0.9981 | 0.0502 | 0.5673 | 0.0533 | 1.1918 | 1.1059 |
| Olaparib | 500 | 0.0223 | 0.0161 | 0.0604 | 0.0118 | 0.7982 | 0.4704 | 0.0053 | 0.0671 | 1.4608 | 0.0559 | 1.1349 | 0.3532 | 0.1048 | 0.4651 | 0.5970 | 0.0242 | 0.3021 | 0.0113 | 1.2007 | 1.0412 |
| Olaparib | 1000 | 0.0128 | 0.0127 | 0.0441 | 0.0070 | 1.6850 | 0.5390 | 0.0289 | 0.0904 | 1.9880 | 0.0635 | 1.8515 | 0.3164 | 0.0563 | 0.4788 | 0.8692 | 0.0057 | 0.3135 | 0.0148 | 1.7538 | 2.2598 |
| Olaparib | 2000 | 0.0059 | 0.0046 | 0.0080 | 0.0029 | 1.3767 | 0.3040 | 0.0059 | 0.0044 | 2.0709 | 0.0072 | 1.2572 | 0.3454 | 0.0291 | 0.2881 | 0.6546 | 0.0065 | 0.1450 | 0.0030 | 1.7340 | 2.0035 |
| Rucaparib | 0 | 1 | 1 | 1 | 1 | 1 | 1 | 1 | 1 | 1 | 1 | 1 | 1 | 1 | 1 | 1 | 1 | 1 | 1 | 1 | 1 |
| Rucaparib | 5 | 0.4600 | 0.5208 | 0.6877 | 0.6809 | 0.7158 | 0.9567 | 0.6860 | 0.5473 | 1.1242 | 0.9402 | 1.7160 | 0.9152 | 0.6724 | 0.9878 | 1.0377 | 0.5803 | 0.8127 | 0.5783 | 1.0614 | 1.2069 |
| Rucaparib | 10 | 0.4488 | 0.4688 | 0.6143 | 0.4459 | 0.8248 | 1.0468 | 0.4373 | 0.5182 | 1.0811 | 0.6820 | 1.4601 | 0.7022 | 0.5203 | 0.8189 | 0.8741 | 0.4337 | 0.7855 | 0.4334 | 0.8897 | 0.9246 |
| Rucaparib | 20 | 0.3207 | 0.2741 | 0.4934 | 0.2807 | 0.7117 | 0.8040 | 0.5879 | 0.4205 | 0.9619 | 0.4701 | 1.2993 | 0.6544 | 0.4803 | 0.8088 | 0.9846 | 0.2527 | 0.7850 | 0.5123 | 0.9341 | 1.2726 |
| Rucaparib | 50 | 0.2320 | 0.2416 | 0.3046 | 0.1762 | 1.1573 | 0.9243 | 0.3000 | 0.2967 | 1.1745 | 0.5545 | 2.0743 | 0.7283 | 0.4785 | 1.0354 | 1.2218 | 0.1965 | 0.6908 | 0.1795 | 0.9891 | 1.5299 |
| Rucaparib | 100 | 0.1210 | 0.1465 | 0.2958 | 0.1336 | 0.9316 | 0.8578 | 0.1532 | 0.2707 | 1.5262 | 0.3581 | 2.0728 | 0.5958 | 0.3065 | 0.7901 | 1.3793 | 0.0881 | 0.6877 | 0.0963 | 1.0286 | 1.1420 |
| Rucaparib | 200 | 0.0594 | 0.0732 | 0.1593 | 0.0643 | 1.2617 | 0.7913 | 0.0311 | 0.1915 | 1.9317 | 0.1818 | 1.8031 | 0.5953 | 0.2415 | 0.7846 | 1.0739 | 0.0571 | 0.5789 | 0.0513 | 1.4297 | 1.2478 |
| Rucaparib | 500 | 0.0035 | 0.0149 | 0.0294 | 0.0079 | 1.0872 | 0.3914 | 0.0046 | 0.0475 | 1.3943 | 0.0678 | 1.0475 | 0.2613 | 0.0701 | 0.4666 | 0.8099 | 0.0091 | 0.2130 | 0.0179 | 0.8327 | 1.0782 |
| Rucaparib | 1000 | 0.0054 | 0.0359 | 0.0226 | 0.0060 | 1.3889 | 0.4971 | 0.0065 | 0.0182 | 2.0579 | 0.0467 | 1.4433 | 0.2336 | 0.0473 | 0.4416 | 0.7687 | 0.0080 | 0.2049 | 0.0050 | 1.8781 | 1.8072 |
| Rucaparib | 2000 | 0.0017 | 0.0035 | 0.0031 | 0.0023 | 1.1905 | 0.3480 | 0.0033 | 0.0083 | 2.0316 | 0.0076 | 1.0641 | 0.2301 | 0.0399 | 0.2079 | 0.6807 | 0.0048 | 0.1203 | 0.0033 | 1.2563 | 1.4452 |

| # | | 101 | 102 | 103 | 104 | 105 | 106 | 107 | 108 |
|---|---|---|---|---|---|---|---|---|---|
| Drug | Concentration (nmol/L) | N3124I | I3183V | N3187K | S3291C | P3292L | S3319F | R3385H | T3387A |
| CBDCA | 0 | 1 | 1 | 1 | 1 | 1 | 1 | 1 | 1 |
| CBDCA | 20 | 0.9611 | 1.1591 | 0.9103 | 1.3518 | 1.0378 | 1.0055 | 1.2002 | 1.3580 |
| CBDCA | 50 | 1.3625 | 1.2235 | 0.9375 | 1.5334 | 1.0699 | 1.3146 | 1.2615 | 1.2128 |
| CBDCA | 100 | 0.8901 | 0.9638 | 1.1057 | 1.2464 | 1.0484 | 1.2110 | 1.1796 | 1.1807 |
| CBDCA | 200 | 0.5776 | 1.1622 | 0.9116 | 0.9385 | 1.1074 | 1.2603 | 1.0870 | 1.2375 |
| CBDCA | 500 | 0.1203 | 1.1836 | 0.7564 | 0.4942 | 0.9589 | 1.0169 | 1.1166 | 1.1568 |
| CBDCA | 1000 | 0.0218 | 1.6019 | 0.3972 | 0.1233 | 0.5137 | 0.6598 | 0.6835 | 0.7415 |
| CBDCA | 2000 | 0.0047 | 2.3878 | 0.4326 | 0.0759 | 0.3472 | 0.5501 | 0.3821 | 0.5019 |
| Niraparib | 0 | 1 | 1 | 1 | 1 | 1 | 1 | 1 | 1 |
| Niraparib | 5 | 0.8119 | 1.2934 | 1.1283 | 1.2824 | 1.2556 | 1.4708 | 1.1174 | 1.1905 |
| Niraparib | 10 | 0.9678 | 1.2156 | 0.8364 | 0.8601 | 1.1719 | 1.2062 | 1.1140 | 1.1200 |
| Niraparib | 20 | 0.6379 | 0.9949 | 0.9277 | 1.4518 | 1.0784 | 1.1000 | 1.2387 | 1.2625 |
| Niraparib | 50 | 0.4893 | 1.0668 | 0.9256 | 1.0891 | 0.9575 | 1.3374 | 1.2690 | 1.2089 |
| Niraparib | 100 | 0.3567 | 1.0058 | 0.6809 | 0.6897 | 0.6844 | 0.8803 | 1.1424 | 1.0726 |
| Niraparib | 200 | 0.1573 | 0.9154 | 0.6188 | 0.4539 | 0.7456 | 0.9727 | 0.8899 | 1.0304 |
| Niraparib | 500 | 0.0061 | 0.7089 | 0.3407 | 0.0361 | 0.3662 | 0.6924 | 0.5246 | 0.6036 |
| Olaparib | 0 | 1 | 1 | 1 | 1 | 1 | 1 | 1 | 1 |
| Olaparib | 10 | 0.7091 | 1.1582 | 0.9386 | 1.1049 | 1.2615 | 1.2418 | 1.1531 | 1.3836 |
| Olaparib | 20 | 0.7861 | 1.4563 | 1.1300 | 0.9307 | 0.9850 | 1.2676 | 1.2952 | 1.1949 |
| Olaparib | 50 | 0.2776 | 1.7974 | 0.8806 | 0.7586 | 1.0402 | 1.2952 | 1.1993 | 1.4620 |
| Olaparib | 100 | 0.2302 | 0.9889 | 0.6725 | 0.4523 | 0.9039 | 1.0563 | 0.9709 | 1.1940 |
| Olaparib | 200 | 0.2919 | 1.5412 | 0.6530 | 0.3959 | 0.8073 | 0.9416 | 1.1796 | 1.2200 |
| Olaparib | 500 | 0.0339 | 1.2935 | 0.3841 | 0.2083 | 0.5282 | 0.6986 | 0.6717 | 1.1874 |
| Olaparib | 1000 | 0.0087 | 1.9922 | 0.4549 | 0.0843 | 0.5729 | 0.7606 | 0.7495 | 1.4314 |
| Olaparib | 2000 | 0.0099 | 2.2233 | 0.1615 | 0.0249 | 0.4720 | 0.3880 | 0.4921 | 1.0403 |
| Rucaparib | 0 | 1 | 1 | 1 | 1 | 1 | 1 | 1 | 1 |
| Rucaparib | 5 | 0.8205 | 0.9709 | 0.9808 | 0.9937 | 0.9358 | 0.9341 | 1.1569 | 1.1180 |
| Rucaparib | 10 | 0.5203 | 1.0385 | 0.7811 | 0.7820 | 0.8903 | 1.0708 | 0.9991 | 1.0608 |
| Rucaparib | 20 | 0.2389 | 0.9108 | 0.7175 | 0.7577 | 0.9303 | 0.6810 | 1.0052 | 1.1990 |
| Rucaparib | 50 | 0.2426 | 1.4776 | 0.7515 | 0.7230 | 0.8959 | 0.9651 | 1.1137 | 1.4301 |
| Rucaparib | 100 | 0.1892 | 1.2439 | 0.7668 | 0.5149 | 0.9729 | 1.1259 | 1.0543 | 1.4345 |
| Rucaparib | 200 | 0.1256 | 1.3133 | 0.7987 | 0.3297 | 0.7502 | 1.1707 | 0.9999 | 1.4718 |
| Rucaparib | 500 | 0.0218 | 1.2025 | 0.4819 | 0.0837 | 0.4786 | 0.5614 | 0.5805 | 0.6161 |
| Rucaparib | 1000 | 0.0105 | 1.2142 | 0.5642 | 0.1074 | 0.5198 | 0.7025 | 0.8633 | 1.0846 |
| Rucaparib | 2000 | 0.0025 | 1.4346 | 0.3433 | 0.0348 | 0.3901 | 0.5781 | 0.4063 | 0.7331 |

Values are cell viability ratio normalized to wild-type.

Supplementary Table 3: Raw data of Supplementary Figure 3.

| # | Variant | Relative viability ranking | | | |
|---|---|---|---|---|---|
| | | Olaparib | Niraparib | Rucaparib | CBDCA |
| 1 | Wild-type | 17 | 10 | 14 | 15 |
| 2 | Empty-vector | 108 | 99 | 101 | 104 |
| 3 | R18H | 59 | 46 | 49 | 78 |
| 4 | K21R | 20 | 21 | 18 | 29 |
| 5 | W31C | 96 | 95 | 105 | 107 |
| 6 | P41L | 43 | 77 | 69 | 60 |
| 7 | Y42C | 82 | 57 | 80 | 47 |
| 8 | N55S | 72 | 54 | 73 | 83 |
| 9 | P59A | 23 | 15 | 37 | 14 |
| 10 | N108S | 41 | 45 | 41 | 52 |
| 11 | Q147R | 13 | 27 | 31 | 40 |
| 12 | D156G | 27 | 20 | 33 | 12 |
| 13 | V159E | 60 | 43 | 47 | 21 |
| 14 | P168T | 48 | 37 | 40 | 38 |
| 15 | V208G | 8 | 6 | 4 | 22 |
| 16 | F266L | 40 | 32 | 35 | 39 |
| 17 | R324T | 22 | 49 | 61 | 44 |
| 18 | S326R | 29 | 58 | 42 | 61 |
| 19 | S445Y | 24 | 31 | 23 | 17 |
| 20 | E462G | 50 | 30 | 29 | 30 |
| 21 | N588D | 70 | 81 | 71 | 53 |
| 22 | H595Y | 21 | 35 | 20 | 25 |
| 23 | T598A | 12 | 11 | 9 | 23 |
| 24 | Y600C | 31 | 26 | 32 | 16 |
| 25 | G602R | 10 | 22 | 15 | 31 |
| 26 | G602V | 46 | 66 | 62 | 51 |
| 27 | E747G | 14 | 33 | 19 | 9 |
| 28 | S755C | 66 | 51 | 66 | 67 |
| 29 | N854S | 25 | 7 | 12 | 8 |
| 30 | L1019V | 78 | 67 | 55 | 59 |
| 31 | S1074C | 55 | 73 | 58 | 35 |
| 32 | I1167V | 34 | 39 | 38 | 28 |
| 33 | M1168I | 6 | 14 | 8 | 19 |
| 34 | A1170V | 28 | 18 | 17 | 11 |
| 35 | S1172L | 69 | 70 | 77 | 77 |
| 36 | N1228D | 76 | 76 | 74 | 79 |
| 37 | R1329S | 42 | 16 | 30 | 33 |
| 38 | D1420Y | 57 | 47 | 60 | 48 |
| 39 | F1524V | 71 | 75 | 72 | 74 |
| 40 | K1530N | 39 | 17 | 25 | 26 |
| 41 | V1532F | 16 | 41 | 48 | 7 |

| | | | | | |
|---|---|---|---|---|---|
| 42 | K1690N | 7 | 1 | 1 | 2 |
| 43 | E1695V | 35 | 53 | 43 | 54 |
| 44 | G1696V | 68 | 72 | 79 | 80 |
| 45 | D1699N | 49 | 42 | 26 | 55 |
| 46 | D1728N | 18 | 9 | 16 | 63 |
| 47 | D1737V | 26 | 28 | 59 | 24 |
| 48 | N1878K | 53 | 36 | 52 | 43 |
| 49 | H1918R | 75 | 79 | 81 | 65 |
| 50 | S1970L | 52 | 19 | 46 | 45 |
| 51 | E2020K | 32 | 64 | 57 | 32 |
| 52 | H2074N | 37 | 29 | 28 | 20 |
| 53 | K2075N | 45 | 50 | 36 | 66 |
| 54 | V2138D | 51 | 63 | 50 | 41 |
| 55 | H2324R | 80 | 74 | 78 | 71 |
| 56 | P2329L | 85 | 87 | 85 | 85 |
| 57 | R2336G | 4 | 3 | 7 | 3 |
| 58 | T2337A | 65 | 78 | 75 | 69 |
| 59 | K2411T | 73 | 59 | 70 | 72 |
| 60 | T2412A | 36 | 24 | 21 | 34 |
| 61 | T2412I | 58 | 44 | 39 | 56 |
| 62 | R2418G | 67 | 69 | 68 | 81 |
| 63 | N2436I | 63 | 62 | 67 | 58 |
| 64 | Y2601C | 74 | 83 | 82 | 76 |
| 65 | L2604P | 90 | 93 | 106 | 101 |
| 66 | G2609D | 91 | 102 | 108 | 96 |
| 67 | N2622S | 103 | 89 | 95 | 90 |
| 68 | H2623R | 104 | 104 | 100 | 97 |
| 69 | W2626R | 88 | 94 | 86 | 88 |
| 70 | W2626C | 95 | 103 | 94 | 89 |
| 71 | I2627V | 33 | 55 | 54 | 57 |
| 72 | P2639L | 87 | 84 | 87 | 86 |
| 73 | E2663K | 86 | 101 | 90 | 93 |
| 74 | I2664M | 81 | 68 | 63 | 73 |
| 75 | M2676T | 64 | 61 | 56 | 64 |
| 76 | L2688P | 100 | 100 | 92 | 87 |
| 77 | S2697N | 30 | 12 | 3 | 13 |
| 78 | A2717S | 19 | 48 | 27 | 50 |
| 79 | L2721H | 93 | 107 | 91 | 92 |
| 80 | T2722R | 105 | 91 | 103 | 99 |
| 81 | T2722K | 99 | 105 | 107 | 103 |
| 82 | D2723H | 101 | 97 | 96 | 108 |
| 83 | D2723E | 92 | 90 | 99 | 105 |
| 84 | G2724W | 107 | 106 | 104 | 94 |
| 85 | K2729N | 9 | 5 | 11 | 6 |
| 86 | V2747I | 61 | 60 | 51 | 62 |

| | | | | | |
|---|---|---|---|---|---|
| 87 | G2748D | 98 | 96 | 98 | 95 |
| 88 | R2842H | 102 | 85 | 88 | 98 |
| 89 | R2896C | 2 | 4 | 2 | 1 |
| 90 | V2908G | 94 | 86 | 89 | 91 |
| 91 | A2911E | 11 | 8 | 13 | 10 |
| 92 | A2911V | 56 | 71 | 64 | 75 |
| 93 | D2913H | 83 | 82 | 83 | 84 |
| 94 | I2944F | 62 | 65 | 65 | 68 |
| 95 | I3044V | 38 | 38 | 24 | 27 |
| 96 | R3052W | 97 | 92 | 93 | 106 |
| 97 | P3054S | 79 | 80 | 76 | 70 |
| 98 | D3095E | 106 | 108 | 97 | 102 |
| 99 | E3096K | 5 | 13 | 10 | 18 |
| 100 | Y3098H | 3 | 2 | 5 | 5 |
| 101 | N3124I | 89 | 98 | 102 | 100 |
| 102 | I3183V | 1 | 23 | 6 | 4 |
| 103 | N3187K | 77 | 56 | 53 | 42 |
| 104 | S3291C | 84 | 88 | 84 | 82 |
| 105 | P3292L | 47 | 52 | 45 | 49 |
| 106 | S3319F | 54 | 25 | 34 | 36 |
| 107 | R3385H | 44 | 40 | 44 | 46 |
| 108 | T3387A | 15 | 34 | 22 | 37 |

High ranking corresponds to high fold-change (neutral group).

Supplementary Table 3: Raw data of 7,344 relative viability values.

| Variant | IARC classification | Batch | Experiment | Drug | Log10(Fold-change) |
|---|---|---|---|---|---|
| G2508S | Class 4 | 2 | 1 | CBDCA | -2.807 |
| G2508S | Class 4 | 2 | 2 | CBDCA | -2.657 |
| G2508S | Class 4 | 2 | 3 | CBDCA | -2.614 |
| G2508S | Class 4 | 3 | 1 | CBDCA | -1.760 |
| G2508S | Class 4 | 3 | 2 | CBDCA | -2.567 |
| G2508S | Class 4 | 3 | 3 | CBDCA | -1.772 |
| G2508S | Class 4 | 2 | 1 | niraparib | -2.184 |
| G2508S | Class 4 | 2 | 2 | niraparib | -1.967 |
| G2508S | Class 4 | 2 | 3 | niraparib | -1.852 |
| G2508S | Class 4 | 3 | 1 | niraparib | -3.156 |
| G2508S | Class 4 | 3 | 2 | niraparib | -2.184 |
| G2508S | Class 4 | 3 | 3 | niraparib | -3.191 |
| G2508S | Class 4 | 2 | 1 | olaparib | -2.035 |
| G2508S | Class 4 | 2 | 2 | olaparib | -1.898 |
| G2508S | Class 4 | 2 | 3 | olaparib | -1.829 |
| G2508S | Class 4 | 3 | 1 | olaparib | -1.893 |
| G2508S | Class 4 | 3 | 2 | olaparib | -1.462 |
| G2508S | Class 4 | 3 | 3 | olaparib | -1.865 |
| G2508S | Class 4 | 2 | 1 | rucaparib | -1.421 |
| G2508S | Class 4 | 2 | 2 | rucaparib | -1.320 |
| G2508S | Class 4 | 2 | 3 | rucaparib | -1.270 |
| G2508S | Class 4 | 3 | 1 | rucaparib | -1.196 |
| G2508S | Class 4 | 3 | 2 | rucaparib | -0.970 |
| G2508S | Class 4 | 3 | 3 | rucaparib | -1.182 |

Supplementary Table 3: Raw data of Figure 3. class1245

| Variant | IARC Classification | Olaparib | | | | | Niraparib | | | | | Rucaparib | | | | | CBDCA | | | | |
|---|---|---|---|---|---|---|---|---|---|---|---|---|---|---|---|---|---|---|---|---|---|
| | | η | | | BF | fClass | η | | | BF | fClass | η | | | BF | fClass | η | | | BF | fClass |
| | | Mean | 95% CI | | | | Mean | 95% CI | | | | Mean | 95% CI | | | | Mean | 95% CI | | | |
| | | | LL | UL | | | | LL | UL | | | | LL | UL | | | | LL | UL | | |
| G2508S | Class 4 | -1.89 | -2.36 | -1.37 | 3.00E+00 | fClass 3 | -2.42 | -2.85 | -2.01 | 1.54E+02 | fClass 4 | -1.83 | -2.31 | -1.31 | 3.77E+00 | fClass 3 | -2.34 | -2.82 | -1.86 | 3.16E+01 | fClass 4 |
| T2607P | Class 4 | -2.12 | -2.56 | -1.65 | 1.22E+01 | fClass 3 | -2.70 | -3.14 | -2.30 | 1.54E+03 | fClass 5 | -2.05 | -2.52 | -1.59 | 1.27E+01 | fClass 3 | -2.59 | -3.08 | -2.13 | 1.52E+02 | fClass 4 |
| G2609D | Class 4 | -2.40 | -2.79 | -2.03 | 1.48E+02 | fClass 4 | -2.80 | -3.20 | -2.43 | 4.22E+03 | fClass 5 | -2.57 | -3.00 | -2.17 | 4.46E+02 | fClass 5 | -2.49 | -2.94 | -2.08 | 1.22E+02 | fClass 4 |
| W2626C | Class 5 | -2.67 | -3.08 | -2.31 | 8.30E+02 | fClass 5 | -2.97 | -3.39 | -2.60 | 1.14E+04 | fClass 5 | -2.71 | -3.14 | -2.31 | 1.15E+03 | fClass 5 | -2.78 | -3.26 | -2.36 | 4.23E+02 | fClass 5 |
| I2627F | Class 5 | -2.74 | -3.19 | -2.33 | 1.09E+03 | fClass 5 | -3.16 | -3.64 | -2.73 | 2.71E+04 | fClass 5 | -2.77 | -3.26 | -2.32 | 1.24E+03 | fClass 5 | -2.54 | -3.03 | -2.09 | 1.25E+02 | fClass 4 |
| L2647P | Class 4 | -2.63 | -3.08 | -2.21 | 5.07E+02 | fClass 5 | -2.56 | -3.01 | -2.16 | 4.98E+02 | fClass 5 | -2.64 | -3.12 | -2.20 | 5.74E+02 | fClass 5 | -2.78 | -3.31 | -2.32 | 3.47E+02 | fClass 4 |
| L2653P | Class 5 | -2.78 | -3.24 | -2.37 | 1.29E+03 | fClass 5 | -2.78 | -3.22 | -2.36 | 2.33E+03 | fClass 5 | -2.83 | -3.31 | -2.38 | 1.61E+03 | fClass 5 | -2.75 | -3.26 | -2.29 | 3.55E+02 | fClass 5 |
| R2659G | Class 5 | -2.34 | -2.77 | -1.92 | 6.31E+01 | fClass 4 | -2.65 | -3.09 | -2.23 | 8.56E+02 | fClass 5 | -2.31 | -2.77 | -1.85 | 5.91E+01 | fClass 4 | -2.48 | -2.96 | -2.02 | 9.13E+01 | fClass 4 |
| S2670L | Class 4 | -2.71 | -3.17 | -2.29 | 7.20E+02 | fClass 5 | -3.08 | -3.55 | -2.65 | 1.74E+04 | fClass 5 | -2.75 | -3.24 | -2.30 | 1.07E+03 | fClass 5 | -2.73 | -3.25 | -2.26 | 3.27E+02 | fClass 4 |
| L2688P | Class 4 | -2.54 | -2.93 | -2.18 | 3.77E+02 | fClass 5 | -2.91 | -3.33 | -2.54 | 8.29E+03 | fClass 5 | -2.47 | -2.88 | -2.09 | 2.44E+02 | fClass 4 | -2.44 | -2.87 | -2.03 | 8.97E+01 | fClass 4 |
| T2722R | Class 5 | -2.57 | -2.96 | -2.21 | 4.53E+02 | fClass 5 | -2.66 | -3.05 | -2.30 | 1.39E+03 | fClass 5 | -2.54 | -2.96 | -2.16 | 3.86E+02 | fClass 5 | -2.59 | -3.03 | -2.18 | 2.18E+02 | fClass 4 |
| D2723H | Class 5 | -2.52 | -2.52 | -2.52 | 5.06E+02 | fClass 5 | -2.52 | -2.52 | -2.52 | 7.61E+02 | fClass 5 | -2.52 | -2.52 | -2.52 | 4.90E+02 | fClass 5 | -2.52 | -2.52 | -2.52 | 2.15E+02 | fClass 4 |
| G2748D | Class 5 | -2.59 | -2.99 | -2.23 | 4.94E+02 | fClass 5 | -2.57 | -2.95 | -2.21 | 6.80E+02 | fClass 5 | -2.62 | -3.05 | -2.23 | 6.37E+02 | fClass 5 | -2.50 | -2.96 | -2.10 | 1.30E+02 | fClass 4 |
| R3052W | Class 5 | -2.69 | -3.09 | -2.31 | 8.13E+02 | fClass 5 | -2.90 | -3.31 | -2.53 | 7.64E+03 | fClass 5 | -2.69 | -3.12 | -2.29 | 9.25E+02 | fClass 5 | -2.85 | -3.34 | -2.43 | 4.49E+02 | fClass 5 |
| D3095E | Class 5 | -2.85 | -3.26 | -2.47 | 2.23E+03 | fClass 5 | -3.00 | -3.41 | -2.62 | 1.33E+04 | fClass 5 | -2.80 | -3.25 | -2.40 | 1.76E+03 | fClass 5 | -2.63 | -3.08 | -2.21 | 2.44E+02 | fClass 4 |
| N3124I | Class 5 | -2.50 | -2.88 | -2.14 | 2.57E+02 | fClass 4 | -3.17 | -3.61 | -2.79 | 3.70E+04 | fClass 5 | -2.63 | -3.06 | -2.24 | 5.96E+02 | fClass 5 | -2.61 | -3.08 | -2.20 | 2.20E+02 | fClass 4 |
| Empty-vector | Class 5 | -2.60 | -2.98 | -2.24 | 5.62E+02 | fClass 5 | -2.53 | -2.92 | -2.17 | 5.34E+02 | fClass 5 | -2.50 | -2.92 | -2.12 | 3.17E+02 | fClass 4 | -2.47 | -2.91 | -2.06 | 1.03E+02 | fClass 4 |
| K485X | Class 5 | -2.67 | -3.12 | -2.25 | 6.40E+02 | fClass 5 | -3.02 | -3.50 | -2.60 | 1.27E+04 | fClass 5 | -2.69 | -3.18 | -2.24 | 7.22E+02 | fClass 5 | -2.72 | -3.25 | -2.26 | 3.07E+02 | fClass 4 |
| L997X | Class 5 | -2.96 | -3.44 | -2.54 | 2.83E+03 | fClass 5 | -3.49 | -4.00 | -3.02 | 1.04E+05 | fClass 5 | -3.03 | -3.54 | -2.57 | 5.41E+03 | fClass 5 | -2.91 | -3.47 | -2.43 | 4.23E+02 | fClass 5 |
| Q1502X | Class 5 | -2.78 | -3.24 | -2.36 | 1.24E+03 | fClass 5 | -2.98 | -3.44 | -2.56 | 1.10E+04 | fClass 5 | -2.82 | -3.32 | -2.37 | 1.68E+03 | fClass 5 | -2.75 | -3.28 | -2.28 | 3.22E+02 | fClass 4 |
| K1984X | Class 5 | -2.99 | -3.47 | -2.55 | 2.97E+03 | fClass 5 | -2.96 | -3.42 | -2.55 | 6.83E+03 | fClass 5 | -3.06 | -3.58 | -2.59 | 6.21E+03 | fClass 5 | -2.48 | -2.96 | -2.03 | 9.00E+01 | fClass 4 |
| C2535X | Class 5 | -2.34 | -2.75 | -1.92 | 6.19E+01 | fClass 4 | -2.17 | -2.59 | -1.73 | 1.97E+01 | fClass 4 | -2.30 | -2.77 | -1.85 | 6.37E+01 | fClass 4 | -2.43 | -2.91 | -1.97 | 6.42E+01 | fClass 4 |
