## Supplementary_Figures for "High-Throughput Functional Evaluation of *BRCA2* Variants of Unknown Significance"

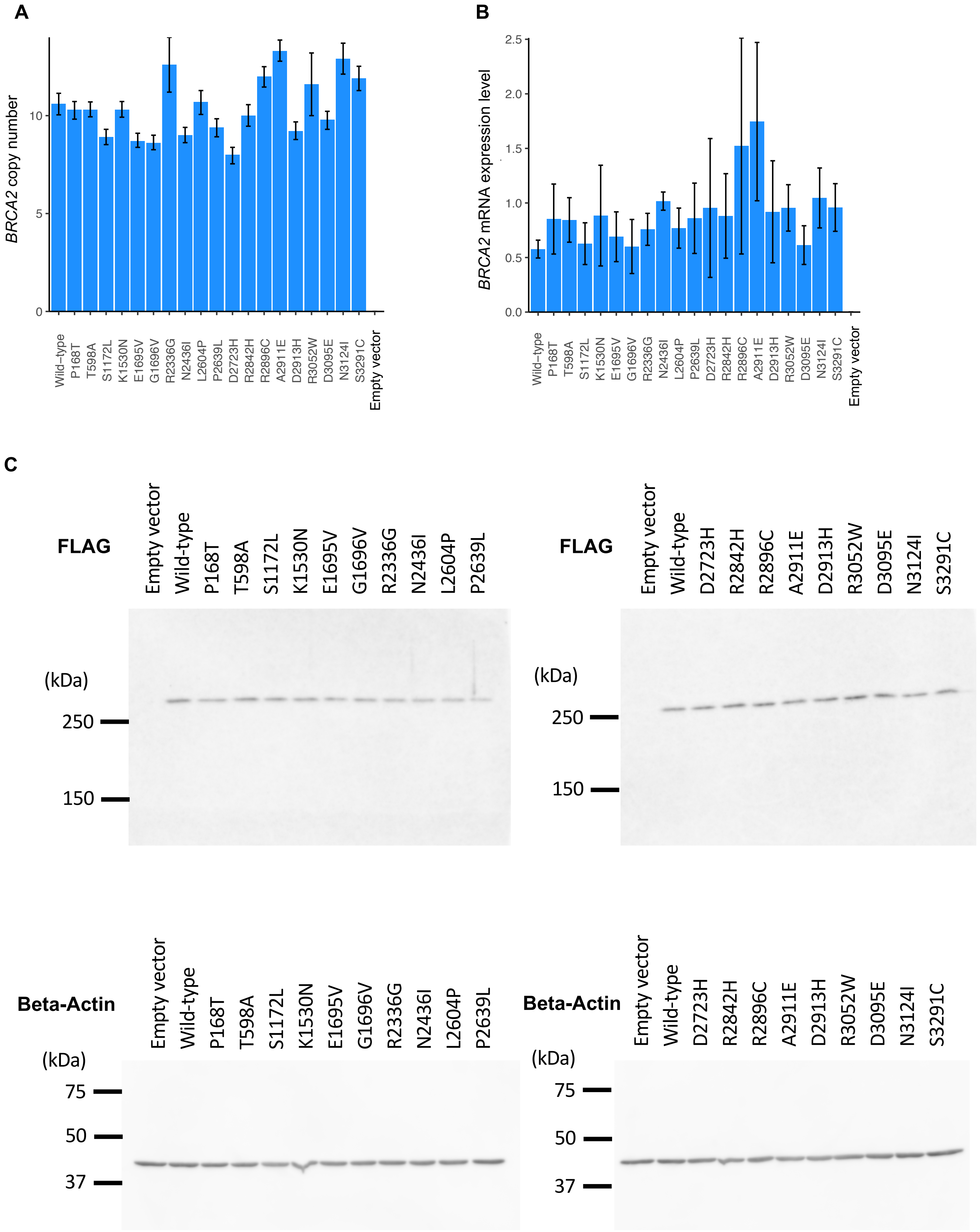


**Supplementary Figure 1**: Establishment of DLD1 BRCA2 (-/-) cells expressing BRCA2 variants. **A,** The induced *BRCA2* cDNA copy number of each variant was assessed by digital droplet PCR. Error bars, 95% confidence intervals. **B,** Real-time RT-PCR analysis of BRCA2 mRNA expression in DLD1 parental cells and DLD1 BRCA2 (-/-) cells. The BRCA2 expression levels were normalized to those of ACTB and to those in DLD1 parental cells. Experiments were performed in technical triplicate. Data are shown as the average of biological triplicates. No significant difference between variants was observed (Kruskal-Wallis rank sum test, p = 0.37). Error bars, SD (n = 3). **C**, Immunoblot analysis of DLD1 BRCA2 (−/−) cells stably expressing the FLAG-tagged BRCA2 variants. Cell lysates prepared from each cell were immunoblotted with antibodies against FLAG or vinculin. The protein expression levels of the 19 BRCA2 variants were generally equal to that of the wild-type BRCA2.

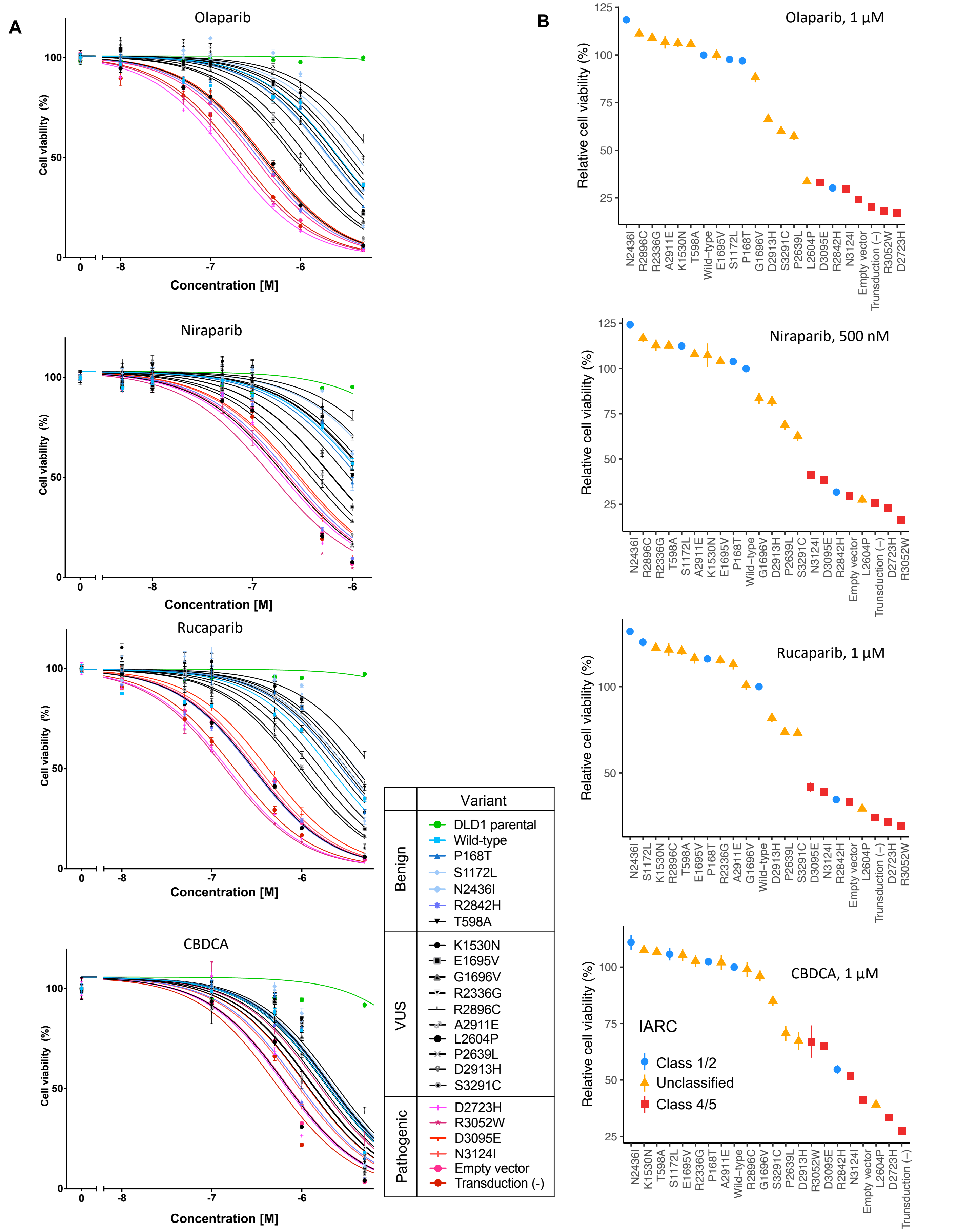


**Supplementary Figure 2:** Sensitivities of cells harboring *BRCA2* variants to PARP inhibitors and CBDCA. DLD1 parental cells, DLD1 *BRCA2* (-/-) cells harboring cDNA for one of 20 *BRCA2* variants or empty vector, or untransduced DLD1 *BRCA2* (-/-) cells were treated with the indicated concentrations of drugs for 144 hours. **A,** Cell viability was measured using PrestoBlue cell viability reagent. The sensitivity to the drugs was in concordance with the IARC classification. IARC class 1/2 (benign), blue; unclassified (VUS), black; class 4/5 (pathogenic), red. Error bars, SEM (n = 5). **B,** Relative viability of cells harboring *BRCA2* variants relative to those harboring the wild-type *BRCA2* with the indicated concentrations of drugs. The IARC classification of each variant is indicated by the color and shape of the lines and plots. Error bars, SEM (n = 5).

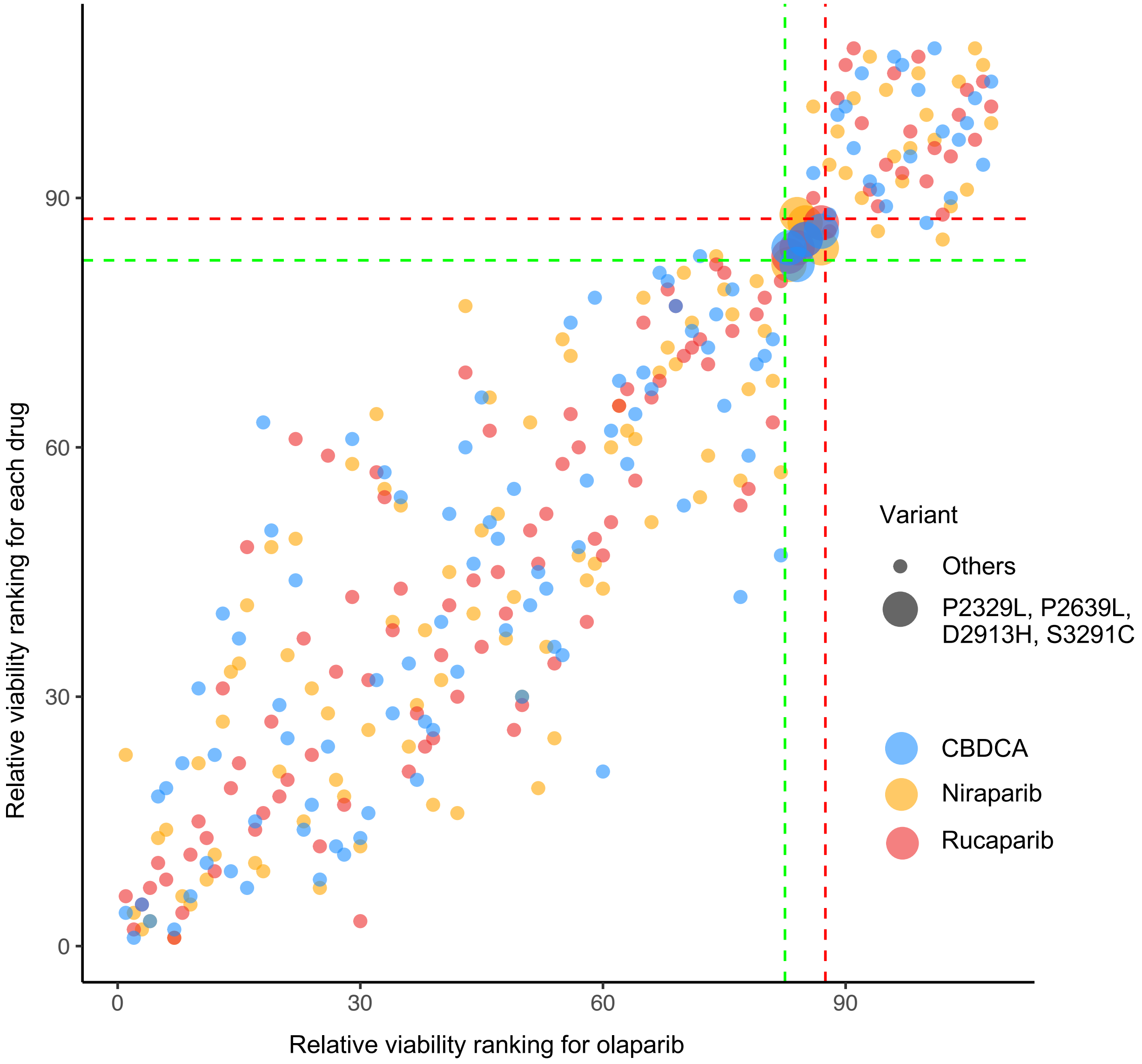


**Supplementary Figure 3:** Relative viability ranking of the 107 *BRCA2* variants and the empty vector at the highest drug concentration, as calculated by the MANO method. The relative viability rankings of each variant for niraparib, rucaparib, and CBDCA are plotted against that for olaparib. Neutral variants showing a high relative viability are located in the lower left area, whereas deleterious variants are located in the upper right area. Three components are shown: neutral (a rank of 1–82), intermediate (83–87), and deleterious (88–108). P2329L, P2639L, D2913H, and S3291C, thought to be intermediate variants, are represented by large circles.

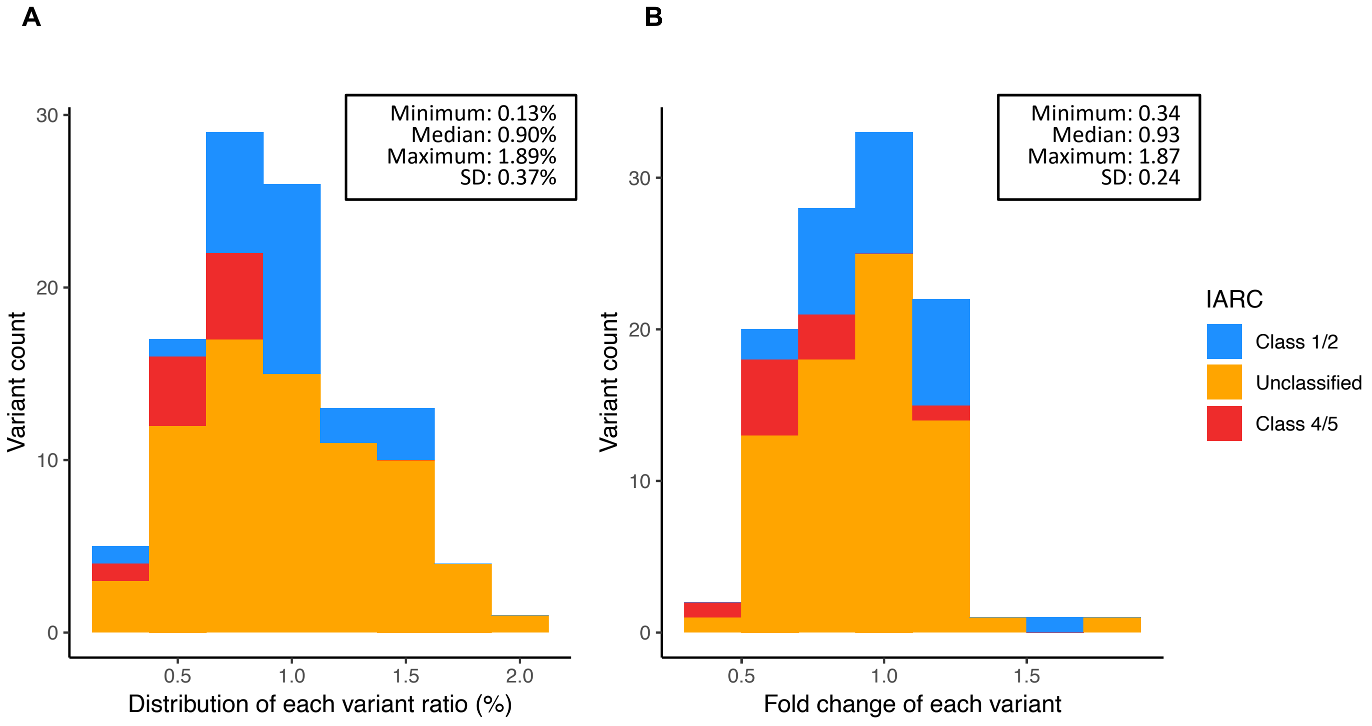


**Supplementary Figure 4:** Histograms of variant distribution in a batch of the MANO-B method**. A,** The distribution of each variant ratio at day 0 is shown. **B,** The fold change of each variant from day 0 to day 12 with DMSO treatment is shown. SD, standard deviation.

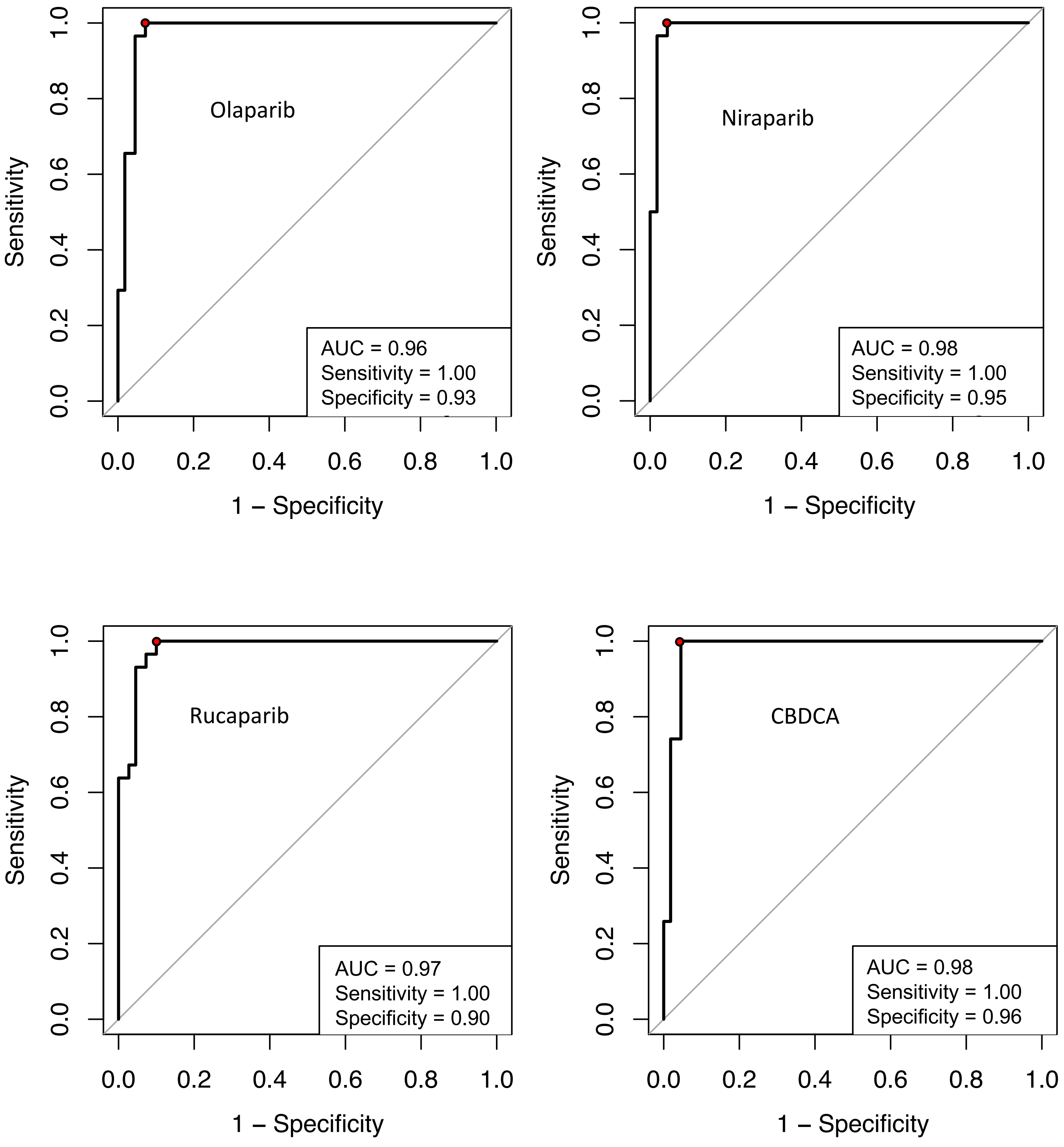


**Supplementary Figure 5:** ROC curves for the data obtained from 37 known benign and 22 known pathogenic *BRCA2* variants and the empty vector. All established benign (IARC class 1/2) and pathogenic (IARC class 4/5) variants were extracted from the 244 variants. Potential hypomorphic variants, R2784Q, R2842H, and G2908V, were classified as pathogenic at the indicated threshold, and all other variants were correctly classified.

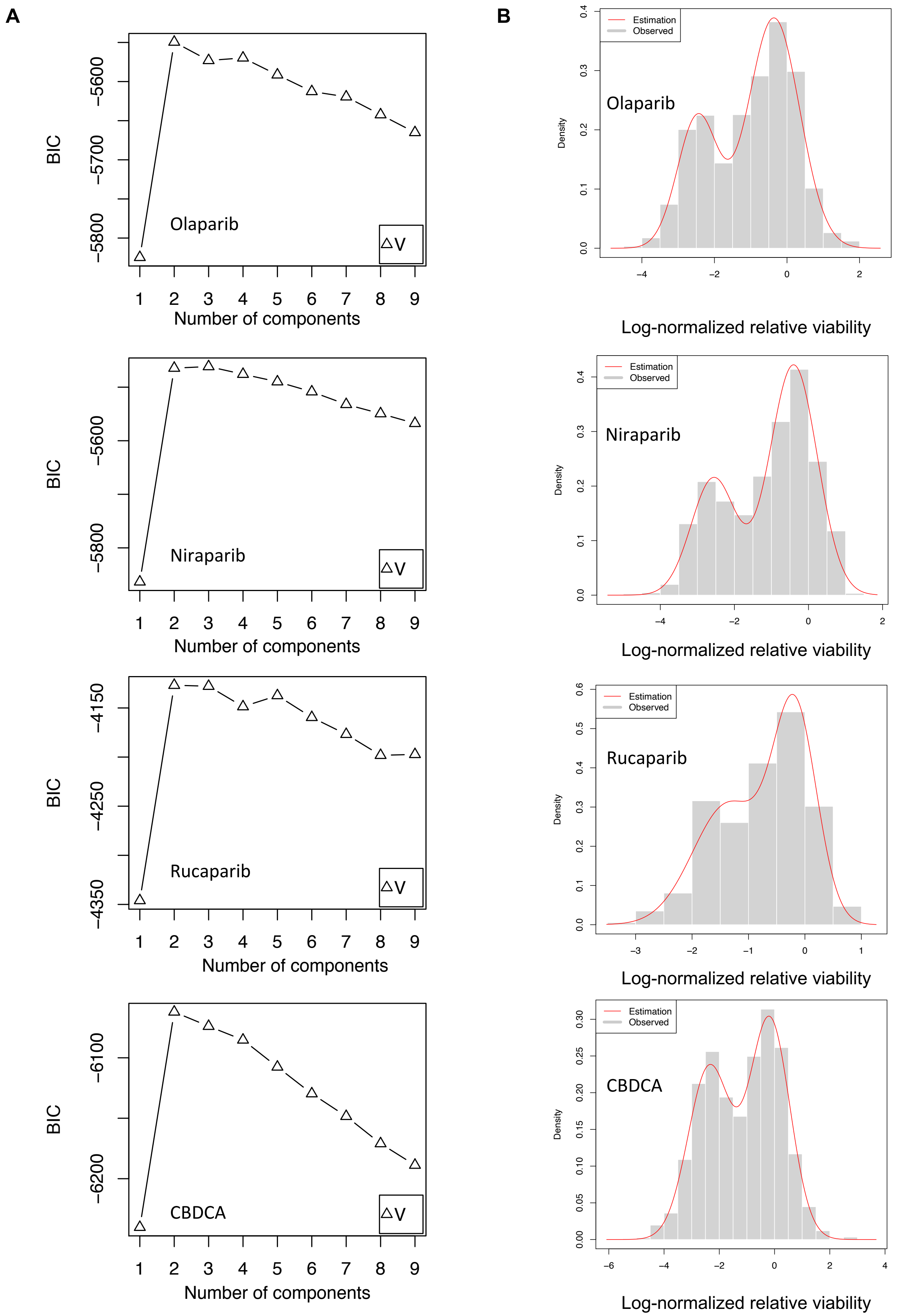


**Supplementary Figure 6:** Model estimation for the data obtained from 244 BRCA2 variants and the empty vector. **A,** Expectation–maximization algorithm to determine the appropriate model for the data. According to the Bayesian information criterion (BIC), a 2-component Gaussian mixture model was the appropriate model. **B,** The estimated 2-component Gaussian mixture distribution density curve (red line) fit the distribution of the observed log-normalized relative viability data. The left component corresponds to deleterious variants, whereas the right component corresponds to neutral variants.

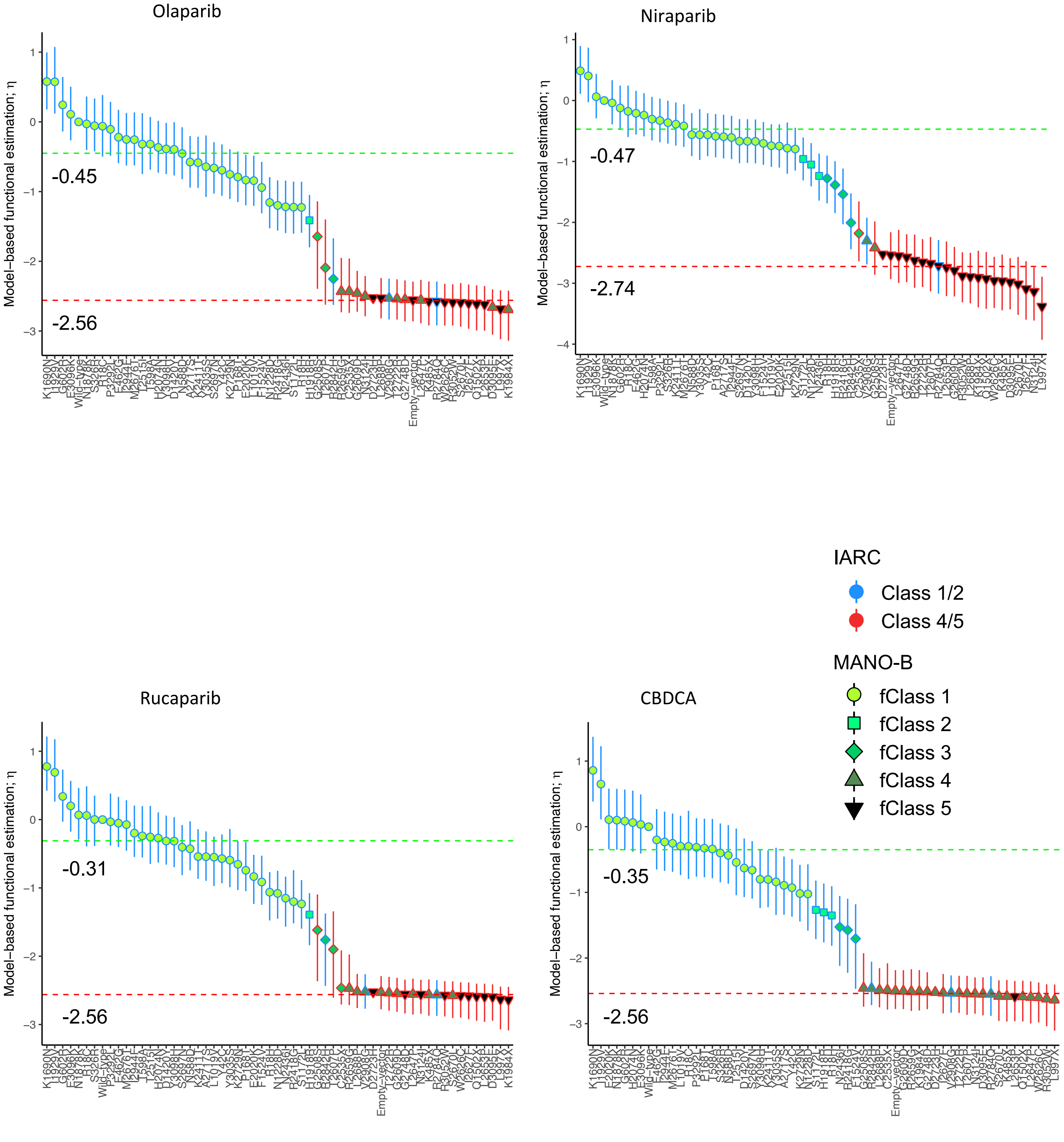


**Supplementary Figure 7:** Estimated function and classification for the training set comprising known benign and pathogenic *BRCA2* variants defined by the IARC classification. The barplots show the functional variant-specific effect η. The classification of each variant is indicated by the color and shape of the lines and plots, as shown in the legend. The $\eta_v$ values of the wild-type and the D2723H variant are anchored to $\log_{10}(1.0)$ and $\log_{10}(0.003)$, respectively. The dotted lines indicate the center of the neutral and deleterious distributions. Open error bars, 95% CIs.

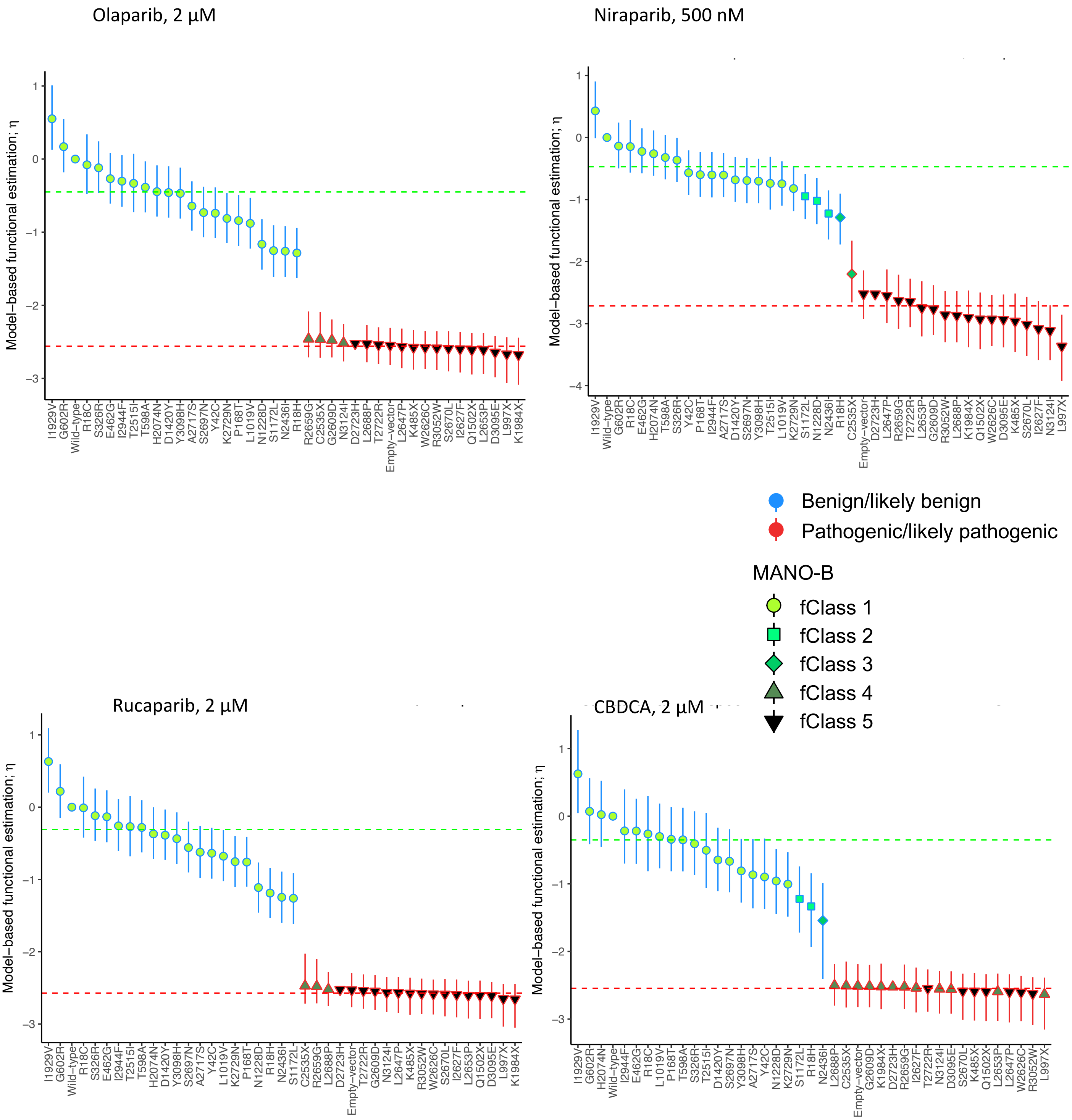


**Supplementary Figure 8:** Estimated function and classification for all the established benign and pathogenic *BRCA2* variants defined by the ACMG guidelines criteria. The barplots show the functional variant-specific effect η. The classification of each variant is indicated by the color and shape of the lines and plots, as shown in the legend. The $\eta_v$ values of the wild-type and the D2723H variant are anchored to $\log_{10}(1.0)$ and $\log_{10}(0.003)$, respectively. The dotted lines indicate the center of the neutral and deleterious distributions. Open error bars, 95% CIs.

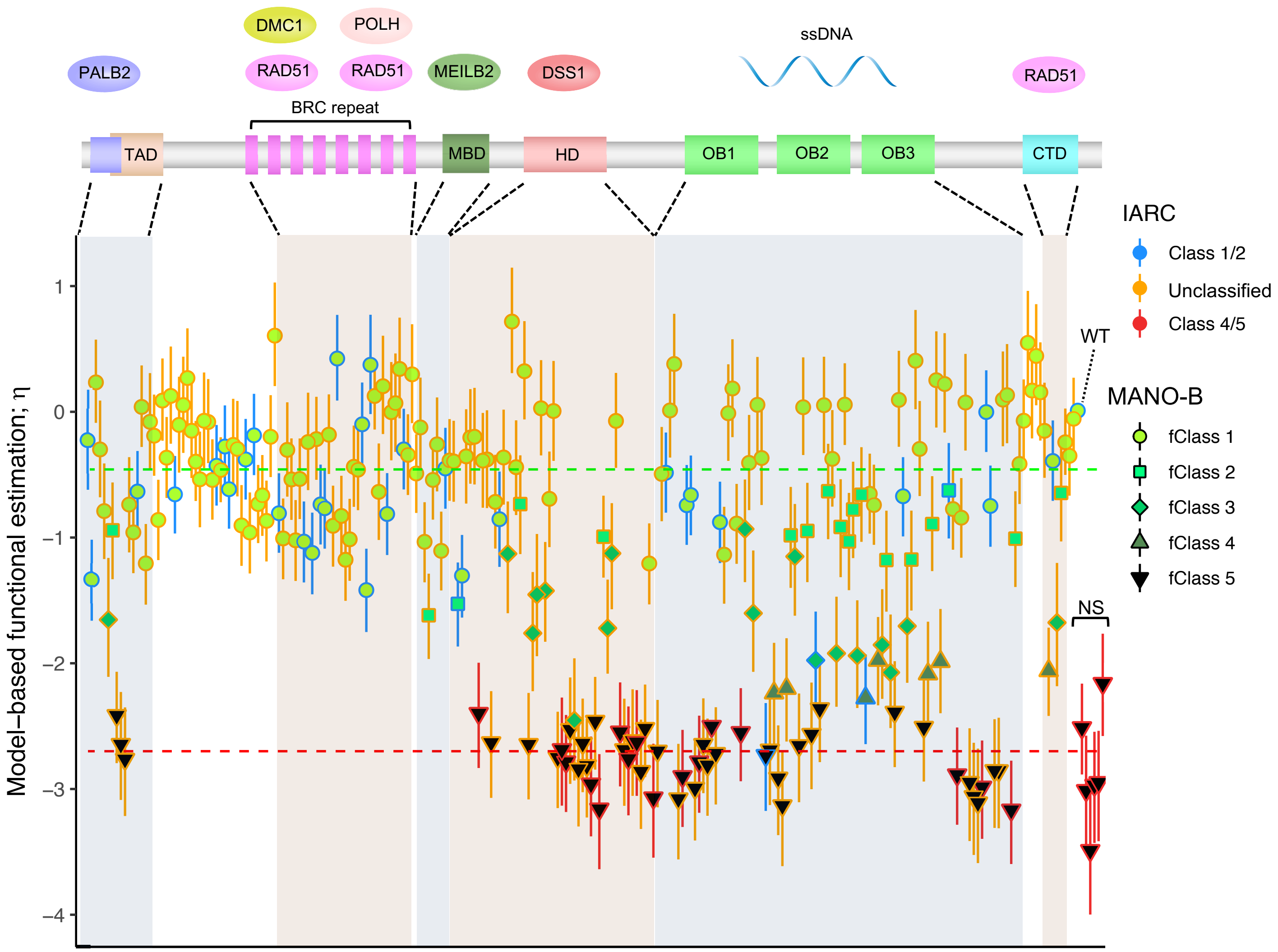


**Supplementary Figure 9:** Barplots of functional variant effects and functional diagnosis of 244 *BRCA2* variants analyzed with Align-GVGD-based prior probability. The barplots show the functional variant-specific effect η. Bayesian inference was performed with prior probability values according to the Align-GVGD algorithm. The classification of each variant is indicated by the color and shape of the lines and plots, as shown in the legend. The $\eta_v$ values of the wild-type and the D2723H variant are anchored to $\log_{10}(1.0)$ and $\log_{10}(0.003)$, respectively. The dotted lines indicate the median value of the neutral and deleterious distributions. Open error bars, 95% CIs. WT, wild-type; NS, nonsense variants.

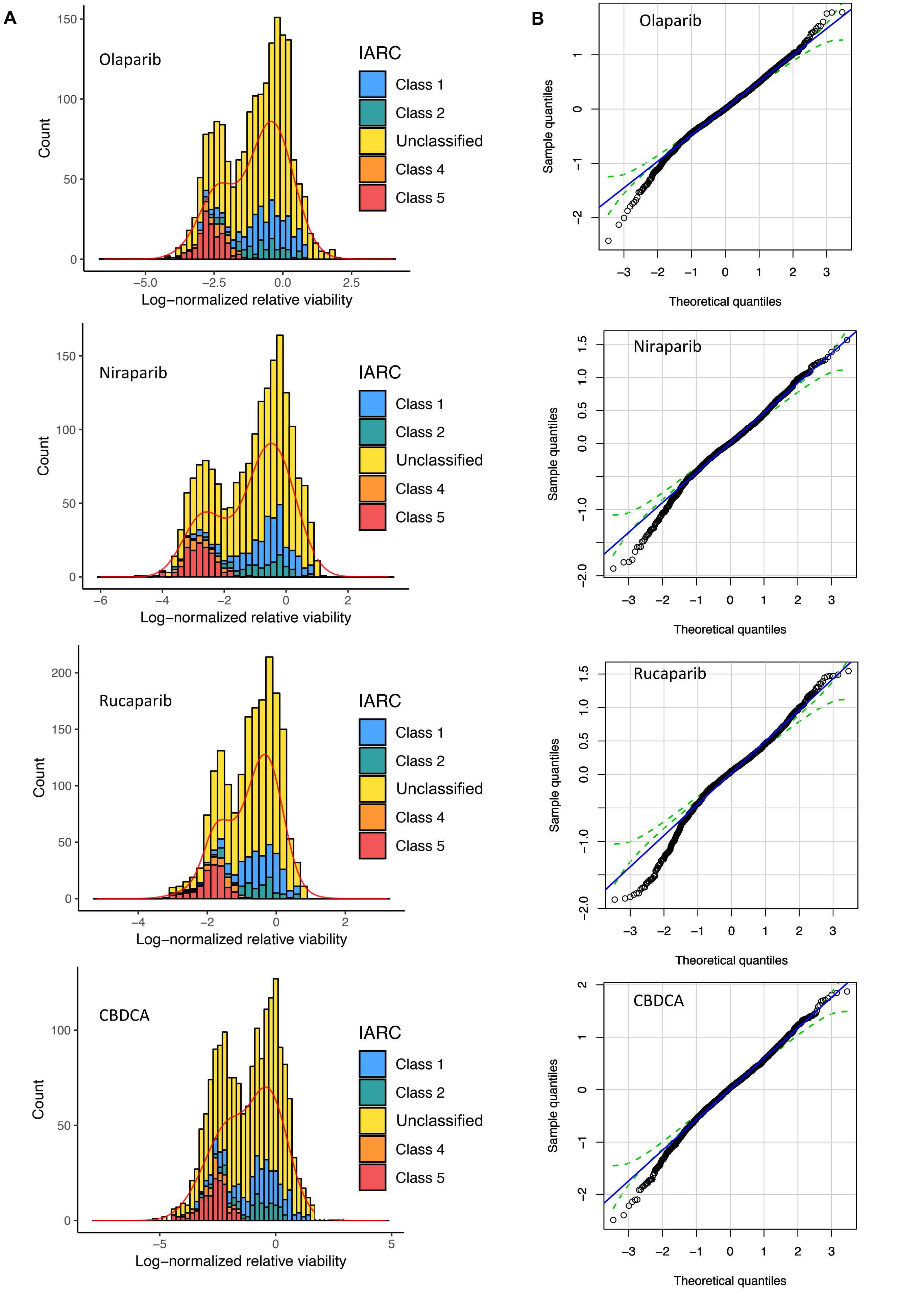


**Supplementary Figure 10:** Posterior predictive checks of the model parameters. **A,** Posterior predictive density curves of the log-normalized relative viability values for the 244 variants fit to the histogram of observed data. **B,** Normal QQ plots of the expected standardized residuals of the log-normalized relative viability.

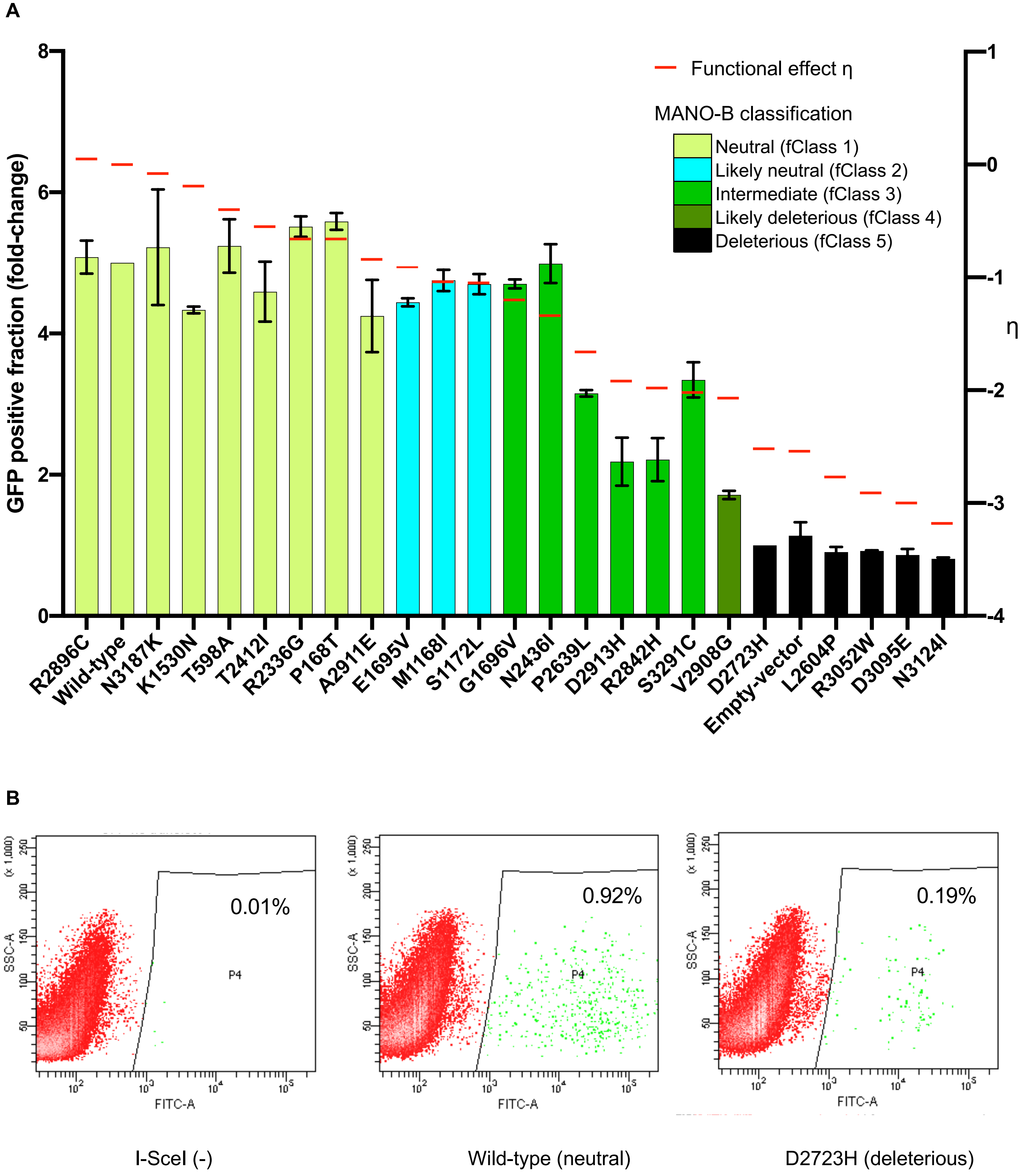


**Supplementary Figure 11:** Homology-directed repair assay of 24 *BRCA2* variants. A**,** The MANO-B classifications exhibited good correlation with the results of the HDR assay. *BRCA2* variant expression vectors and an I-SceI expression vector were cotransfected into DLD1 *BRCA2* (-/-) cells harboring the DRGFP sequence, and GFP-positive cells were counted by FACS 4 days after transfection. The GFP-positive fraction was normalized and rescaled relative to a 1:5 ratio of D2723H:wild-type. All variants were analyzed in biological duplicate and technical triplicate. The colors of the bar graphs indicate the MANO-B classification of each variant. The heights of the bar graphs indicate the GFP-positive fractions (HDR assay). Red lines show the functional effects of each variant ($\eta_v$). Error bars, s.d. (n = 2). **B,** Gating for GFP-positive cells.
